# *MaLAdapt* reveals novel targets of adaptive introgression from Neanderthals and Denisovans in worldwide human populations

**DOI:** 10.1101/2022.05.16.491756

**Authors:** Xinjun Zhang, Bernard Kim, Armaan Singh, Sriram Sankararaman, Arun Durvasula, Kirk E. Lohmueller

**Author notes:** Address correspondence to Xinjun Zhang or Kirk E. Lohmueller. Joint first authors (Xinjun Zhang, Bernard Kim). Joint senior authors (Arun Durvasula, Kirk E. Lohmueller).

## Abstract

Adaptive introgression (AI) facilitates local adaptation in a wide range of species. Many state-of-the-art methods detect AI with *ad-hoc* approaches that identify summary statistic outliers or intersect scans for positive selection with scans for introgressed genomic regions. Although widely used, these outlier-based approaches are vulnerable to a high false-negative rate as the power of different methods vary, especially for complex introgression events. Moreover, population genetic processes unrelated to AI, such as background selection or heterosis, may create similar genomic signals as AI, compromising the reliability of methods that rely on neutral null distributions. In recent years, machine learning (ML) methods have been increasingly applied to population genetic questions. Here, we present an ML-based method called *MaLAdapt* for identifying AI loci from genome-wide sequencing data. Using an Extra-Trees Classifier algorithm, our method combines information from a large number of biologically meaningful summary statistics to capture a powerful composite signature of AI across the genome. In contrast to existing methods, *MaLAdapt* is especially well-powered to detect AI with mild beneficial effects, including selection on standing archaic variation, and is robust to non-AI selection sweeps, heterosis, and demographic misspecifications. Further, *MaLAdapt* outperforms existing methods for detecting AI based on the analysis of simulated data and on a validation of empirical signals through visual impaction of haplotype patterns. We apply *MaLAdapt* to the 1000 Genomes Project human genomic data, and discover novel AI candidate regions in non-African populations, including genes that are enriched in functionally important biological pathways regulating metabolism and immune responses.

## Introduction

The discovery of archaic hominins, such as the Neanderthals in Western Eurasia and the mysterious Denisovans in Asia and Oceania^1,2,11–14,3–10^, is one of the most important scientific findings in human evolution over the last century. The high-quality ancient genomes from both Neanderthals and Denisovans^2,3,5^ further revealed that our ancestors not only overlapped with the archaic hominins in space and time during Out-of-Africa migrations, but also interbred with them, through a process known as archaic introgression. Subsequent work has shown the genomic variants from the archaic hominins played a key role in shaping the phenotypic and genotypic landscapes observed in modern humans^10,15–18^, including through adaptive introgression. Adaptive introgression refers to a process by which adaptation occurs via genetic variants that were introgressed into the modern population from the archaic population^19–21^. Currently, there is evidence of adaptive introgression in modern humans from both Neanderthals and Denisovans in worldwide populations^7,17,19,22–27^, including but not limited to the adaptation to UV radiation^16,22,23,28,29^, cold climate^29,30^, infectious diseases^11,31,32^, and high altitude environments^33–38^. Outside of modern humans, adaptive introgression also has been observed in a large range of organisms, including plants (maize, *Arabidopsis*), invertebrates (*Drosophila*, butterfly), and vertebrates(mice, fish) ^21,39–41^.

The traditional methodology to detect adaptive introgression typically relies on the “outlier approach”. Current implementations typically take on one of two flavors. The most commonly used method is to infer genome-wide signals of positive selection and introgressed ancestry separately, and then classify regions that are outliers for both attributes as targets of adaptive introgression^7,15–17,19,22–24^. Alternatively, one can use standalone summary statistics that capture signature of adaptive introgression ^1,19,42,43^. If a genomic region is an outlier to one or two of such signature statistics, it would be identified as an adaptive introgression candidate region.

Despite their wide use, both implementations of outlier approaches suffer from a series of issues that compromise power and precision. For the methods that intersect outliers from different methods, because methods to detect positive selection and archaic ancestry vary in power and have different error rates, intersecting outlier signals from these two signals can lead to a high false negative rate. This may particularly be an issue for the inference of archaic AI in modern humans, as the methods for detecting positive selection are generally more powerful at detecting recent sweep events, whereas archaic introgression occurred over more ancient time scales. The standalone statistics, on the other hand, are particularly prone to high false positive rates due to non-adaptive mechanisms compromising the null distributions for adaptive introgression ^44–46^. For example, recessive deleterious variants may accumulate privately in isolated populations. Once admixture occurs, their fitness effects become masked in hybrid individuals, leading to a heterosis effect, where introgressed ancestry increases in frequency in the absence of positive selection. Previous works^45,46^ suggest that the false positives may particularly be magnified in genomic regions with high exon density and low recombination rate, due to the elevated levels of recessive deleterious mutations leading to heterosis effects in such regions upon introgression.

In addition to challenges related to the population genetic signals of AI, genome-wide scans for selection face several statistical challenges as well. One major challenge with developing genome-wide inference tools is that the genomic regions containing the signature of interest typically represent a small proportion of the genome, compared to the proportion of genomic regions not containing the signatures. Therefore, the highly imbalanced ratio of a few true positives in a background of true negatives can easily lead to a high false discovery rate due to multiple testing^47,48^, even if a method has high power and a nominally low false-positive rate. In addition, genome-wide inference methods to detect selection often have low power due to the presence of various confounding factors, combined with the fact that most of the signatures are mild and hard to be distinguished from the genomic background.

With the rapid emergence of genomic data, machine learning (ML) and deep learning-based methods have recently been increasingly applied to the study of population genomics^49^. Compared to traditional model-based methods, ML algorithms show great promise at overcoming the restrictions of traditional statistical methods. Specifically, ML methods can have high power to detect mild signals, high precision at distinguishing confounding mechanisms, and easier implementation of realistic, complex models. In population genetics studies, recent applications of ML include the inference of selective sweeps^49–52^, archaic ancestry^22,53,54^, population demographic models^55,56^ and recombination rates^57,58^. For the detection of adaptive introgression, however, the application of ML is still in its infancy. So far, only one study^59^ has presented a deep learning method called *genomatnn*. This method is trained using genomic haplotype images, which shows high accuracy, but is computationally expensive. Furthermore, a key challenge for ML and deep learning methods is that the underlying model is unknown, therefore the deterministic mechanism for the trained model remains a black box. Here we address this issue by using biologically meaningful features in the model, and use decision tree-based algorithm so that the importance of all features in making predictions can be retrieved.

Here, we present *MaLAdapt*, a novel ML-based method for detecting genome-wide adaptive introgression in modern humans. *MaLAdapt* is trained using the pattern of functional elements in the human genome^60,61^, and modern Eurasian demographic history including single pulse of archaic introgression^2,62^. *MaLAdapt* utilizes a decision tree-based model called ExtraTreeClassifiers (ETC)^63^ as its main algorithm, and shows high power and high precision at detecting adaptive introgression signals at 50kb-resolution across the whole genome. *MaLAdapt* infers AI signature through a large composite of biologically meaningful population genetic statistics, which addresses a key challenge that it is hard to get mechanistic insights from ML/deep learning predictions. *MaLAdapt* outperforms existing methods for detecting adaptive introgression, especially given highly imbalanced class ratios, and its performance is robust to demographic misspecifications and other confounding mechanisms such as recessive deleterious mutations and positive selection unrelated to introgression. By applying *MaLAdapt* to empirical human genetic variation data from the 1000 Genomes Project^64^, we discover targets of adaptive introgression candidate regions in all non-African human populations by both Neanderthals and Denisovans that were previously undetected. We additionally present a pre-trained version of *MaLAdapt* optimized for modern human applications, as well as the simulation and machine learning pipeline scripts that enable the application of *MaLAdapt* in non-human organisms with different genomic structures and demographic histories.

## Results

### Overview of *MaLAdapt*

*MaLAdapt* is a supervised Machine Learning method for detecting genome-wide Adaptive Introgression, currently optimized at detecting adaptive introgression from archaic hominins in non-African modern human populations (Figure 1). The goal of *MaLAdapt* is to predict whether an adaptive introgression has occurred in a given 50kb genomic window. Essentially, this is a binary classification problem, where each window can be classified as “*AI*” vs. “*non-AI*”. The window-length was chosen to capture the mean length of archaic introgressed haplotypes in humans (>44kb)^3^ (see Methods). The underlying machine learning model for *MaLAdapt* is a decision tree-based algorithm called the Extra-Tree Classifier (ETC)^63^, which creates a hierarchical structure of numerous randomized decision trees that each takes a subset of features computed per 50kb window. The model further implements a meta estimator that fits the joint prediction of all decision trees. *MaLAdapt* relies on the genomic sequence and knowledge of the demographic history of a donor population, a putatively non-introgressed outgroup population, and a recipient population that experienced introgression from the donor population.

**Figure 1:**
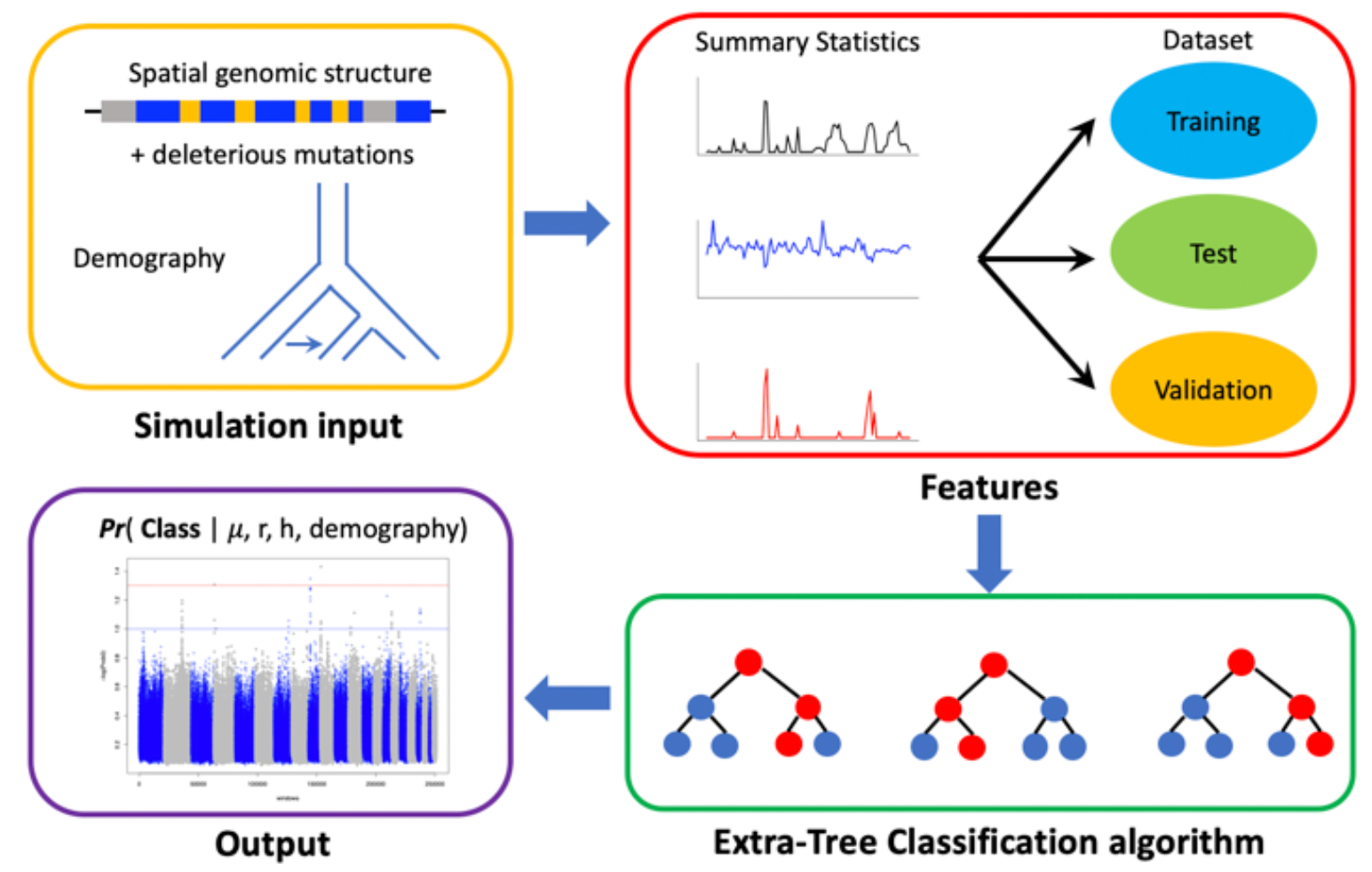
Schematic overview of *MaLAdapt* workflow. To train MaLAdapt, we simulate 1000 randomly sampled genomic segments of 5MB length with realistic genic structure, recombination rates and distribution of deleterious mutations under a modern human demography with archaic adaptive introgression (AI). We extract summary statistics in sliding 50kb-windows as features, and train a hierarchical decision tree algorithm (ERC) with data labeled with binary AI and non-AI classes. After a comprehensive model optimization, testing, and feature selection, we apply the trained model to empirical modern human genomics data to predict AI candidates.

The ETC model is trained using labeled simulation data obtained from forward-in-time simulations in SLiM^65^ of 5MB genomic segments with genic structure and recombination rates sampled from the empirical human genome, under a modern Eurasian demographic model that experienced a single pulse of archaic introgression. In each simulation, an adaptive mutation with a selection coefficient drawn from a prior distribution arises and becomes fixed in the archaic population prior to introgression, and become adaptive in the recipient Eurasian population. We vary the number of generations after the introgression (See Methods, Figure 2, and Supplementary Table 1).

**Figure 2:**
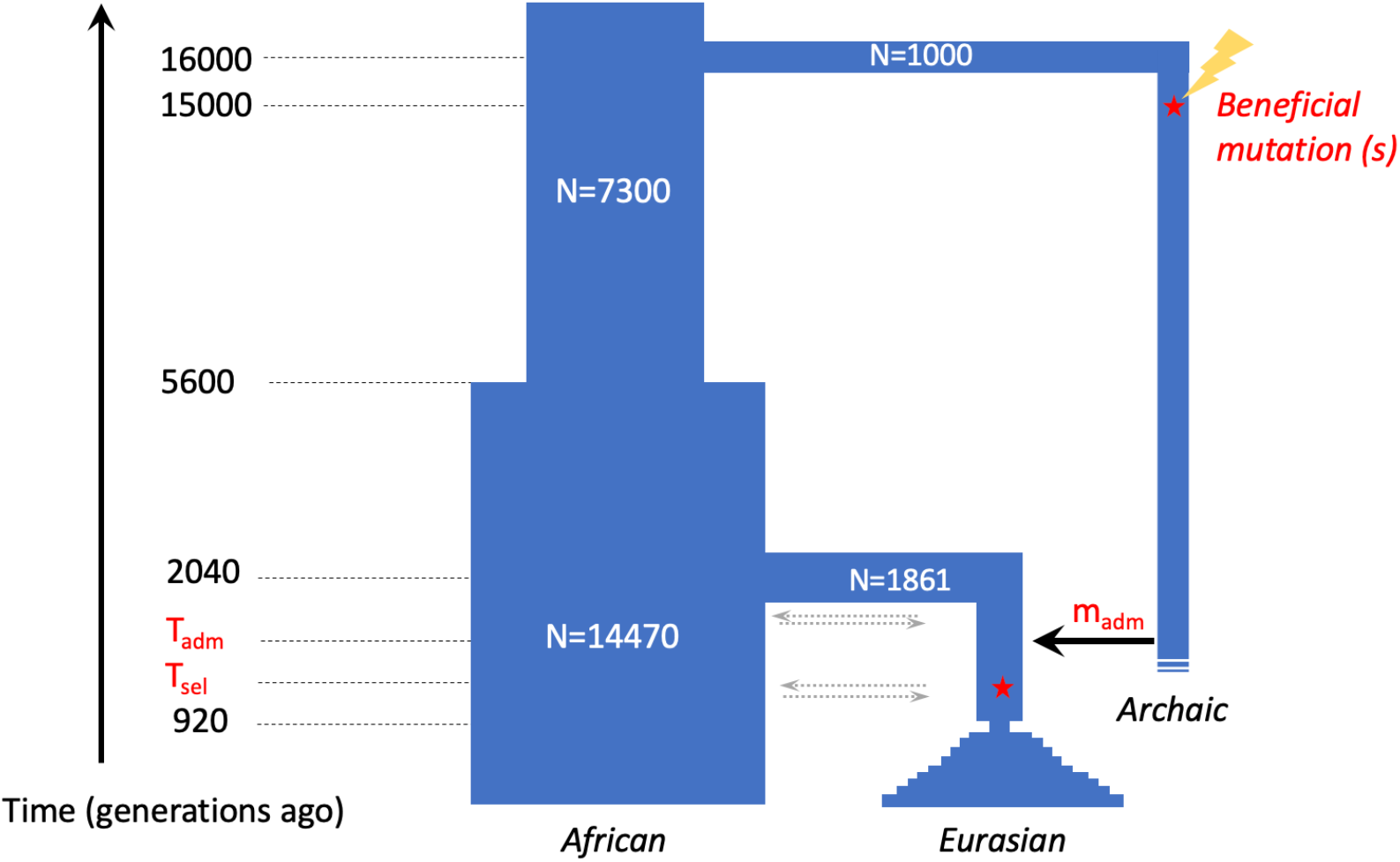
Simulation demography in *MaLAdapt*. We simulated an ancestral human population that diverged into an archaic human population and ancestral African population. The latter population subsequently split into an Eurasian population experienced two bottleneck events, representing Out-of-Africa migrations and European-Asian split, followed by an exponential growth. Sometime between the two bottleneck events, the Eurasian population experienced a single pulse of archaic introgression at a varying time and amount, which introduced a mutation that later became beneficial in the Eurasian population. See Supplementary Table 1 for the full range of simulation parameters.

Features or summary statistics are computed in 50kb sliding windows across the 5MB region. Therefore, each genomic variant is predicted five times in sliding windows. Further, given that only 5 of such 50kb-sliding windows would encompass the beneficial mutation, the ratio between “*AI*” window and “*non-AI*” window across a 5MB segment is approximately 1:100. The simulation data is further divided into training and testing datasets. Some simulations with positive selection not related to adaptive introgression were simulated under the same demography with its data included as “*non-AI*” labels in the training data. The trained model is evaluated for its performance by comparing against other ML algorithms and existing adaptive introgression signature statistics and methods. The finalized model is then used to predict adaptive introgression on all autosomes in 19 non-African populations from the 1000 Genomes project dataset^64^.

### *MaLAdapt* accurately detects adaptive introgression

We first test the accuracy of *MaLAdapt* on simulated full-5MB genomic segments under the same demography as the training data (Figure 2). Here the class ratio between *non-AI* and *AI* used for prediction, reflects the true class ratio used to simulate the test data (∼1:100). The class ratio refers to the proportion of sliding 50kb windows with and without the introgressed beneficial allele. *MaLAdapt* predicts adaptive introgression (AI vs. non-AI) in each 50kb window and returns a prediction probability. We define true or false positive as whether *MaLAdapt* predicts AI in a given 50kb window that contains the beneficial mutation. The prediction probabilities are further summarized by probability thresholds and we compute Receiver Operator Characteristic (ROC) and Precision-Recall curves (Figure 3), in which we visualize the True Positive Rate (TPR), False Positive Rate (FPR), Precision (equivalent to 1-False Discovery Rate [FDR]), and recall (equivalent to TPR) at varying thresholds. Figure 3, shows two curves for *MaLAdapt* in red and blue colors, which represent the accuracy of *MaLAdapt* at detecting adaptive introgression (*AI*) and non-adaptive introgression (*non-AI*), respectively.

**Figure 3:**
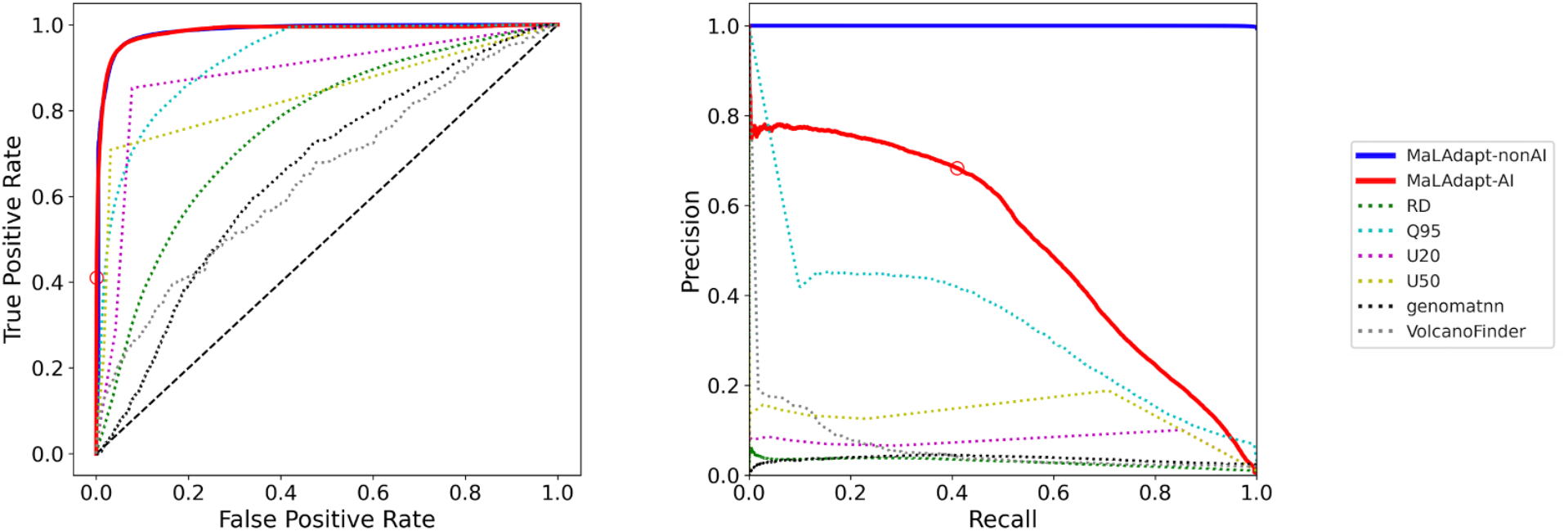
Accuracy of *MaLAdapt* and comparison to related methods. To assess the MaLAdapt performance and accuracy, we plot Receiver Operator Characteristic (ROC, left panel) and Precision-Recall (PR, right panel) curves for the prediction probabilities of MaLAdapt AI class (red solid), non-AI class (blue solid), and other AI signature statistics including RD (green dotted), Q95 (turquoise dotted), U20 (pink dotted), U50 (yellow dotted), genomatnn (black dotted), and VolcanoFinder (gray dotted) on the same testing data obtained from Figure 2 demography. The red circle corresponds to MaLAdapt AI prediction threshold of 0.9.

We compare the accuracy of *MaLAdapt* to other state-of-the-art methods for detecting adaptive introgression by applying all methods to the same testing dataset we obtained from the three-population archaic adaptive introgression model (different from the training data). *MaLAdapt* outperforms all other methods. Across all prediction probability thresholds, *MaLAdapt* has the highest power while maintaining the highest precision and the lowest false positive rate compared to all other methods under comparison, including the *RD, Q95, U20*, and *U50* summary statistics^19^, *genomatnn*^59^ - a deep learning-based method for detecting AI leveraging haplotype structure information, and *VolcanoFinder*^66^ a reference-free method for predicting AI using genomic polymorphic data (Figure 3, Supplementary Table 5). We reject the null hypothesis that the difference in AUROC between *MaLAdapt* (when predicting AI) and *Q95* - the second best-performing method – is zero with a *p*-value < 2.2e-16 via jackknife, and reject the null hypothesis that the difference in AUPR between *MaLAdapt* and *Q95* is zero with a *p*-value=1.438e-7 via jackknife^67^. Thus, we can conclude that *MaLAdapt*’s improvement of power and precision over other methods is statistically significant. We note a substantial reduction of accuracy in both *VolcanoFinder* and *genomatnn*, compared to their respective originally reports. However, there are several key differences between *genomatnn, VolcanoFinder* and *MaLAdapt* that may explain the reduced performance of this method on our simulation data, including the complexity of underlying models considered by different methods (See Discussion).

We weigh both the ROC and Precision-Recall curve to determine a prediction probability threshold for calling AI segments that maximizes the power and precision of *MaLAdapt*. We show in Figure 3 that at *Pr(AI)* = 0.9 (*i*.*e*. Pr (*non-AI*) = 0.1), the precision of *MaLAdapt* is 0.683 (FDR = 0.317), with a recall (TPR) of 0.410, and FPR at 0.001. At this threshold, *MaLAdapt* outperforms all other related methods, especially in the precision-recall curve, showing *MaLAdapt*’s outstanding ability to account for the highly imbalanced ratio between AI and non-AI classes. This is important because the class ratio is likely to be even more skewed in the human genome. Pr (*non-AI*) = 0.1 can also be justified as a multiple testing problem: in sliding 50kb windows, each locus is scanned 5 times, and a significant value for a window being AI (*i*.*e*. not being non-AI) should be the default probability threshold, which is 0.5, divided by 5.

### *MaLAdapt* is robust to misspecification of the demographic model

Next, we assessed the sensitivity of *MaLAdapt* to uncertainty and mis-specification of the demographic parameters. In the training process, most parameters related to adaptive introgression, including the time of introgression (*T*_*adm*_), the time of selection (*T*_*sel*_), selection coefficient (*s*), introgression amount (*m*), are simulated as variables drawn from uniform distributions (see Method section). Additionally, we simulated 1000 randomly sampled genomic segments of 5MB to represent the genic structure and recombination rate distribution on the empirical human genome. The rest of the demography uses a model based on the evolution of modern Eurasians^62^ with a pulse of archaic introgression^2^.

To determine the robustness of *MaLAdapt* to model misspecifications, we perturb the key adaptive introgression-related parameters one at a time, and with each alternative parameter, we simulate adaptive introgression of 5MB genomic segments (100 replicates per parameter) as new testing dataset, and apply *MaLAdapt* trained on the original model to the new testing data and evaluate its accuracy. Specifically, we ask how *MaLAdapt* performs when: 1) *T*_*sel*_ is 200 generations lower than the original lower bound of *T*_*sel*_ distribution (410 generations ago; denoted as “*T*_*sel*_*_low*”); 2) The introgression fraction (*m*) is 2-fold lower than the original lower bound (at 0.5%; denoted as “*m_low*”); 3) The introgression fraction (*m*) is 2-fold higher than the original upper bound (at 20%; denoted as “*m_high*”); 4) the selection coefficient (*s*) is 10-fold higher than the original upper bound (0.1; denoted as *s_high*); 4) the genomic segments sampled for generating testing data are different from the ones used in the training process (denoted as “*segment*”); and 5) the Eurasian population growth rate and Out-of-Africa bottleneck size are different than the training simulations (denoted as “*demo*”). We did not explore the selection coefficient (*s*) being smaller than the original lower bound (1e-4) because with such weak selection, it would be difficult to generate AI simulations without the beneficial mutation being lost in the recipient population. We also did not perturb the time of introgression (*T*_*adm*_) because the range of *T*_*adm*_ is bounded by the split time between Eurasians and ancestral Africans, as well as the split time between Europeans and Asians.

In addition to Precision, Recall (TPR), and FPR, we also computed the F1 score as an accuracy metric. F1 is defined as the weighted average between Precision and Recall (Methods). We evaluate the performance of *MaLAdapt* at the 5 alternative parameter combinations listed above by computing the Log10 fold change of each accuracy metric when comparing against values obtained from using the original testing data (Figure 4a-b). We find that *MaLAdapt* remains robust, even when most AI-related parameters are mis-specified. Especially noteworthy, the precision of AI detection was not compromised, and even increased slightly, when the selection time is low, representing selection on standing archaic variation in very recent times (<610 generations/15,000 years ago).^68–70^ Further, performance remained high when the introgression amount is low, representing a low initial frequency of archaic variants. These observations, together with the training of *MaLAdapt* accounting for extremely low strength of positive selection, show that *MaLAdapt* is particularly powerful and reliable at detecting mild, incomplete adaptive introgression sweeps. *MaLAdapt* also shows little to moderate precision loss when the demography of the recipient population changes, as well as when the testing genomic segments are different from the training segments.

**Figure 4:**
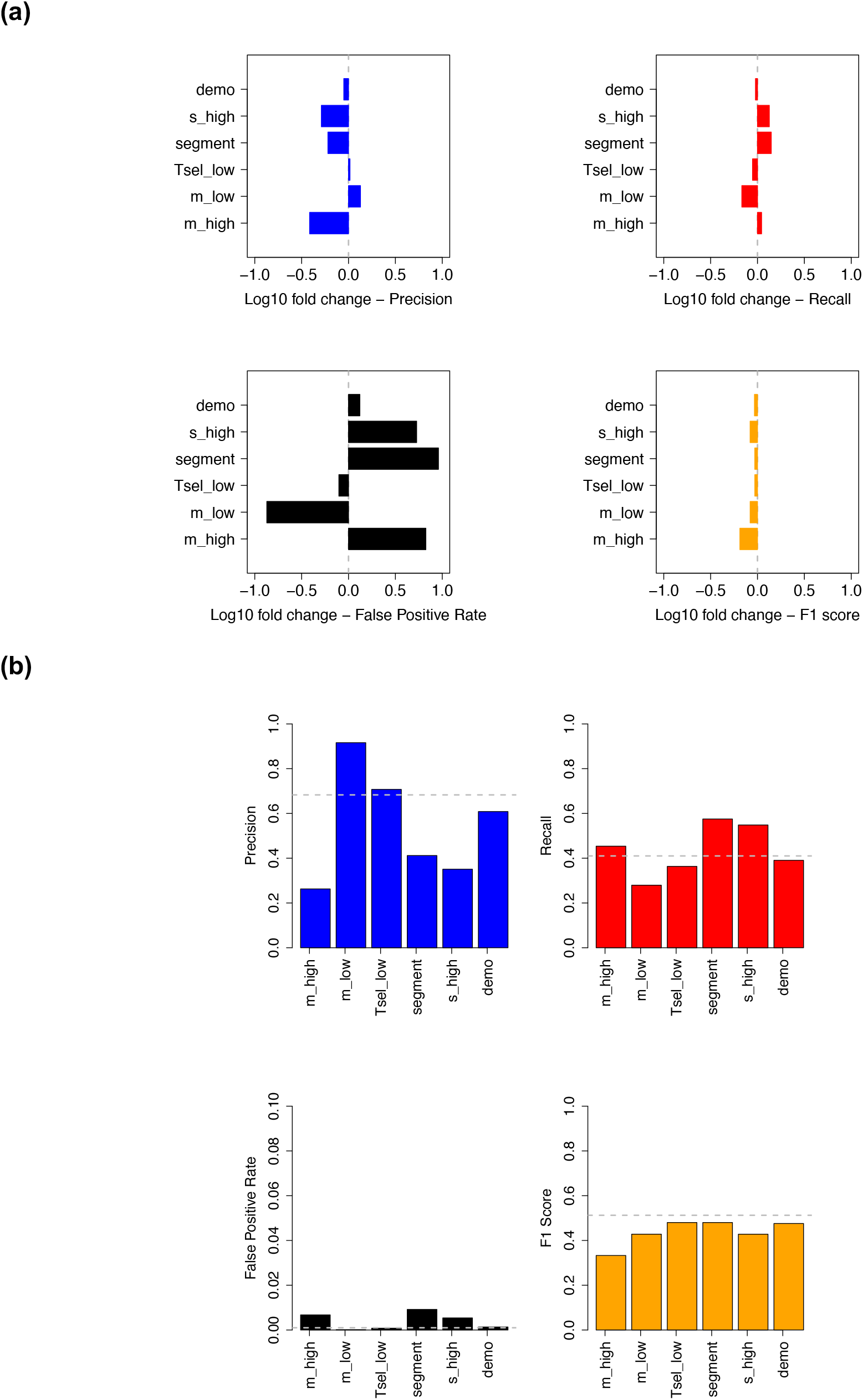
*MaLAdapt* is robust to demographic misspecification. We evaluated MaLAdapt’s robustness by applying the model to different sets of testing data with out-of-bound values of key demographic parameters compared to the training data. We compute the performance metrics (including Precision, Recall, False Positive Rate and F1 score) and compare them against the original data (gray dotted line) under each testing scenario. Panel (a) shows the log of the value difference (testing scenario minus the original), in which a longer bar indicates a higher fold change for the given metric, and the sign of the bar indicates whether the testing metric value increases (positive) or decreases (negative). Panel (b) shows the absolute value of the performance metric under each testing scenario.

There are two parameters that, when mis-specified, reduce the precision of *MaLAdapt* by more than 30%. These include large selection coefficients (*s* = 0.1, 10-fold larger than in simulations) and high introgression fraction (*m* = 20%, two-fold higher than in simulations). Strong positive selection (*s_high*) led to a loss in precision since although both FPR and TPR increased under this scenario, it inflated FPR more than it did to TPR, where a high FPR is potentially caused by falsely classifying windows nearby strong positive selection focal windows as AI. A high amount of introgression, which can be interpreted as either a significant amount of single pulse or a combination of multiple pulses, reduces precision because it increases the FPR more than it does the TPR. Promisingly, the weighted average of precision and recall, which is measured by F1, changes little with regards to any of the alternative parameters, indicating *MaLAdapt*’s robust performance at model misspecification especially with highly imbalanced class ratios.

Additionally, we assess the ability of *MaLAdapt* to distinguish adaptive introgression from positive selection unrelated to adaptive introgression. We simulated non-introgressed positive selection scenarios using 1000 genomic segments that were different from those used in the training data, with the rest of the demography and parameter distributions the same as the training data. We show in a confusion matrix (Supplementary Table 4) that *MaLAdapt* correctly assigned non-introgressed sweeps (to “*non-AI*” class) at 99.87% of the time, in contrast to “*AI*” class at 0.13%

### *MaLAdapt* reveals novel adaptive introgression targets in worldwide population from Neanderthals and Denisovans

We computed features in 50kb sliding windows across the genome using Neanderthals (Altai individual) and Denisovan (Altai Denisovan) as reference genomes respectively, and predicted AI from Neanderthals and Denisovans in 19 non-African populations from the 1000 Genomes Project^64^. In all comparisons, we use the Yorubans (YRI) as the non-introgressed outgroup. We intersected the 50kb windows predicted as AI with GENCODE database to get lists of genes overlapping with the regions, and we merged overlapping AI windows. Here we show Neanderthal AI in Europeans (CEU) as an example in the main text, and the information on Neanderthal AI in other populations as well as Denisovan AIs can be found in the Supplementary Figure 16-17 and Supplementary Table 5-6. By summarizing previously reported Neanderthal AI candidates from relevant studies, and intersecting the findings from *MaLAdapt*, we report novel Neanderthal AI candidates in all non-African populations, highlighted in the Manhattan plots (Figure 5).

**Figure 5.**
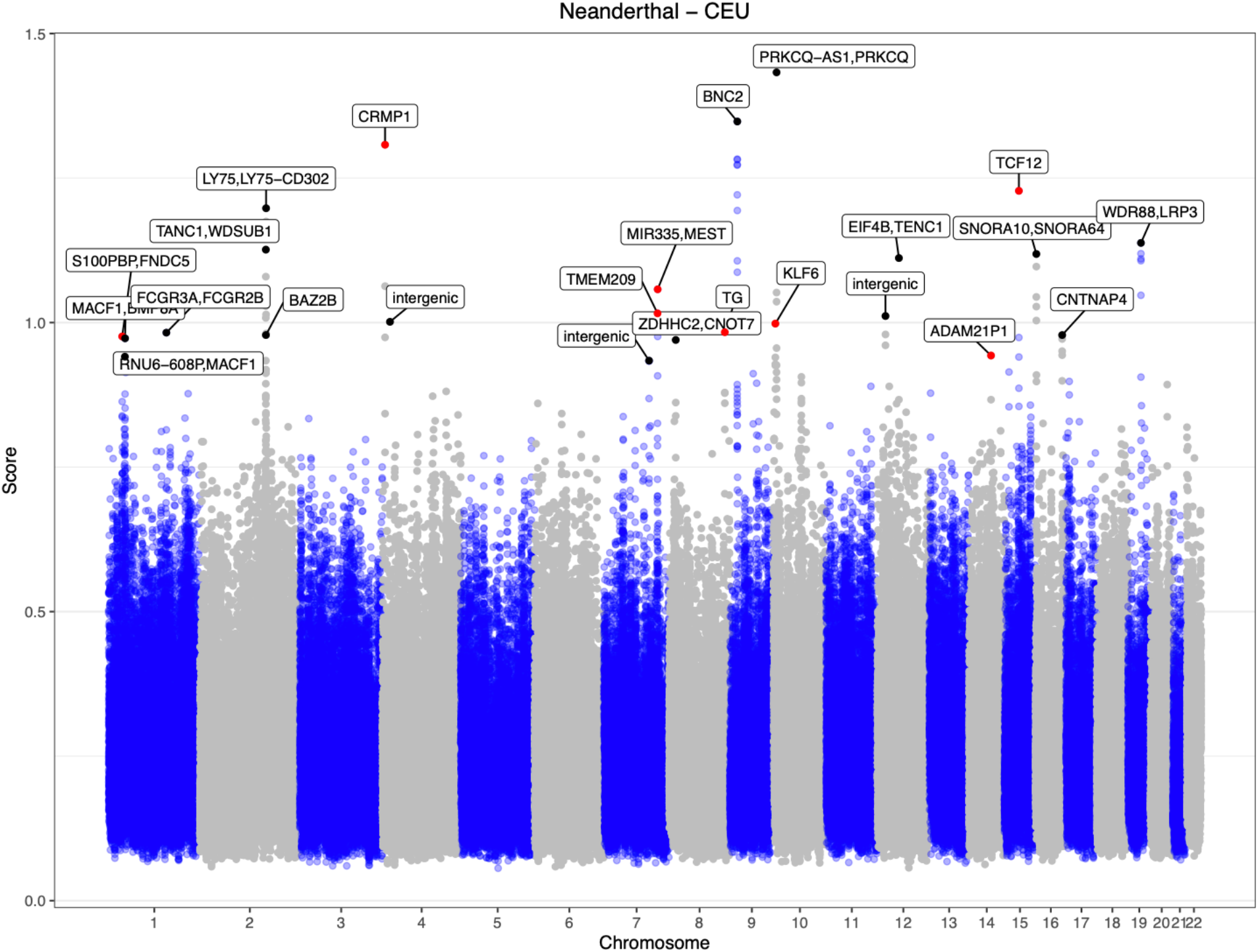
Adaptive Introgression from Neanderthals in European population (CEU) We applied MaLAdapt to predict AI in overlapping 50kb windows (step size 10kb) along the genome of non-African populations of the 1000 Genomes data. Here we show the AI prediction results of the European population (CEU), using African (YRI) as non-introgressed outgroup and Altai Neanderthal as the introgression donor. The Y-axis shows the AI prediction score, which equals the -Log10 transformed value of [1-Pr(AI)]. Each dot in the plot represents a 50kb window. The windows that did not reach the MaLAdapt AI threshold are colored in blue or gray depending on the chromosomes. The windows detected as AI are colored in black if they have been reported by previous studies before, or in red if they are novel findings from this study. The labels highlight the gene names that overlap with the AI windows.

We use a two-step process to evaluate the legitimacy of the novel AI discoveries by *MaLAdapt*. First, we summarize the canonical hits found by previous studies^10,16,17,19,23,59,66,71,72^. These are defined as genes that have been reported as a target of Neanderthal AI by more than 1 study. We ask what proportion of such canonical AI hits did *MaLAdapt* manage to discover. We show that the we found 100% of the most reported hits (those seen by at least 5 studies). On average, *MaLAdapt* detected more than 50% of other repeatedly reported Neanderthal AI hits (Table 1). For the repeatedly identified hits that *MaLAdapt* did not detect as AI, we further examined the prediction probabilities in such regions. We found that that *MaLAdapt* predicted *Pr(AI)* being no less than 0.7, suggesting that *MaLAdapt* did find evidence of AI, despite these genes did not making it over the 0.9 cutoff (Supplementary Figure 6). Next, we examined the haplotype structure of our AI candidates to visually validate the legitimacy of our hits. We show in Figure 6 and Supplementary Figure 5 that all 9 newly-discovered gene regions in CEU appear to be legitimate adaptive introgression candidates. Specifically, under AI, we expect to see a clear block of haplotypes in the introgressed population (*e*.*g*. The Europeans) that have close affinity to the archaic genome (*e*.*g*. The Neanderthals), and we do not expect such blocks of haplotypes to be present in the non-introgressed population (*e*.*g*. Yoruba)^38,73^. Note that this pattern is present in all of these candidate regions.

**Table 1:**
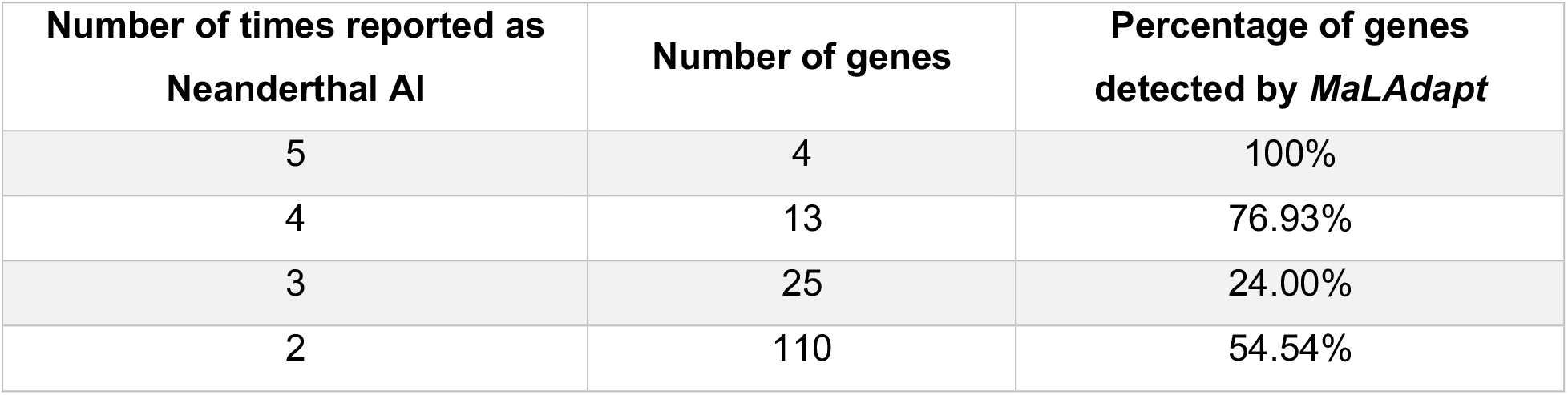
Percentage of previously reported Neanderthal AI regions detect by *MaLAdapt*. We summarize gene regions on the human genome by the number of times they have been reported by previous studies as Neanderthal AI candidates (column 1). We count the number of genes in each category (column 2), and examine the percentage of repeatedly reported AI genes that is recovered by MaLAdapt (column 3).

**Figure 6:**
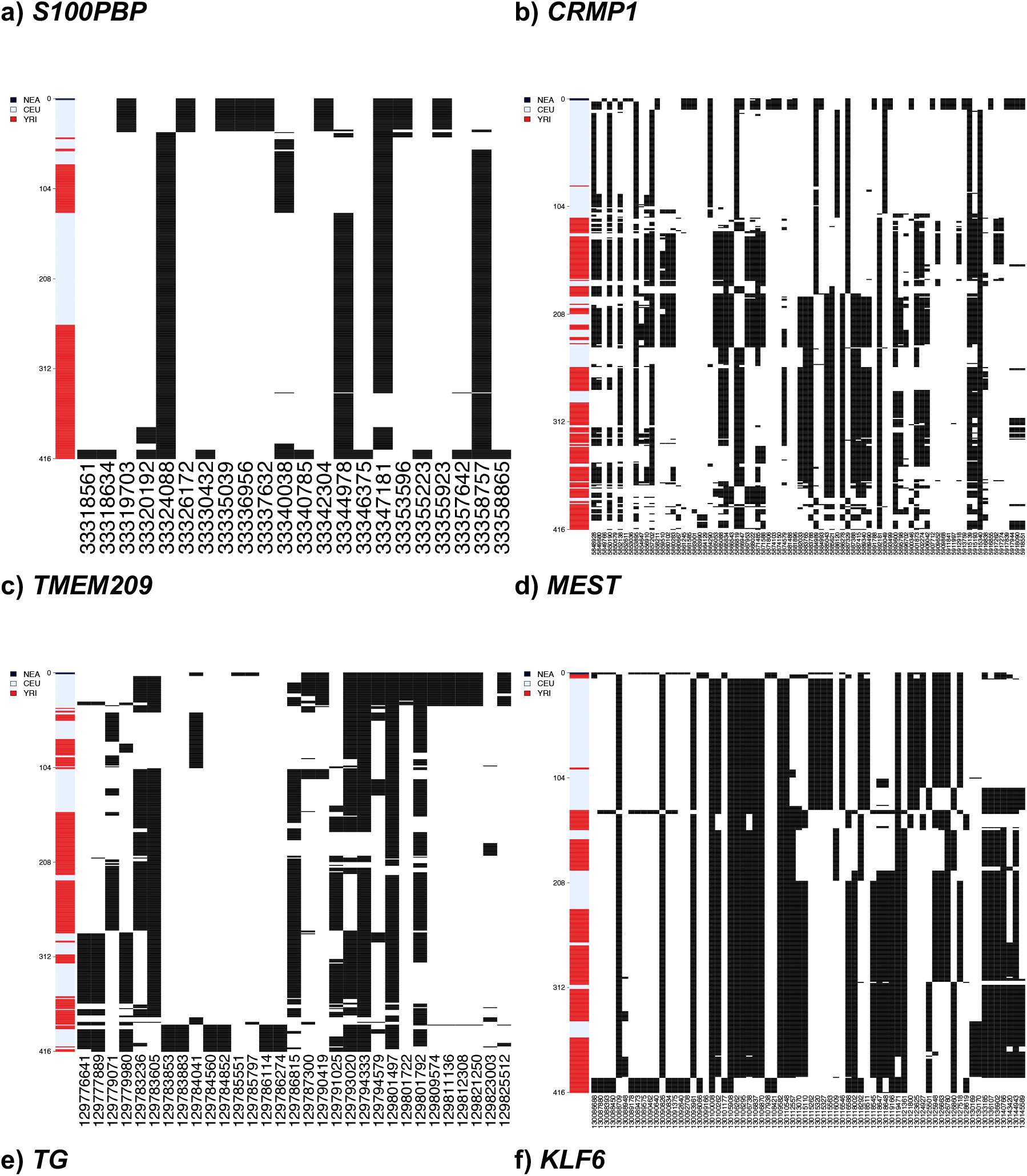
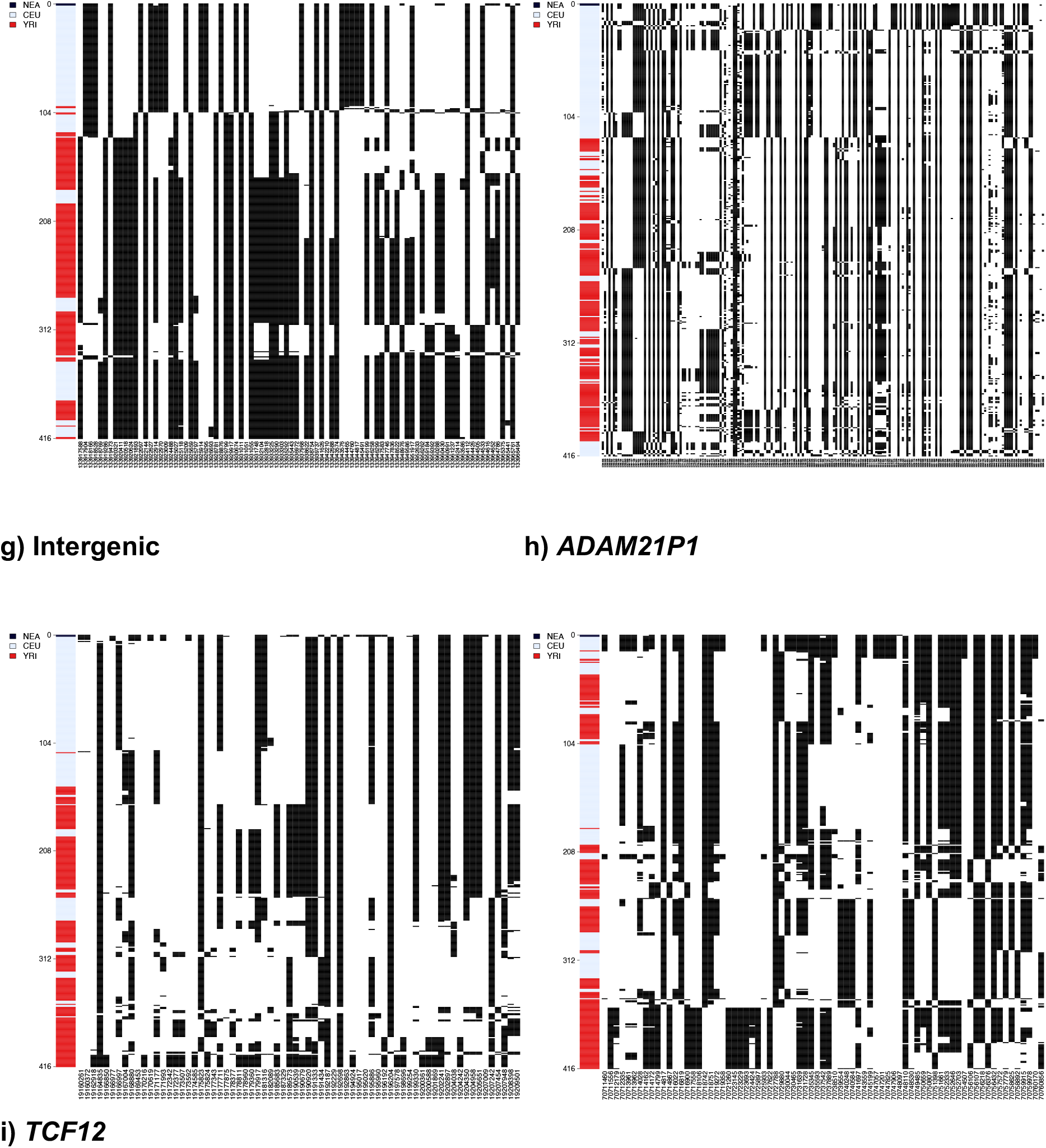
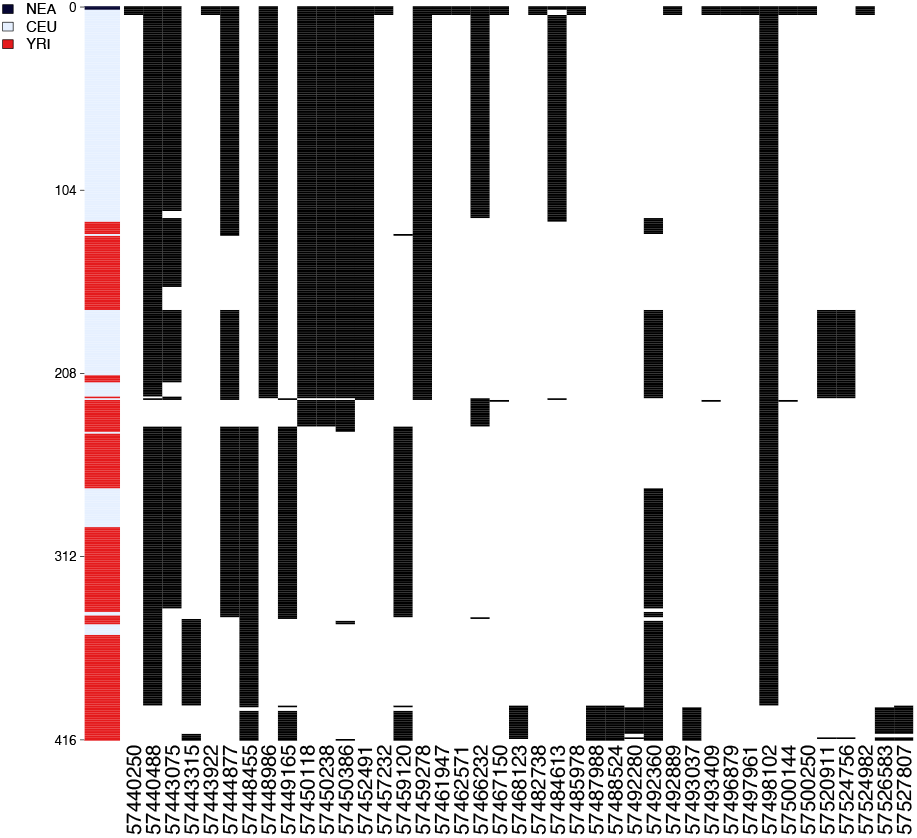
Haplotype structure of the novel Neanderthal AI candidate regions in the CEU. We plotted the haplotype structure of 9 candidate regions predicted by MaLAdapt as AI from Neanderthals in CEU. For each region, we plotted the haplotypes of Altai Neanderthal (black), CEU individuals (blue) and YRI individuals (red), and clustered and sorted the haplotypes by decreasing distance to the Neanderthal genome. In other words, rows closer to the top of the plot represent haplotypes that are more similar to that of the Neanderthal. In the haplotype structure, each row represents a haplotype, and the column denotes a SNP (black lines indicate the presence of alternative allele).

To examine the biological implications of adaptive introgression in non-African populations, first we performed a Gene Oncology (GO) biological processes^74^ enrichment analysis of Neanderthal AI candidates using the *Enrichr* tool^75,76^. We combined the Neanderthal AI candidates identified by *MaLAdapt* in all 19 non-African populations into 4 superpopulations as defined by the 1000 Genomes study. Namely, we grouped the populations as Europeans (EUR), East Asians (EAS), South Asians (SAS) and Americans (AMR). We found that on a global level, introgressed variants from the Neanderthals played a key role in facilitating biological processes involved in metabolism regulation, adaptation to environments, and immune responses (Figure 7).

**Figure 7:**
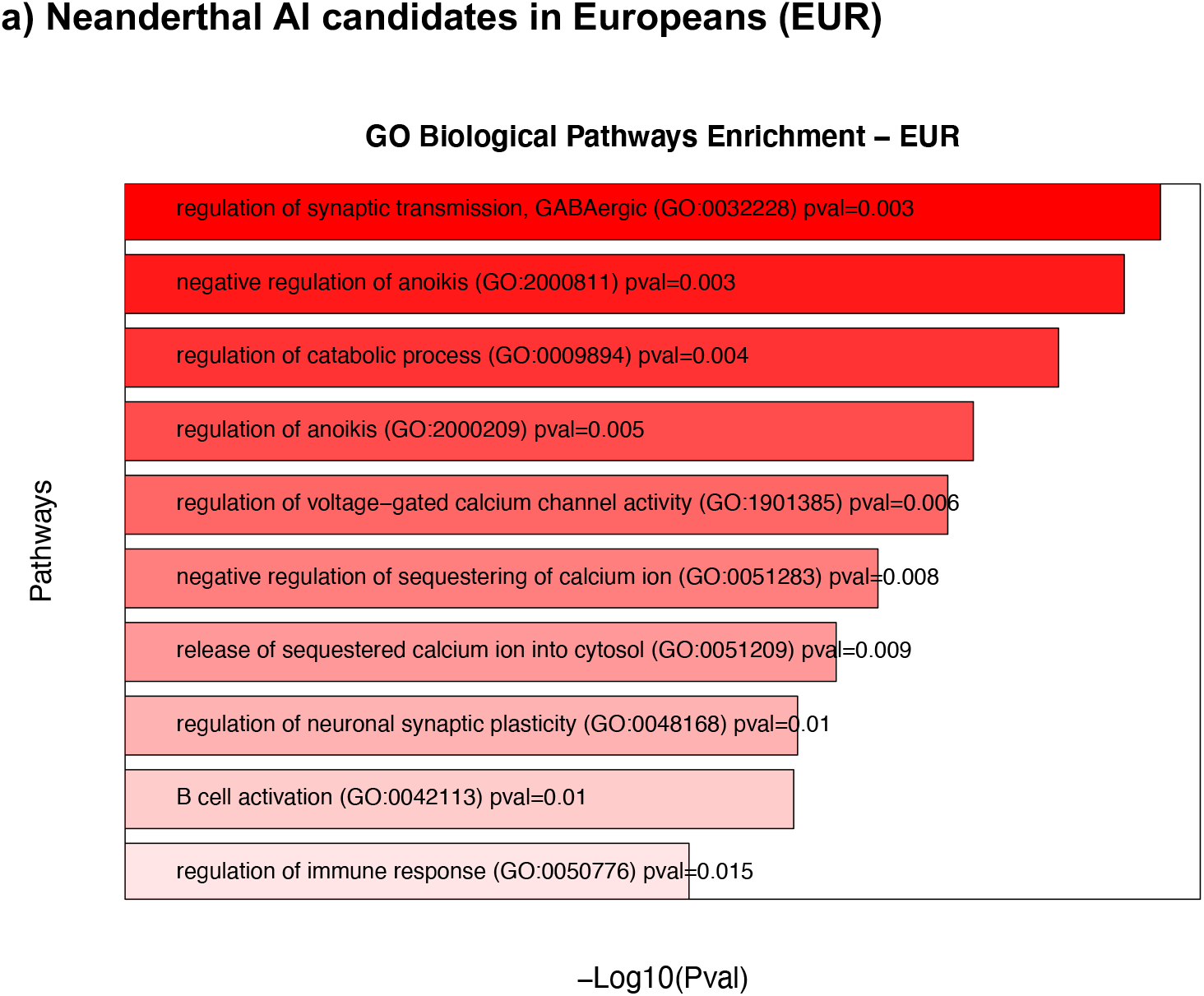
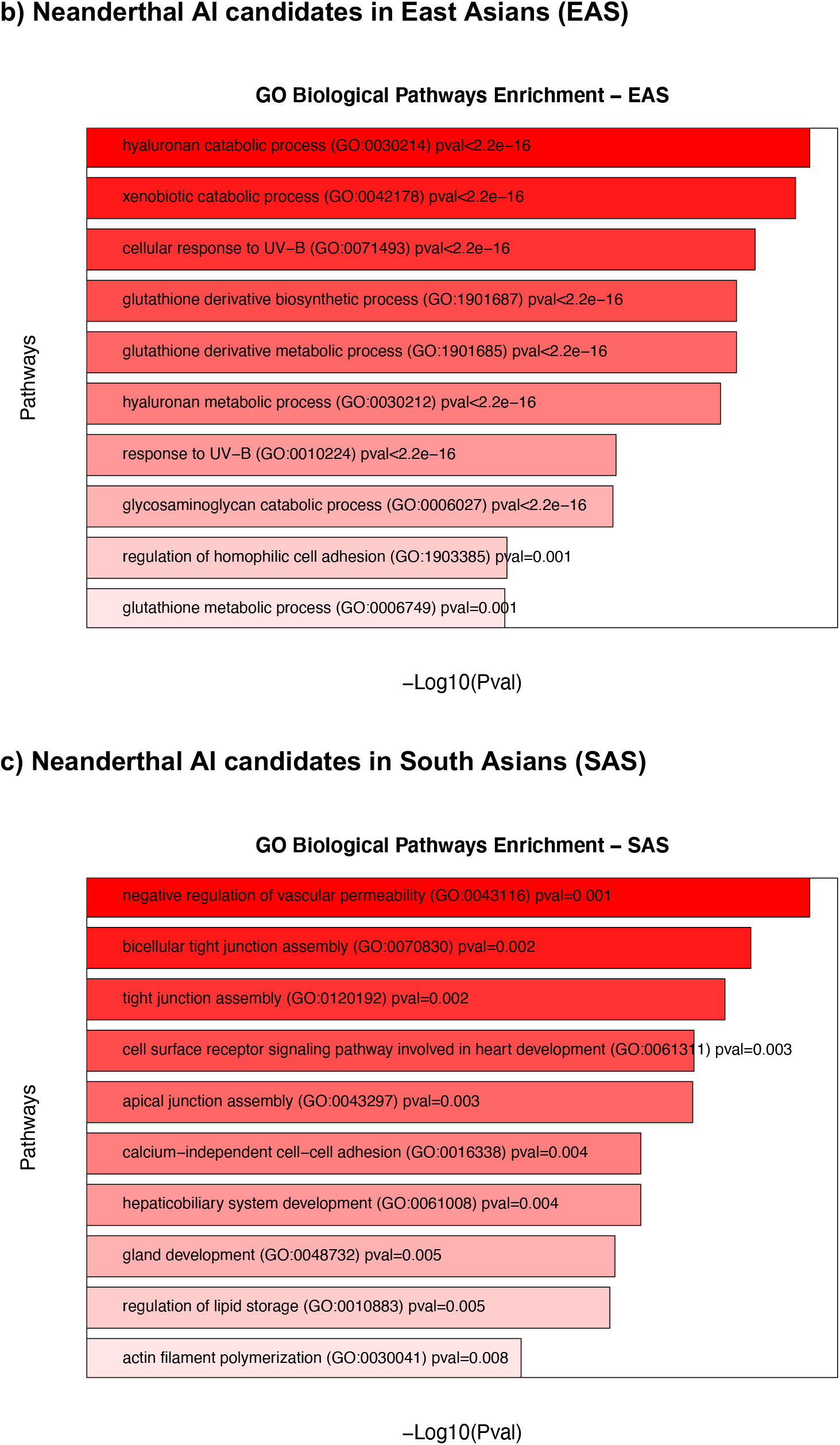
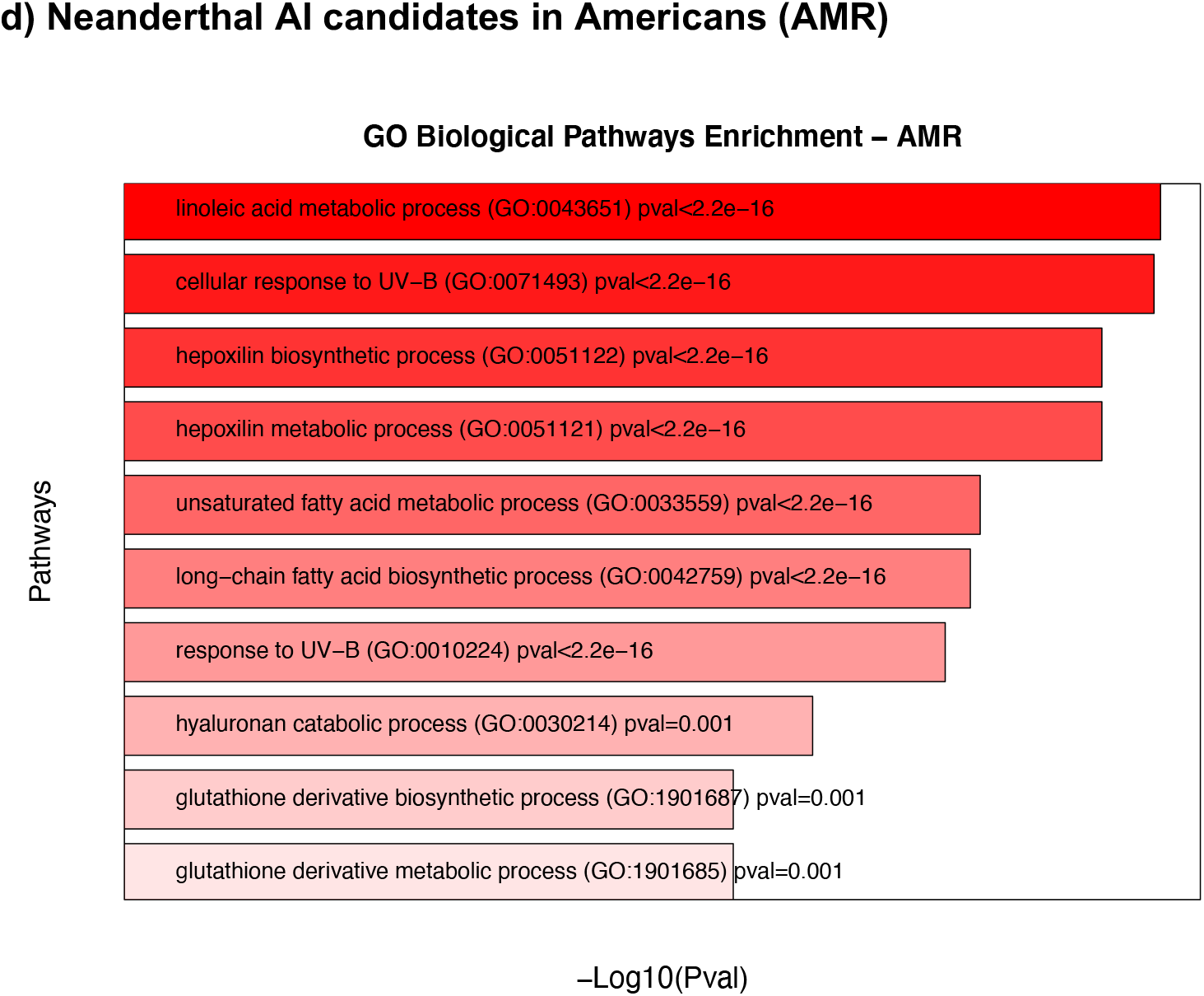
Gene Ontology (GO) Enrichment by Neanderthal adaptive introgression candidates in worldwide non-African superpopulations. We performed Gene Ontology enrichment test of Gene Ontology biological pathways using all AI candidates predicted in Europeans (EUR, Panel a), East Asians (EAS, Panel b), South Asians (SAS, Panel c), and Americans (AMR, Panel d) when using Neanderthal as the donor and African (YRI) as the outgroup population. We show the top 10 pathways in the enrichment test of each population (all pathways that reached significant p-values can be found in Supplementary Table 7). The length and the color intensity of bars indicate the significance of p-values, with the bar length being -log10(p value).

Next, we compared the distribution of Neanderthal AI probabilities as predicted by *MaLAdapt* in genes that code for proteins that interact with RNA viruses (the VIP genes) to other genes and genomic regions. Previous work suggests that the RNA viruses drove the adaptive introgression between Neanderthals and modern humans^77^. Although we find a slight enrichment of AI in VIP genes compared to non-VIP genes (Supplementary Figure 14-15), this difference is not significant (Supplementary Table 8, Fisher’s exact *p*-value=0.846, odds ratio=1.060). However, VIP genes that were reported as AI candidates^77^ show a substantial increase in AI probability in Europeans when compared to the genomic background (*p*-value < 2.2e-16) and other VIP genes (*p*-value < 2.2e-16), further validating our method’s power.

## Discussion

In this study, we present *MaLAdapt* – a machine learning algorithm for detecting signals of adaptive introgression from genome-wide data. Compared to existing methods, such as approaches based on summary statistics, *MaLAdapt* has more power to detect AI, despite the challenges presented by a highly imbalanced class ratio. It is also particularly good at detecting mild, incomplete AI regions, and is robust to most model misspecifications and non-AI sweeps. We have applied *MaLAdapt* to genetic variation data from modern human populations outside of Africa, most of whose ancestral populations experienced at least one archaic introgression event. In doing so, we have discovered AI candidate regions in all non-African populations from both Neanderthals and Denisovans, including novel AI candidates that have not been reported by previous studies.

A key challenge for ML methods is that the deterministic mechanism for the trained model typically remains unknown. Here we address this issue by using biologically meaningful features in the model, and use a decision tree-based algorithm so that the importance of all features in making predictions can be retrieved. By ranking the features by their importance scores (Figure 8, Supplementary Figure 2), we optimize the model by performing feature selection, and in doing so, obtain biological knowledge of adaptive introgression by examining key features being used in the predictions. We show that, the exon density and recombination rates played a critical role in *MaLAdapt’s* underlying prediction mechanism, as both factors jointly determine the extent of heterosis effect^44,46,78^. Additionally, summaries of genetic diversity, such as the number of segregating sites and heterozygosity, are also important factors to distinguish adaptive introgression from other population genetic processes.

One major challenge in genome-wide studies of AI is that the proportion of genome undergoing AI is likely to be substantially smaller than the part of the genome not experiencing AI, resulting in the so-called imbalanced class ratios. If the class ratio is extremely imbalanced, it can lead to an inflated False Discovery Rate (FDR) when performing multiple comparisons. This of course is a general statistical challenge in genome-wide studies. Depending on the signature of interest, different types of studies have used different strategies to account for the multiple testing issue. For example, GWAS studies typically use Bonferroni correction^79–81^ to obtain a genome-wide significant *p*-value threshold of 5e-8^82–84^, which efficiently controls the proportion of false positives in the outstanding signals. However, it can sometime be overly stringent and can lead to a high False Negative Rates^85^. Other ML or deep learning applications rely on the use of imbalanced datasets in the training process, followed by statistical corrections (*e*.*g. genomatnn* uses a beta correction to adjust class ratio in training and testing data sequentially). However, the main problem in this strategy is that none of the arbitrary ratios used in the training or testing data may be close enough to the empirical ratio. In the development of *MaLAdapt*, by utilizing a hierarchical structured algorithm with numerous randomly generated decision trees, we show that in our model, varying class ratios in the training data led to little change in the TPR and FPR (Supplementary Figure 4), so long as the trained model has learned from sufficient observations of both classes, as well as the confounders. To best evaluate the performance of methods on highly imbalanced empirical data, we apply *MaLAdapt* along with other related methods to full 5MB-long genomic segments, which class ratio is approximately 1:100 (*i*.*e*. 1 window of true positives to 100 windows of true negatives). We also show that at this ratio, *MaLAdapt* greatly outperforms all existing methods across all thresholds in terms of Precision, Recall, FPR (Figure 3). Even if the empirical ratio is more extreme than our testing data, all methods including *MaLAdapt* would suffer from a higher FDR, but *MaLAdapt* should still retain the highest precision among all.

Another major motivation for developing *MaLAdapt* is to control for potential false-positive signals due to recessive deleterious mutations in studies of AI. It is known from multiple previous studies^44,46,78^ that the presence of recessive deleterious mutations can lead to an increase in introgressed ancestry, similar to the manner of adaptive introgression, and thus is a confounder of AI detection. This effect is caused by heterosis or heterozygote advantage upon admixture, and is particularly pronounced in genomic regions that have high exon density and low recombination rates. Zhang et al. showed that existing methods for detecting adaptive introgression, such as the signature summary statistics^1,19,42,43^, can have exaggerated FPRs in such compact genomic regions when most deleterious mutations are recessive, and likely can explain the AI signature in *HLA* and *HYAL2* genes, which have been repeatedly discovered as AI candidates in European and Asian populations^26,86^.

*MalAdapt* attempts to control for this potential confounder of recessive deleterious mutations by including them in the simulations used to train the classifier. However, this training process is not without challenges. Similar to the class ratio discussed above, the main challenge for the potential heterosis confounding effect is that the degree of dominance of deleterious mutations in the human genome is poorly known. Most of the studies use models that assume all mutations are either strictly additive or fully recessive, while neither of these extreme assumptions reflect the empirical distribution of dominance. In *MaLAdapt*, we address the uncertainty in the dominance parameters by including three dominance models in the training data, which include an equal ratio of simulations where all deleterious mutations are additive, recessive, or partially recessive.

When applying *MaLAdapt* to empirical human population data, we do not detect *HLA* as an AI candidate in any of the populations. This suggests that *HLA* likely was a false identified AI candidate in previous studies^86–88^. However, although we did not detect AI at *HYAL2* in most Asian populations except one (CHB), we detected AI signatures in the upstream regions of *HYAL2* that overlap with multiple genes. A possible explanation for this observation is that the earlier reports of *HYAL2* being an AI candidate could have been due to linkage to another legitimate AI region upstream of it. However, future studies of the functional changes by the archaic variants in this region are needed to test this hypothesis. Furthermore, it is worth noting that the novel discoveries by *MaLAdapt* show similar distribution of exon density and recombination rates as previously identified AI candidates (Supplementary Figure 9-10), further supporting the conclusion that AI predictions made by *MaLAdapt* are not likely to be false positives due to heterosis from recessive deleterious mutations.

We compared the accuracy of *MaLAdapt* against other state-of-the-art AI detection methods, and noticed that two of the recently developed AI methods - the deep learning-based *genomatnn* and the polymorphism pattern-based *VolcanoFinder* – both suffered from substantial loss of power and robustness compared to what was originally reported when applied to our simulation data (Figure 3). When applied to empirical human genomic data, we noticed that more than half of the candidates predicted by *genomatnn* as well as *VolcanoFinder* received low prediction probabilities by *MaLAdapt* (Supplementary Figure 12). There are some essential differences between *MaLAdapt, genomatnn*, and *VolcanoFinder* that may explain the differences in their accuracy. For *genomatnn*, it is trained on simulations of short segments (100kb) that do not contain genic structure (coding/non-coding regions) similar to what is observed on the empirical human genome. *VolcanoFinder*, on the other hand, models the volcano shape of heterozygosity around the beneficial allele that is introgressed from a diverged population. This pattern is sensitive to adaptive introgression but could also be changed by other non-AI processes and the inherent characteristics of the genome, including the alignability and mappability of sequences. The simulations in our study used a considerable proportion of genomic regions with a high density of exons and low recombination rates due to concerns of heterosis effect and background selection^46,78^. In addition, the demographic parameters differ between the methods. For example, both *VolcanoFinder* and *genomatnn* assumed a fixed introgression amount and a fixed introgression time in their models, in contrast to *MaLAdapt, VolcanoFinder* is also optimized to detect AI with strong selection strength, whereas *MaLAdapt* considers weaker and recent sweeps on introgressed variants. Altogether, the reduction in power/accuracy could reflect the sensitivity of *genomatnn* and *VolcanoFinder* to mis-specification of the demographic model and genomic structures used by *MaLAdapt*.

To further disentangle the potential causes for the discrepancy in accuracy in different methods, we examined the exon density and recombination rates in the AI candidate regions in CEU predicted by *MaLAdapt, genomatnn* and *VolcanoFinder* (Supplementary Figure 11). The AI regions predicted by *genomatnn* tend to have both lower exon density and lower recombination rates than *MaLAdapt* and *VolcanoFinder* predictions, which are also lower than the whole-genome distributions. Next, we examined the haplotype structure of the *genomatnn* candidates using *Haplostrips* program (Supplementary Figure 18) that ranks European (CEU) and African (YRI) haplotypes by their affinity to the Neanderthal genome. To our surprise, the *genomatnn* candidates that received low *MaLAdapt* prediction scores also did not produce a clear AI pattern through this ranking of the haplotypes. This could be due to the fact that *Haplostrips* sorts and ranks the modern human haplotypes by distance to the archaic reference genome, which is different from the method of haplotype sorting in *genomatnn* that group haplotypes by populations. We visually inspected the haplotype structure patterns and annotated them as true positive, false positive, or uncertain labels (Supplementary Figure 13). We found that the *genomatnn* candidates that were not identified by *MaLAdapt* have strikingly low exon density and low recombination rates than the other two groups. In contrast, the visually false positive predictions by *MaLAdapt* are mainly driven by an excess of African (outgroup) haplotypes that also show close affinity to the archaic genome, in which case it is unclear whether it is a result of false detection or legitimate adaptive introgression due to back-to-Africa gene flow from Europeans^24^. Altogether, we believe *MaLAdapt* is more accurate in predicting AI in regions that contain a low number of mutations and few recombination events.

*MaLAdapt* can be used for the study of AI in other populations and organisms with different demographic histories and genomic structures. The simulation and training of *MaLAdapt* is easy to implement and computationally efficient, and is modifiable for other organisms. We provide all necessary scripts not only to replicate our results, but also for modifying the trained model for other population genetics studies. However, application of *MaLAdapt* to other systems requires several additional pieces of information that may not always be available. First, an accurate demographic model of the donor and recipient populations is necessary. For example, *MaLAdapt* currently relies on a well-understood Eurasian population history as its demographic model backbone. This model may not accurately describe the evolutionary history of human populations distantly related to Eurasians, such as the Americans. Further, the current model does not account for the complex demography in some of the regional populations, especially in Asia and Oceania where populations are known to have experienced complex archaic introgression and admixture patterns^6,8,9,11^. However, since *MaLAdapt* can be easily retrained, we expect to continually revisit and revise our model, when better-characterized demographic models for regional human populations become available. And despite the possible deficiencies of the demographic model in simulations, *MaLAdapt* demonstrates its power and accuracy by recovering most of the canonical AI candidates that have been reported by previous studies.

Another requirement for the use of *MaLAdapt* is an archaic reference genome. The empirical findings reported in this study are based on using the Altai Neanderthal individual^3^ as the Neanderthal reference genome, and the Altai Denisovan^5^ as the Denisovan reference genome. Without further discovery of more high-quality archaic hominin genomes, we do not have power to detect AI from unknown, “ghost” introgressions^24,54^ from archaic hominin that are distantly related to either Neanderthal or Denisovan. Nevertheless, we discovered numerous novel AI candidates in all non-African populations by Neanderthals and/or Denisovans that went undetected in previous studies, and have been verified by visual inspection of the haplotype structure^73^ (Figure 6). These genes are enriched in a wide range of biological pathways, which shed light on the functional influence of archaic introgression in general and their contributions to the phenotype spectrum, local adaptation, and health in our species. We provide a comprehensive summary of AI candidates in all non-African populations, with informative annotations of studies that reported them. We hope this can serve as a useful resource for future studies that are interested in the function and evolutionary history of specific genes of interest, especially for the novel AI discoveries in understudied populations with unique archaic ancestry distribution, such as the East Asians and South Americans.

In conclusion, *MaLAdapt* provides an example of how machine learning, especially feature-based algorithms, can help solve complex population genetics and human genomics problems. Such ML models can particularly be powerful at tackling questions with highly imbalanced classes, mild signals, and various confounding factors. We look forward to integrating new knowledge of archaic genomes and human evolutionary history into the *MaLAdapt* model, and to seeing novel methods at detecting AI in other biological systems inspired by *MaLAdapt*.

## Materials and Methods

### Simulation settings

We used the software SLiM (version 3.2.0)^65^ throughout this work for the simulations. We simulated adaptive introgression between archaic humans and modern humans under a three-population demographic model, shown in Figure 2 and Supplementary Table 1. This demographic model is adapted from Gravel et al. 2011^62^ and Prüfer et al. 2017^2^. In this demography, an archaic hominin population (*N*_*arc*_ = 1,000) splits from the ancestral African population (*N*_*anc*_ = 7,300) at 16,000 generations ago. The ancestral African population further splits into a modern African population at 5,600 generations ago (*N*_*afr*_ = 14,470) and a modern Eurasian population at 2,040 generations ago (*N*_*eur_OoA*_ = 1,861). The Eurasian population further experiences a population bottleneck at 920 generations ago (*N*_*eur_split*_ = 550), representing the split of European and East Asian populations, followed by a population expansion at exponential rate of 0.55% per generation until the end of the simulation. In the archaic population, a beneficial mutation with a selection coefficient (*s* ∈ [1e-4, 1e-2]) arises in an exon of the simulated genomic region at 15,000 generations ago and is simulated as fixed in the archaic population by introducing the mutation to all haplotypes. A single pulse of introgression occurs at a random time (*T*_*adm*_ ∈ in [1530, 2030]) at a random proportion (*m* ∈ {1%, 2%, 5%, 10%}). The introgressed beneficial mutation does not necessarily become immediately beneficial in the Eurasian population, depending on the selection time (*T*_*sel*_ ∈ [610, T_adm_-1]). All simulations are conditioned on the introgressed beneficial mutation not being lost in the recipient Eurasian population by the end of simulations.

We simulated 1,000 randomly sampled genomic regions from the modern human genome build GRCh37/hg19 with length of 5MB. As such, the simulated segments represent the empirical distribution of exon density and recombination rates on the human genome so that the inference of *MaLAdapt* accounts for the confounding effect by heterosis due to recessive deleterious mutations^46^. Specifically, we use the exon ranges defined by the GENCODE v.14 annotations^60^ and the sex-averaged recombination map by Kong et al.^61^ averaged over a 10kb scale. The per base pair mutation rate was fixed at 1.08e-8. Deleterious mutations can only occur in exonic regions of the segment with fitness effect drawn from a distribution estimated from modern humans^89^, with a shape parameter of 0.186 and average selection coefficient of -0.01315, as well as a 2.31:1 ratio of nonsynonymous to synonymous mutations^90^. Additionally, to account for the heterosis effect in the inference of adaptive introgression while accounting for the fact that the dominance distribution on the human genome is poorly understood, we simulated three models of dominance effects. In the first model, all deleterious mutations were fully additive (*h*=0.5). In the second, all were fully recessive (*h*=0). In the third model, all were partially recessive (*hs* relationship)^91^, where more strongly deleterious mutations were more likely to be recessive. For each of the sampled genomic segments, we repeated simulations 1,000 times under the Figure 2 demography using a given dominance model (deleterious mutations being additive, recessive, or partially recessive). Because there are three dominance models and 1,000 sampled segments in total, this exercise resulted in 3×1,000×1,000 = 3 million simulation replicates.

For computational efficiency of the simulations, we scale the simulation parameters by a scaling factor of c (c=10). In all simulations, the population size is rescaled to N/c, generation times to t/c, selection coefficient to s*c, mutation rate to μ*c, and the recombination rate to 0.5(1-(1-2r)c). Other evolutionary parameters remained the same.

### Features used by *MaLAdapt*

We consider biologically meaningful summary statistics that are likely informative of archaic adaptive introgression. The untrained *MaLAdapt* model learns which features are most important. All statistics are calculated in Python3. For each simulation replicate, we compute features in sliding 50kb windows (step size 10kb) throughout the simulated segments. We used 50kb as the prediction window size because it encompasses the average archaic introgressed haplotype length in modern humans, which is approximately 44kb^3^. We define adaptive introgression (label “AI”) as genomic windows in the admixed Eurasian population that contain beneficial mutations originating from archaic introgression. In contrast, windows with label “non-AI” do not contain the beneficial mutation, even if such windows are on the same genomic segment as the “AI” windows. Therefore, at most only 5 out of 496 windows per segment contain the beneficial mutations.

A full list of features used by the *MaLAdapt* can be found in Table 2, which include summary statistics that are informative about archaic introgression^1,42,43^, positive selection^19,92^, linkage disequilibrium^93–96^, genetic diversity^97–100^, and the genic structure and recombination rates^60,61^.

**Table 2:** Features used by *MaLAdapt*. From left to the right, this table summarizes the features used by MaLAdapt, including the biological signature they capture, notation in model, and a brief description.

| Information | Statistics | Description |
| --- | --- | --- |
| Archaic Introgression | $D$ | ABBA-BABA statistics |
| | $fD$ | |
| Adaptive Introgression | $R_D$ | Sequence divergence ratio |
| | $U_{20/50/80}$ | Number of uniquely shared alleles |
| | $Q_{90/95}$ | Quantile of derived allele frequency distribution |
| Selection | $H_{12}, H_2/H_1$ | Haplotype homozygosity |
| Spatial structure | $r^2$ | Linkage disequilibrium |
|  | **ZnS** |  |
| *Genetic Diversity* | **2pq** | Expected heterozygosity |
|  | **S** | Number of segregating sites |
| | $\theta_H, \theta_S, \pi$ | Estimates of $\theta$ |
| Genic Structure | **e** | Exon density |
|  | **r** | Recombination rate |

### Training *MaLAdapt* and the choice of the ETC algorithm

Using features computed from all windows in all simulated replicates, we further divided the dataset into training and testing datasets at 9:1 ratio. For the training dataset, we added additional segments containing selective sweeps due to de novo beneficial mutations. As these windows were not due to AI, these simulations were added to the “non-AI” labels. Up to 10% of the training dataset was comprised of these particular windows. In these selective sweep simulations, the beneficial mutations are de novo mutations in the Eurasian populations (rising at *T*_*sel*_), rather than introduced by archaic introgression. In the testing data, the original simulation class ratio (*AI*:*non-AI* ∼ 1:100) and genomic segment structures are preserved. In the training data, on the other hand, we shuffle the dataset to break down the genomic structure of the segments, and we further evaluate the influence of class ratios on the performance of *MaLAdapt* (Supplementary Figure 4). We show that in the training data, a relatively balanced class ratio optimizes the performance of *MaLAdapt* as the model is trained by observing sufficient examples of both classes. Therefore, we downsize the “non-AI” labeled windows to be twice the amount of the “AI” labeled windows. The final training data contains “AI” and “non-AI” windows at approximately 1:2 ratio.

We compared the performance of five machine learning algorithms to be used in *MaLAdapt* including Logistic Regression, LASSO, Ridge, traditional Random Forest, and ETC. The algorithms are trained and tested using the same datasets, and are evaluated in terms of different performance metrics including the True positive rates (TPR), False positive rates (FPR), Precision (1-False Discovery Rates), Recall (TPR), and F1 Score at different prediction probability thresholds (Supplementary Figure 1). We show that ETC is the best-performing algorithm at detecting genome-wide adaptive introgression, as its hierarchical structure is optimized at detecting mild adaptive introgression signature, especially when the class ratio is highly imbalanced. Therefore, we chose to use the ETC algorithm.

### Feature selection for model optimization

We additionally performed a feature selection process based on the feature importance score ranking from the original ETC-based *MaLAdapt*. We first determined 6 sets of features that contain different subsets of all 39 summary statistics given the feature importance scores from the pre-feature selection version of *MaLAdapt* (Supplementary Figure 2): 1) top high-ranking features (18 in total); 2) top high-ranking features minus *Qmax* (17 in total); 3) mid-ranking features (top features minus the *Q* stats; 18 in total); 4) all features minus the *Q* stats (37 in total); 5) all features minus the *Qmax* stat (38 in total); 6) all features (39 in total). For each model trained by a unique set of features, we apply them to the same testing data and evaluate the accuracy of predictions (Supplementary Figure 3). We show that despite all models have consistently low false positive rates (FPRs) across most prediction thresholds, the performance on other accuracy metrics, such as true positive rates (TPRs), false discovery rates (FDR) and F1 score (harmonic mean between precision and true positive rates), varies substantially between sets of features. We chose a subset that contains most of the summary statistics except the *Q* statistics (“set4”) to be the features included in the final version of *MaLAdapt* because of its low false discovery rate and the best F1 score across all thresholds. We use this version as the trained model reported in this study and for further application to empirical data.

### *MaLAdapt* robustness and model misspecification analysis

To evaluate the robustness of *MaLAdapt* to model misspecifications, we obtained a different set of testing data that includes 6 independent scenarios where one of the key parameter variables in the simulation model is perturbed (Supplementary Table 1). Specifically, we define *1) “T*_*sel*_*_low*” as the selection time being 200 generations lower than the original lower bound, 2) “*m_low”* as the introgression fraction (*m*) being 2-fold lower than the original lower bound, 3) “*m_high*” as the introgression fraction (*m*) being 2-fold higher than the original upper bound, 4) “*s_high*” as the selection coefficient (*s*) being 10-fold higher than the original upper bound, 5) “*segment*” as the genomic segments in simulations being different from the training data, and 6) “*demo*” as the Eurasian population growth rate and Out-of-Africa bottleneck size being different than the training simulations. We did not explore the selection coefficient (*s*) being smaller than the original lower bound due to extremely low chance of generating sensible amount of successful AI simulations conditioned on the beneficial mutation not being lost by the end of the simulation. We also did not perturb the time of introgression as its range is bounded by the split time between Eurasians and ancestral Africans, as well as the split time between Europeans and Asians.

We applied *MaLAdapt* to each of the above 6 perturbation testing datasets, and computed accuracy metrics including False Positive Rate (FPR), Precision, True Positive Rate (TPR, Recall), and F1 score with prediction probability threshold being at 0.9. We compared the metrics with the values obtained from *MaLAdapt* applying to the original testing dataset (without parameter perturbation), and compute the log10-fold change of the metrics to the original values.

### Analysis of AI in the 1000 Genomes Data

For the application of trained *MaLAdapt* on empirical modern human population data, we scanned the autosomes of human genomes data from Phase 3 of the 1000 Genomes Project, and computed the features used in Table 2 in 50kb sliding windows (step size = 10kb). Specifically, we first defined the genomic coordinates of the sliding 50kb windows throughout each of the autosomes (excluding the telomere and centromere regions). Within each window, we use the start and end position to extract the genotypes from the Yoruba (YRI, phased) as the non-introgressed population/outgroup, one of the 19 non-African populations (phased) as the introgressed population/recipient group, and one of the high-quality archaic genomes (Altai Neanderthal^3^ or Altai Denisovan^5^, unphased) as the introgressing population/donor group. We join the genotypes together as a matrix, and additionally removed sites in the archaic genomes having potential quality issues (quality score < 40 and/or mapping quality < 30). We computed all summary statistics included in the feature set in *MaLAdapt*, and repeated the process across all windows across all autosomes. We computed features for Neanderthal introgression and Denisovan introgression separately for all populations. We applied the trained model to all 19 non-African populations and obtained prediction probabilities in all windows across the whole genome for Neanderthal or Denisovan adaptive introgression, respectively. We further converted the prediction probability of *Pr(AI)* to a prediction score, which equals -log10(1-Pr(AI)). We plot the prediction scores of all windows for each population, and label the gene names in AI regions.

## Author contribution

BK and AD conceived the study. XZ designed the study, carried out the simulations, machine learning implementation, empirical data analyses, and wrote the manuscript. BK, AD, SS, KEL contributed to the design of the study, data analysis, and participated in manuscript writing. BK designed the simulation framework. AS participated in code optimization and machine learning data analysis. All authors read and approved the manuscript.

## Acknowledgements and Funding Information

XZ was supported by NIH Grant K99GM143466 and UCLA Quantitative and Computational Biosciences (QCBio) Collaboratory Fellowship. BYK was supported by NIH grant F32GM135998. KEL was supported by NIH Grant R35GM119856. SS was supported by R35GM125055 and an Alfred P. Sloan Research Fellowship. We thank Dr. Graham Gower and Dr. Xiaoheng Cheng for sharing scripts of *genomatnn* and *VolcanoFinder*, and providing insightful comments related to the comparison of adaptive introgression inference results from different methods. We also thank Dr. David Enard for providing the VIP-related datasets for analyses in this study.

## Data Availability Statement

All scripts necessary to recreate the simulations, machine learning training and testing, robustness analysis, and empirical predictions can be found at GitHub: https://github.com/xzhang-popgen/maladapt

