## SupplementaryMaterial for "*MaLAdapt* reveals novel targets of adaptive introgression from Neanderthals and Denisovans in worldwide human populations"

### Supplementary Materials

#### Supplementary Methods

The comparison of performance between *MaLAdapt* and other AI detection methods

We obtained a separate set of testing data using 1000 randomly sampled 5MB genomic segments that are different from the 1000 segments used in the training process. We obtained 10 replicates of simulations per segment under the same simulation set-up for the training data, including the demographic model, parameters, and ranges where the variable parameters draw from. Features are computed over 50kb sliding windows (step size=10kb) across the segments after each iteration of simulation, and combined all replicates of all segments together as one complete testing dataset. We labeled windows that contained the beneficial mutation as class "*AI*" and otherwise as class "*non-AI*". We applied all methods, including *MaLAdapt* and other previously published AI detection methods to the same testing dataset for accuracy comparison purposes (Figure 3). For the predictions, we define True Positive as methods predicting "*AI*" in windows that contain the beneficial mutation, and accordingly, False Positive as windows being predicted as "*AI*" but do not contain the beneficial mutation. For methods like *MaLAdapt* and *genomatnn* that yield posterior prediction probabilities (between 0-1), accuracy metrics were computed across different probability thresholds. For other standalone statistics and *VolcanoFinder* (in which case, its prediction Likelihood Ratio) that are outlier-based methods, we used the percentile ranking of the statistical values as prediction probabilities, with more extreme percentiles indicating AI.

For the implementation of *genomatnn*, we trained a Convolutional Neural Network (CNN) model using 250,000 simulations with 64 rows in the genotype matrices, and applied the trained model to the VCFs of the aforementioned testing dataset. For *VolcanoFinder*, we computed allele frequency and site frequency spectrum of derived alleles in the VCFs of the testing dataset, and scanned for AI at every 1kb across the simulated segment, under Model 1 defined by *VolcanoFinder* and searching through a grid of valid genetic distances.

Visualizing haplotype structure of adaptive introgression candidate regions

To visualize the haplotype structure of AI candidate regions detected by *MaLAdapt* and other related methods, we used the program *Haplostrips* to plot the haplotypes. For each detected region, we extracted the haplotypes of all individuals in the respective non-African population as the AI recipient population and the Yorubans (YRI) as the non-introgressed, outgroup population from the Phase 3 of the 1000 Genomes Project. We additionally extracted the unphased genotypes of the Altai Neanderthal and the Denisovan individual at the candidate region as the donor population. We constructed a haplotype matrix consisting of the donor, recipient, and the outgroup population as the input for *Haplostrips* program. The program displays the haplotype structure with genomic variants within the region as columns, and haplotypes as rows. Each population was assigned a unique color on the left side of the haplotypes. The haplotypes are hierarchically sorted and clustered by a decreasing distance to the reference archaic genome (donor population). Supplementary Figure 5 shows the haplotype structures of all Neanderthal AI candidates in CEU population detected by *MaLAdapt*. Supplementary Figure 16 (zipped files) includes all *MaLAdapt* detected AI candidates' haplotype structures and their corresponding distribution of haplotypes in terms of genetic distance to donor population. Supplementary Figure 17 (zipped files) includes the haplotype structures of *genomatnn* detected Neanderthal AI candidate regions in CEU population plotted by *Haplostrips*.

#### Gene Oncology (GO) enrichment test and VIP gene analysis

We grouped the 19 non-African populations by their superpopulations defined by the 1000 Genomes Project, including the Europeans (EUR), East Asians (EAS), South Asians (SAS) and the Americans (AMR), and summarized all genes that overlap with the AI windows predicted in all populations within each superpopulation to get a unique gene list. We then used the unique gene list from each superpopulation as the input for the program Enrichr[1,2] to get the p-values and odds ratio of GO biological processes[3] enrichment. We extracted the top 10 processes in each superpopulation's test that reached statistical significance (defined as p-value < 0.05), and plotted the significance of the processes (measured by significance score, which equals -log10(p-value); Figure 7).

For the AI enrichment test in VIP genes, we first downloaded the VIP gene set and 1000 bootstrap-controlled non-VIP gene sets from Enard and Petrov 2018[4]. Additionally, we extracted a set of randomly selected genomic windows (with matched size of the VIP gene set) as the genomic background comparison, and a randomly selected human gene set (with matched size of the VIP gene set) as random gene comparison. For genes in the these datasets that span over multiple 50kb windows in the empirical European genomic data (CEU), we intersect the genes with all overlapping windows that received Pr(AI) predictions from *MaLAdapt*, and assign the Pr(AI) of each gene as the maximum Pr(AI) among the overlapping windows. Additionally, we extracted the VIP genes that are also AI candidates in CEU as suggested by Enard and Petrov 2018, and summarize the Pr(AI) in these genes as the VIP-AI set. We then plotted the distribution of Pr(AI) of all aforementioned comparison sets in Supplementary Figure 14, and computed the Wilcoxon p-values between pairs of datasets. We show that the AI probabilities in the VIP-AI set are significantly higher than all other comparison sets, validating the power of *MaLAdapt*. We further divide all human genes into 4 categories including VIP, non-VIP, AI, and non-AI, and performed Fisher's exact test on the contingency table (Supplementary Table 8). We show that although there is a subtle enrichment of AI signature in the VIP genes (odds ratio > 1), the enrichment is not significant (p-value=0.8467).

**Supplementary Table 1: All parameters in *MaLAdapt*-related simulations**

| Parameter | Description | Value in training | Value in specificity/robustness analysis |
|---|---|---|---|
| ***c*** | Scaling factor in simulations | 10 | |
| $m_{adm}$ | Archaic introgression rate | {0.01, 0.01, 0.05, 0.1} | *m_low* = 0.005, *m_high* = 0.2 |
| $T_{adm}$ | Time of archaic introgression | [1530, 2030] | |
| $T_{sel}$ | Time of positive selection on archaic introgressed mutation | [610, $T_{adm}$-1] | $T_{sel}$*_low* = 410 |
| ***s*** | Selection coefficient of archaic introgressed mutation (during positive selection) | [1e-4, 1e-2] | *s_high* = 0.1 |
| ***h*** | Dominance of deleterious mutations | {0, 0.5, *h(s)*}<br><br>*h(s)* = (0.5)/(1 + 7071.07*s) | |
| ***a*** | Shape factor of Gamma-distributed fitness effect in deleterious mutations | 0.186 | - |
| ***E[s]*** | Expected selection coefficient of Gamma-distributed fitness effect in deleterious mutations | -0.01315 | - |
| $R_{nonsyn-syn}$ | nonsynonymous to synonymous mutations ratio in coding regions | 2:31:1 | - |
| $T_{burn-in}$ ***(generations)*** | Burn-in time in simulations | 73,000 | - |
| $T_0$ ***(generations ago)*** | Archaic split time | 16,000 | - |
| $T_s$ | Beneficial mutation occurrence time | 15,000 | - |
| $T_{AF}$ | African population expansion time | 5,600 | - |
| $N_{Anc}$ | Ancestral population size | 7,300 | - |
| $N_{Arc}$ | Archaic population size | 1,000 | - |
| $N_{Afr}$ | African population size | 14,470 | - |
| $N_{OoA}$ | Eurasian population size during Out-of-Africa migrations | 1,861 | - |
| $N_{eur\_split}$ | Eurasian Population size during European-Asian split | 550 | *demo* = 1320 |
| $r_{Eur}$ ***(%)*** | | 0.55 | *demo* = 0.40 |
| $m_{Afr-Arc}$ | Migration rate between archaic and African populations | 15e-5 | - |
| $m_{Afr-Eur}$ | Migration rate between Eurasian and African populations | 0.78e-5 | - |
| ***L*** | Segment length | 5MB | - |
| $N_{seg}$ | Number of sampled segments | 1,000 | - |
| ***e*** | Exon density | GENCODE | Different segment sampling |
| ***r*** | Recombination rate | Kong et al. 2010 | Different segment sampling |
| $\mu$ | Mutation rate | 1.08e-8 | - |

| | | | |
|---|---|---|---|
| ***$N_{sim}$*** | Number of simulation replicate per set of parameters | 1,000 | 100 |

*This table lists all parameters used in the simulations for MaLAdapt training and testing purposes, including the parameters that are variables. From left to the right, the columns represent the parameter name, description, fixed values or ranges variables draw from in the simulations, and the perturbed values used in model robustness and misspecification analysis.*

**Supplementary Table 2: AI detection methods accuracy comparison across key thresholds**

| Method | Metric | thresh_0.5 | thresh_0.8 | thresh_0.88 | thresh_0.9 | thresh_0.95 |
|---|---|---|---|---|---|---|
| ***MaLAdapt*** | FPR | 0.079 | 0.003 | 0.001 | 0.001 | 0 |
| ***RD*** | | 0.477 | 0.178 | 0.104 | 0.086 | 0.042 |
| ***Q95*** | | 0.394 | 0.146 | 0.072 | 0.054 | 0.017 |
| ***U20*** | | 0.067 | 0.067 | 0.067 | 0.067 | 0.036 |
| ***U50*** | | 0.022 | 0.022 | 0.022 | 0.022 | 0.022 |
| ***MaLAdapt*** | Precision | 0.061 | 0.502 | 0.638 | 0.683 | 0.769 |
| ***RD*** | | 0.02 | 0.035 | 0.043 | 0.046 | 0.048 |
| ***Q95*** | | 0.08 | 0.165 | 0.263 | 0.31 | 0.533 |
| ***U20*** | | 0.125 | 0.125 | 0.125 | 0.125 | 0.083 |
| ***U50*** | | 0.263 | 0.263 | 0.263 | 0.263 | 0.263 |
| ***MaLAdapt*** | Recall | 0.963 | 0.585 | 0.475 | 0.41 | 0.045 |
| ***RD*** | | 0.848 | 0.569 | 0.414 | 0.365 | 0.188 |
| ***Q95*** | | 0.995 | 0.842 | 0.749 | 0.71 | 0.552 |
| ***U20*** | | 0.853 | 0.853 | 0.853 | 0.853 | 0.288 |
| ***U50*** | | 0.708 | 0.708 | 0.708 | 0.708 | 0.708 |
| ***MaLAdapt*** | F1 | 0.115 | 0.541 | 0.545 | 0.513 | 0.086 |
| ***RD*** | | 0.038 | 0.065 | 0.078 | 0.081 | 0.076 |
| ***Q95*** | | 0.148 | 0.276 | 0.39 | 0.432 | 0.542 |
| ***U20*** | | 0.218 | 0.218 | 0.218 | 0.218 | 0.128 |
| ***U50*** | | 0.383 | 0.383 | 0.383 | 0.383 | 0.383 |

*The ballmark accuracy values, including False Positive Rate (FPR), Precision, Recall, and F1 score, of MaLAdapt and other AI detection methods at different posterior prediction probability thresholds. The thresholds used by MaLAdapt at predicting empirical data are highlighted in red.*

**Supplementary Table 3: Area Under the Curve for ROC curve (AUROC) and Precision-Recall curve (AUPR) in AI detection methods**

| Method | AUROC (± SE) | AUPR (± SE) |
|---|---|---|
| ***MaLAdapt (AI)*** | 0.984 (±0.0002) | 0.519 (±0.004) |
| ***RD*** | 0.769 (±0.001) | 0.035 (±0.0002) |
| ***Q95*** | 0.938 (±0.0002) | 0.492 (±0.001) |
| ***U20*** | 0.888 (±0.001) | 0.097 (±0.001) |
| ***U50*** | 0.942 (±0.001) | 0.173 (±0.002) |
| ***genomatnn*** | 0.655 (±0.006) | 0.036 (±0.001) |
| ***VolcanoFinder*** | 0.644 (±0.003) | 0.056 (±0.0006) |

**We summarize area under the curves from Figure 3, and compute standard errors (SE) of the AUPR and AUROC using a jackknife method where we divide the test data into 1000 subsets and drop one subset each time to recompute the accuracy metrics.*

**Supplementary Table 4: Confusion Matrix of *MaLAdapt* performance on non-AI positive selection sweeps**

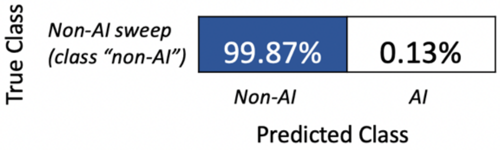

*The confusion matrix that shows the accuracy of MaLAdapt at positive selection sweeps that act on de novo mutations instead of AI. We show that at the posterior probability threshold of 0.9, MaLAdapt is highly robust to the confounding factor of non-AI sweeps by correcting assigning non-AI sweep as class "non-AI" 99.87% of times.*

**Supplementary Table 5-7: See Separate Microsoft Excel file.**

**Supplementary Table 8: Contingency Table for AI enrichment in VIP and nonVIP genes**

| Genes | *VIP* | *nonVIP* |
|---|---|---|
| ***AI*** | 9 | 28 |
| ***nonAI*** | 3483 | 11490 |

*Fisher's Exact Test p-value = 0.8467, odds ratio = 1.0604*

**Supplementary Figure 1: Performance of different machine learning models considered by *MaLAdapt* using the same training dataset**

**a) Extra-Tree Classifier (*MaLAdapt*)**

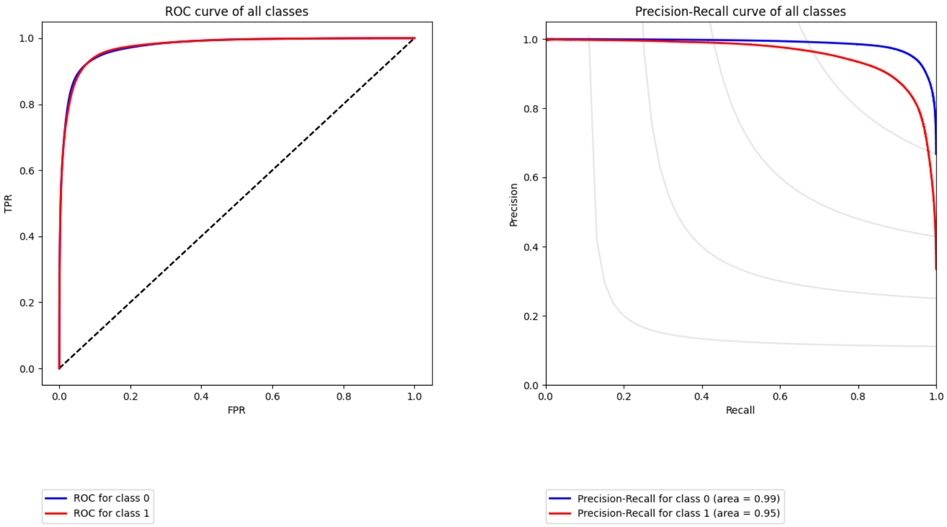

#### b) Random Forest

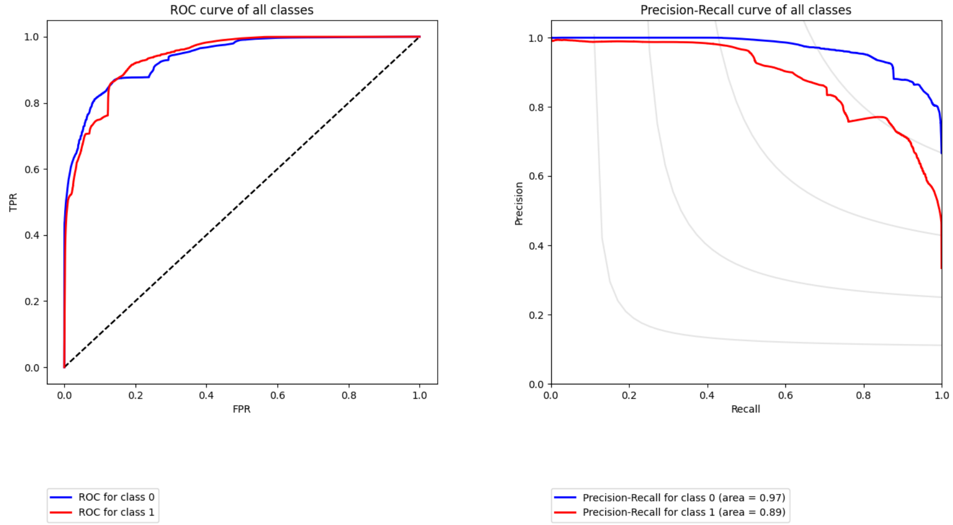

#### c) Logistic Regression

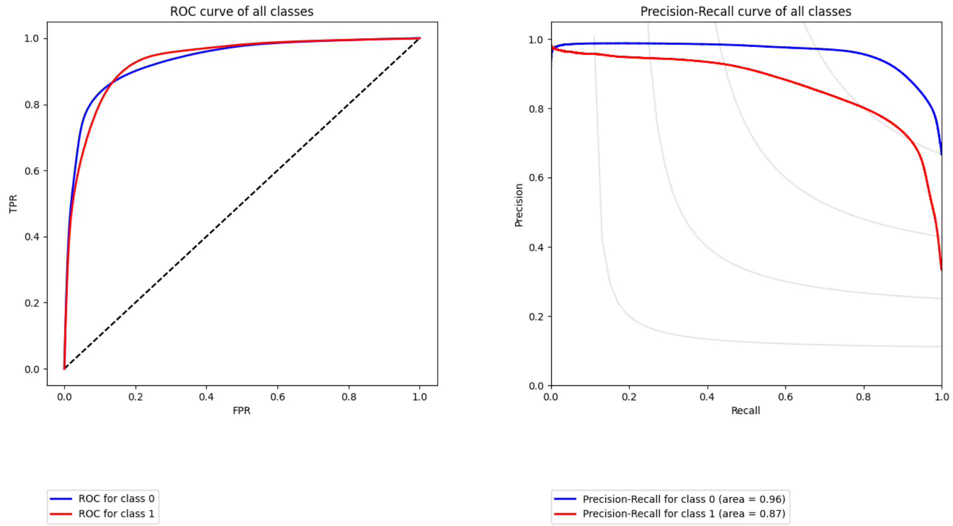

#### d) Logistic Regression – Lasso (L1 Penalty)

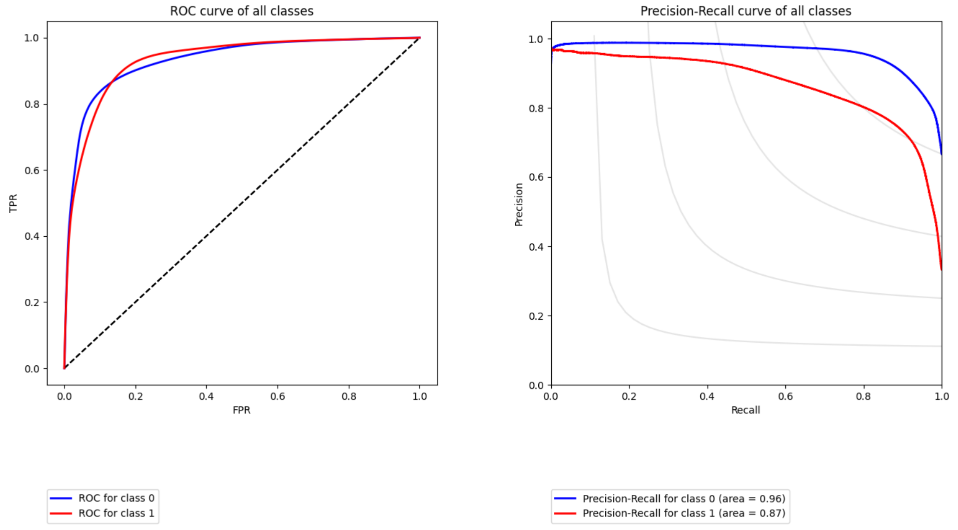

#### e) Logistic Regression – Ridge (L2 Penalty)

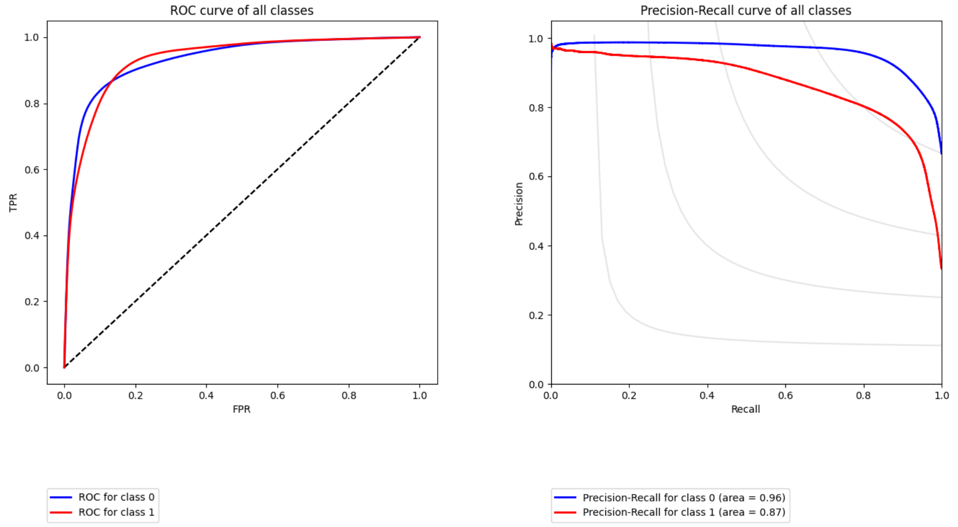

*This figure compares the accuracy of 5 different ML algorithms trained and tested on the same data (class ratio between AI:nonAI:sweep = 1:2:0.1). We chose Extra-Trees Classifier (panel a) for MaLAdapt for its highest power and precision.*

**Supplementary Figure 2: Feature importance score ranking in different subsets of features**

a) **Set 1: top high-ranking features (18 in total)**

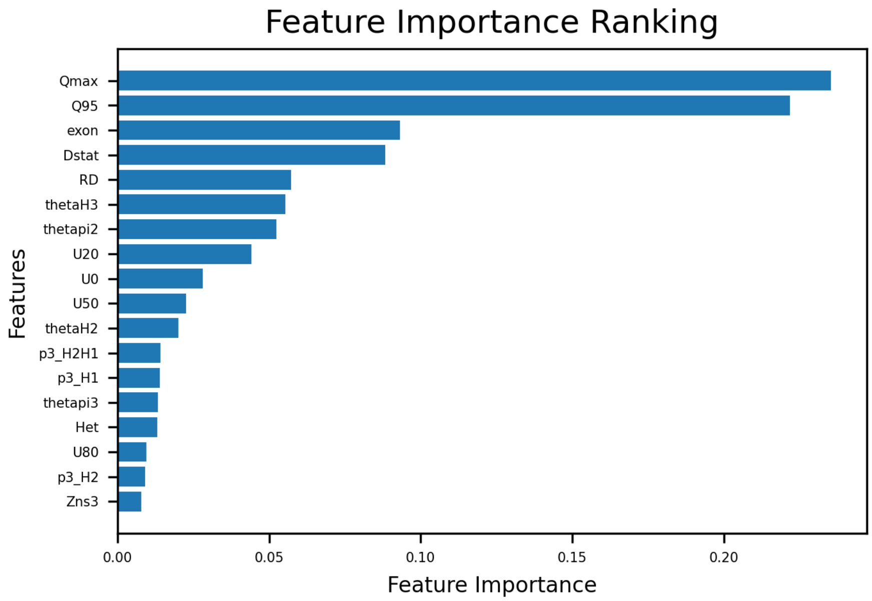

b) **Set 2: top high-ranking features minus Qmax (17 in total)**

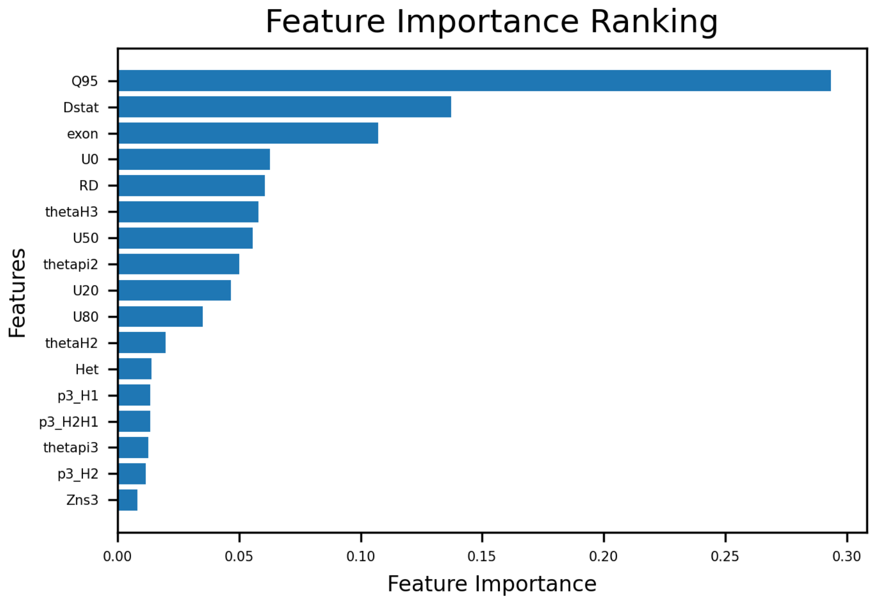

c) **Set 3: mid-ranking features (top features minus the Q stats; 18 in total)**

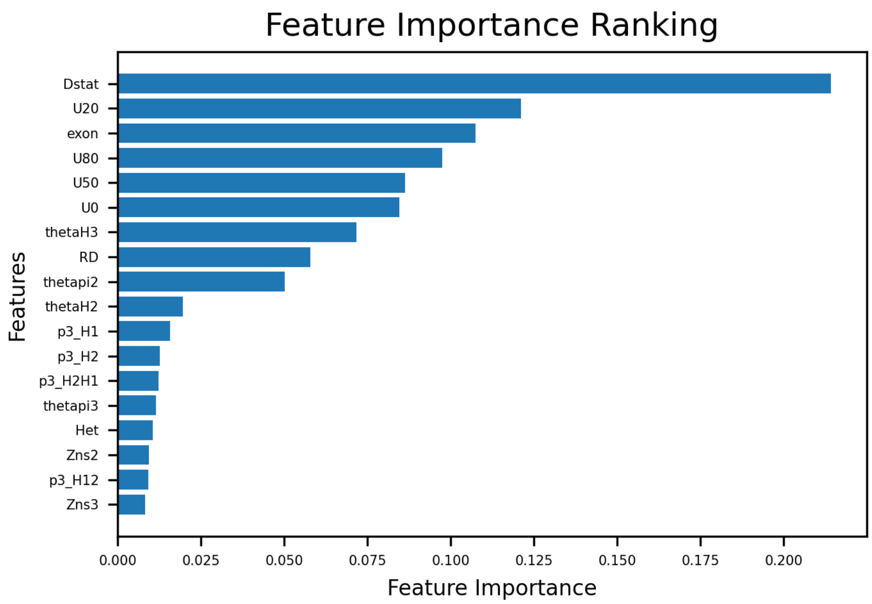

**d) Set 4 (*MaLAdapt*): all features minus the Q stats (37 in total)**

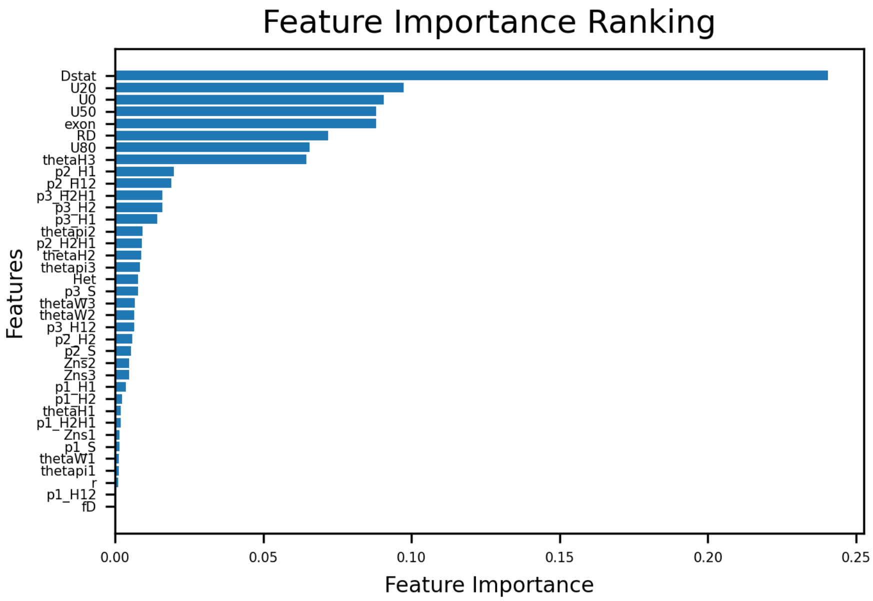

**e) Set 5: all features minus the Qmax stat (38 in total)**

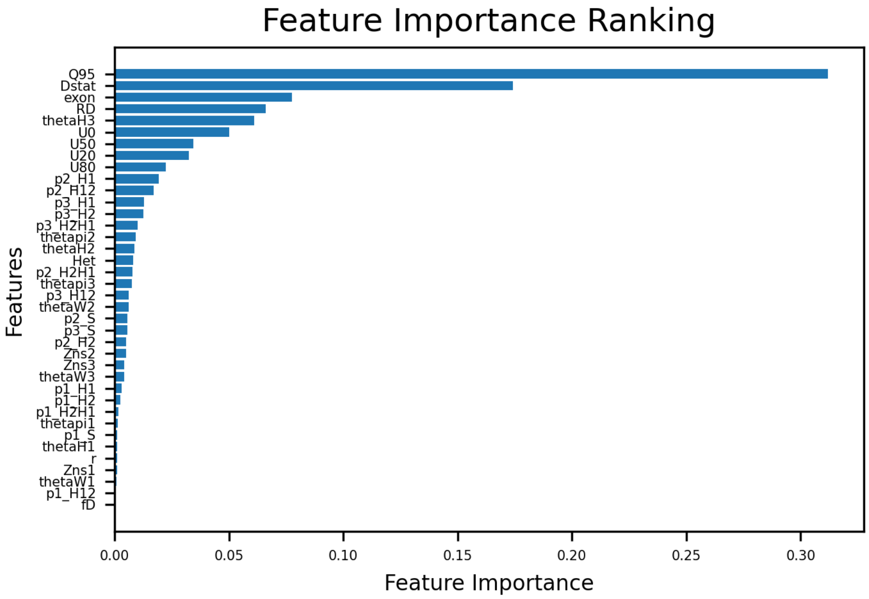

**f) Set 6: all features (39 in total)**

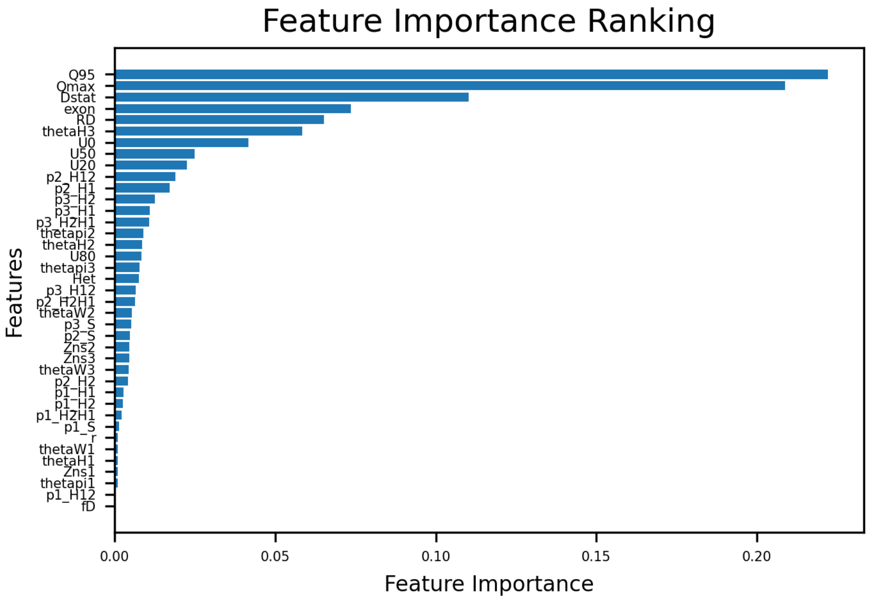

*We retrieve the feature importance scores (value between 0 to 1, all features sum up to 1) from MaLAdapt to understand its underlying deterministic model. First, we created 6 subsets of features that each accounted for the uneven distribution of feature importance differently. We ranked and plotted the importance of each of the 6 feature sets (panel a-f). After evaluating the performance and accuracy of 6 versions of MaLAdapt using each of the feature sets (Supplementary Figure 3), we concluded a final set of 39 features used by MaLAdapt (panel d).*

**Supplementary Figure 3: Performance of *MaLAdapt* trained with different feature sets (testing data class ratio 1:100)**

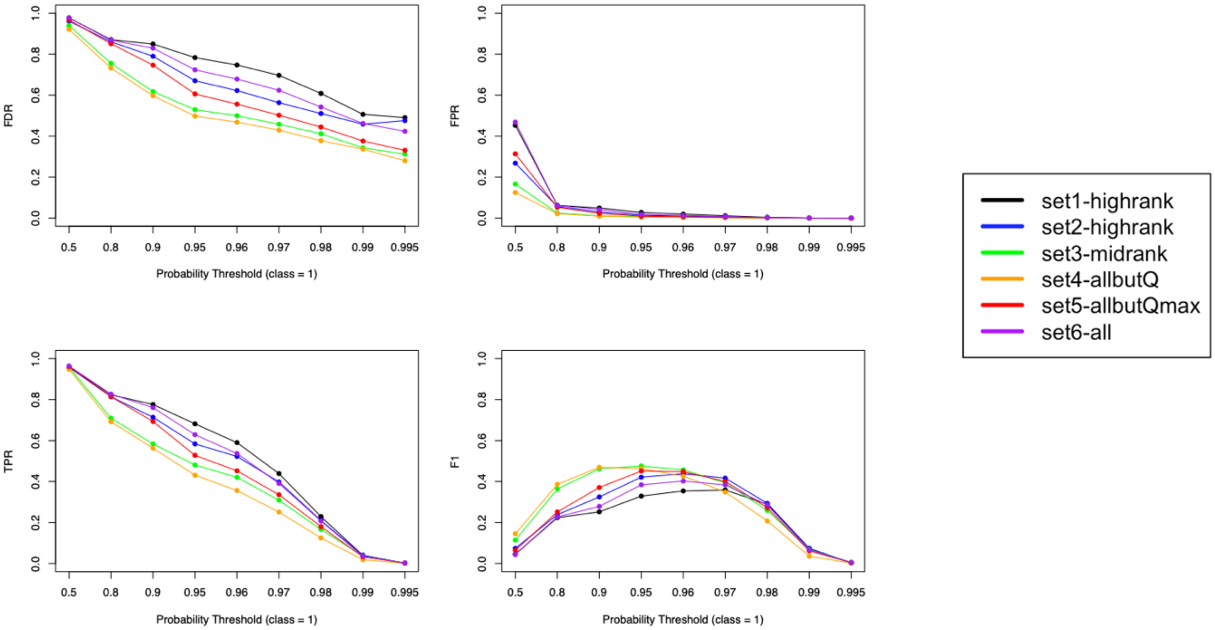

**This figure compares the accuracy metrics (TPR, FPR, FDR, F1 Score) of 6 versions of MaLAdapt using different feature subsets at posterior prediction probability thresholds ranging from 0.5 to 0.995. We show that feature set4 (final version of MaLAdapt) shows the highest accuracy across prediction probability thresholds. Furthermore, at probability threshold = 0.9, MaLAdapt has the highest F1 score (weighed mean of power and precision), justifying the use of this threshold for empirical predictions.*

**Supplementary Figure 4: Performance of *MaLAdapt* tested on different class ratios**

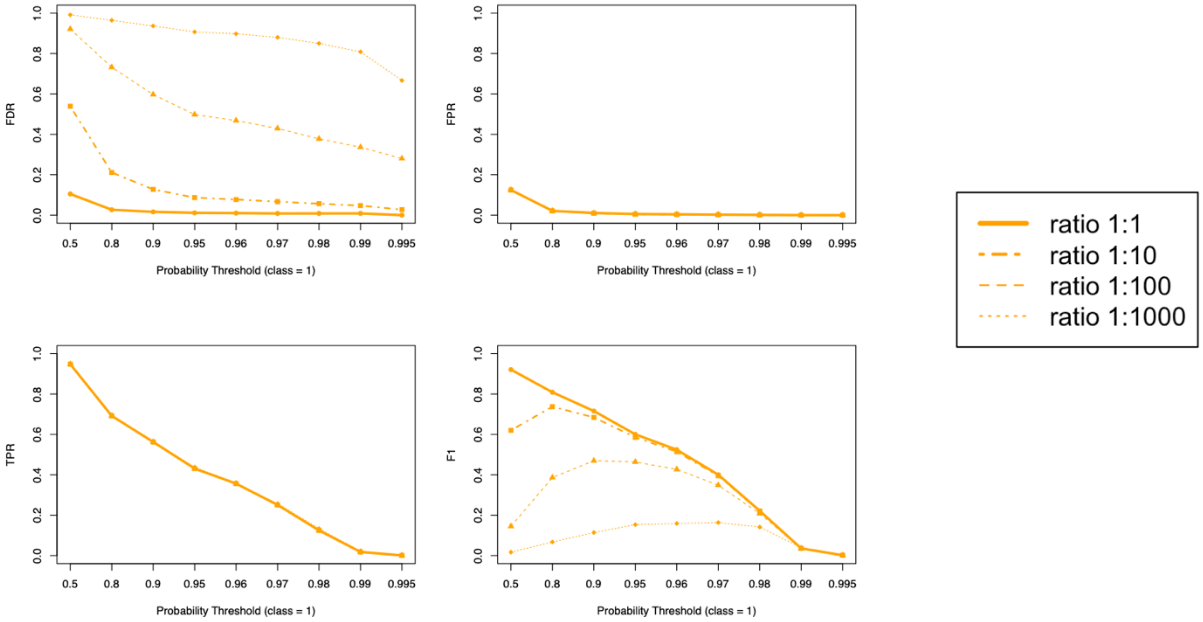

*We train and test MaLAdapt to different dataset with varying class ratios between AI and nonAI, and show that varying class ratios in the training data led to little change in the TPR and FPR, so long as the trained model has learned from sufficient observations of both classes, as well as the confounders.*

**Supplementary Figure 5: Distribution of novel Neanderthal AI in CEU haplotypes in terms of genetic distance to the donor population (Altai Neanderthal)**

**a) *S100PBP***

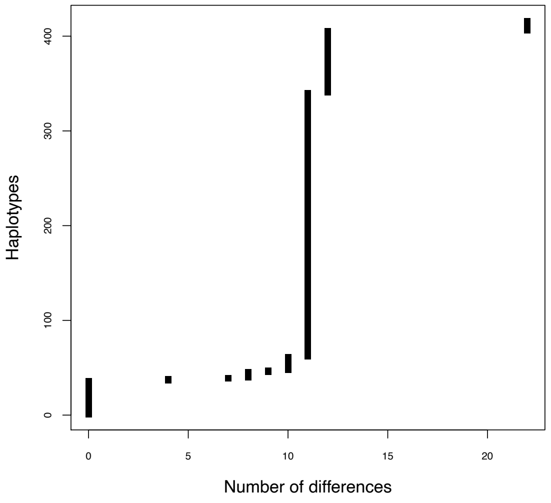

**b) *CRMP1***

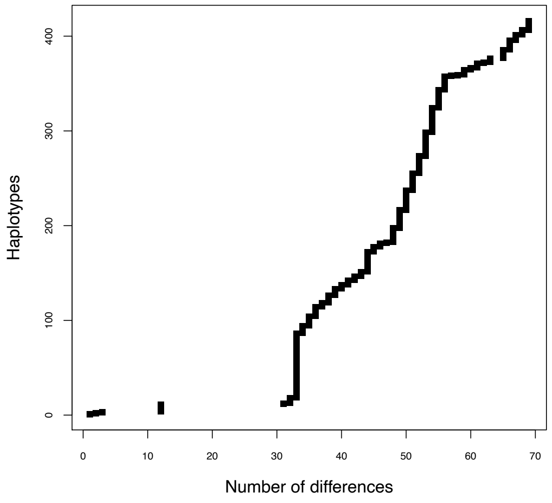

**c) *TMEM209***

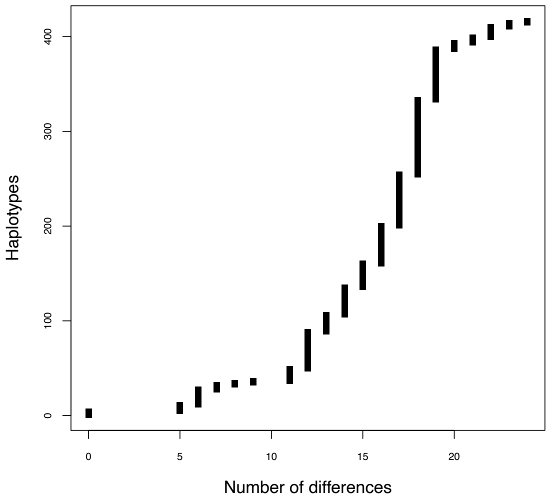

**d) *MEST***

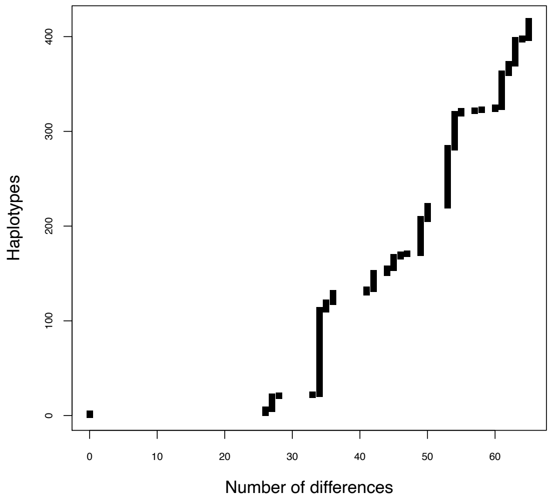

**e) *TG***

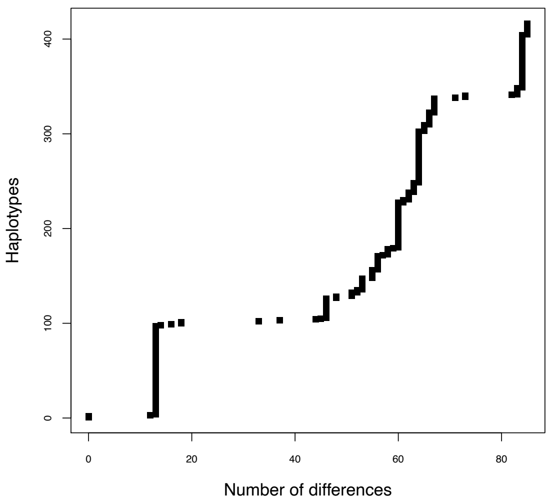

**f) *KLF6***

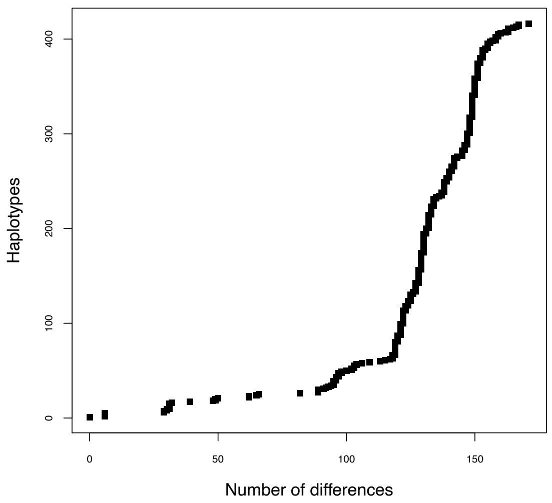

**g) Intergenic**

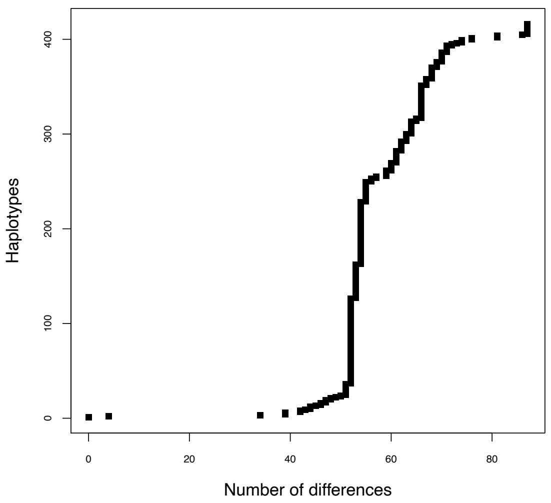

**h) *ADAM21P1***

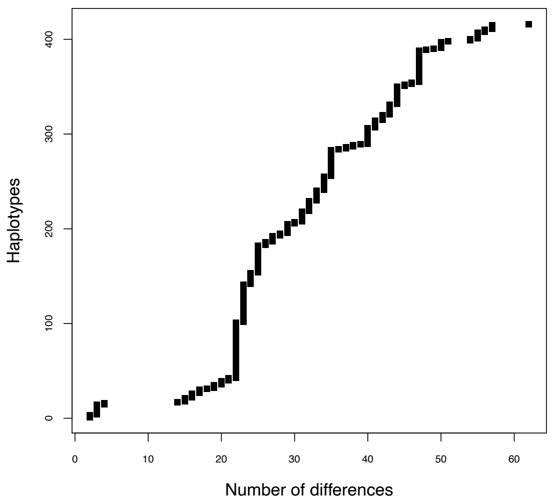

**i) *TCF12***

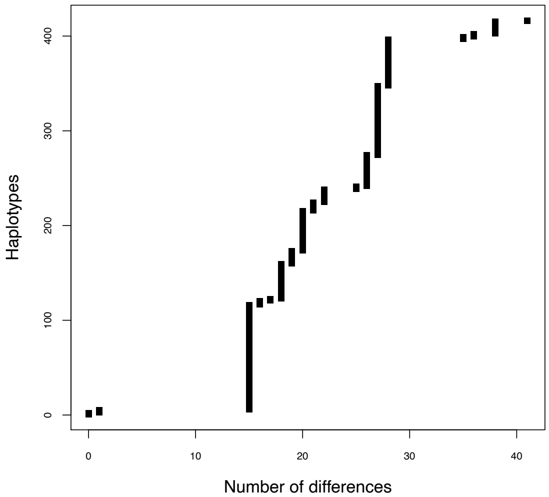

*We extracted the haplotypes of novel Neanderthal AI candidate regions in CEU, and plotted the distribution of haplotypes in the recipient populations (CEU) in terms of their genetic distance to the donor population (Altai Neanderthal). Each y-axis point indicates an unique haplotype, and the x-axis shows the number of differences between the donor haplotypes and the individuals in the recipient population sorted by decreasing similarity*

**Supplementary Figure 6: *MaLAdapt* prediction probability distribution of AI candidates identified by previous studies but missed by *MaLAdapt***

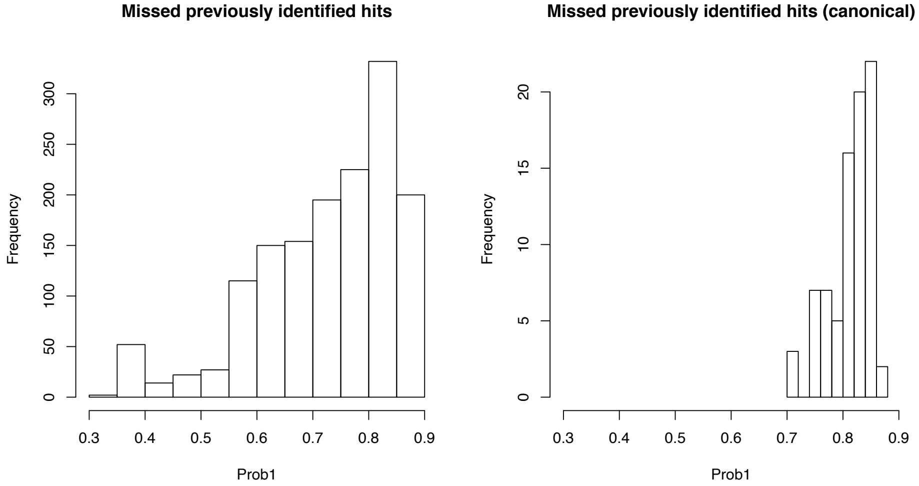

*We examined the distribution of Pr(AI) on AI candidates that have been identified by previous studies but were not predicted as AI by MaLAdapt (probability threshold = 0.9). The left panel shows the Pr(AI) distribution of all previously identified AI candidates, and the right panel shows the Pr(AI) distribution of the "canonical" AI hits, defined as candidates that are suggested as AI by at least 2 studies. We show that even when MaLAdapt missed the canonical AI hits, it still predicted a high probability of AI (>0.7). For all previously identified AI candidates, the vast majority of them were predicted with a high Pr(AI) despite missing the threshold.*

**Supplementary Figure 7: Exon density and recombination rate distribution of "AI hits identified by previous studies but missed by *MaLAdapt*", grouped by prediction probability (threshold = 0.6)**

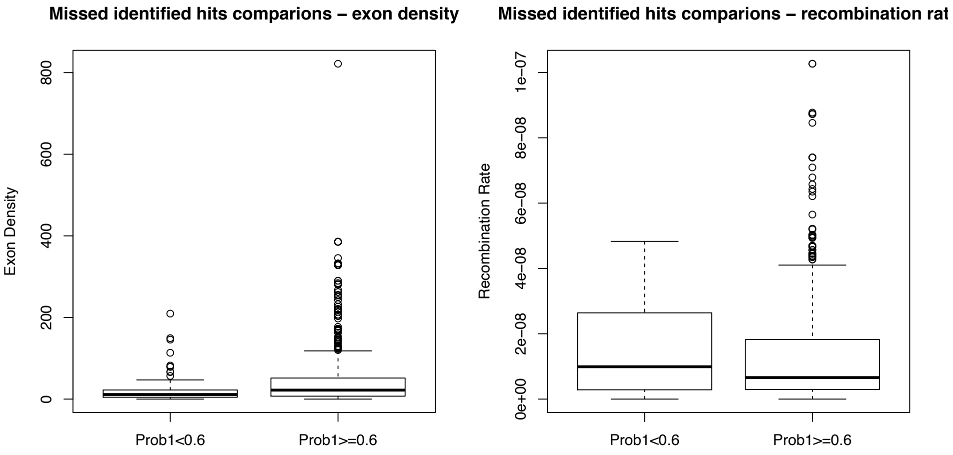

*We examined the distribution of exon density and recombination rates of previously identified AI candidates that are missed by MaLAdapt, and we group such AI candidates as having MaLAdapt Pr(AI) larger than or less than 0.6. We show that the AI candidates that are missed by MaLAdapt completely (Pr(AI)<0.6) have particularly low exon density, indicating that either MaLAdapt is underpowered at detecting AI in regions with fewer exons to produce sufficient informative mutations, or these regions are likely false positives of AI and were detected by other methods due to haplotype structures in these regions from other processes.*

**Supplementary Figure 8: Probability score (-log10(1-PrAI)) distribution of *MaLAdapt* hits, grouped by if a hit has been previously reported**

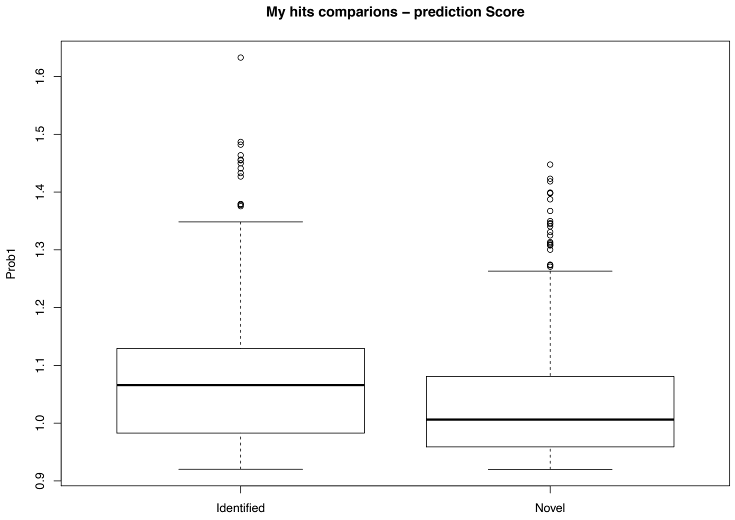

*We compared the AI scores, defined as -log10(1-Pr(AI)) in MaLAdapt-predicted AI candidates, and observed little differentiation in AI score between candidates that are novel discoveries and candidates that have been reported by other studies previously.*

**Supplementary Figure 9: Exon density and recombination rate distribution of *MaLAdapt* hits, grouped by if a hit has been previously reported**

a) Recombination rate – exon density joint distribution

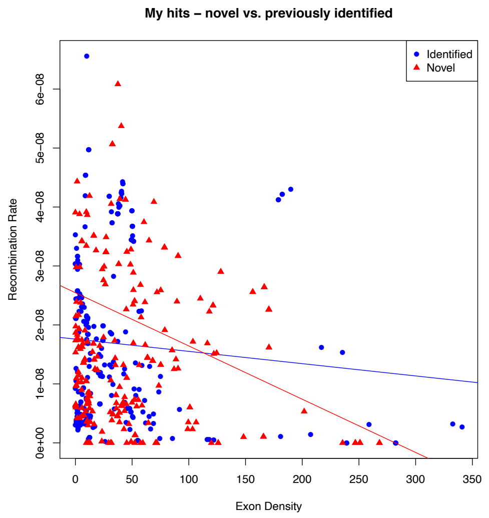

b) Recombination rate and exon density standalone distribution

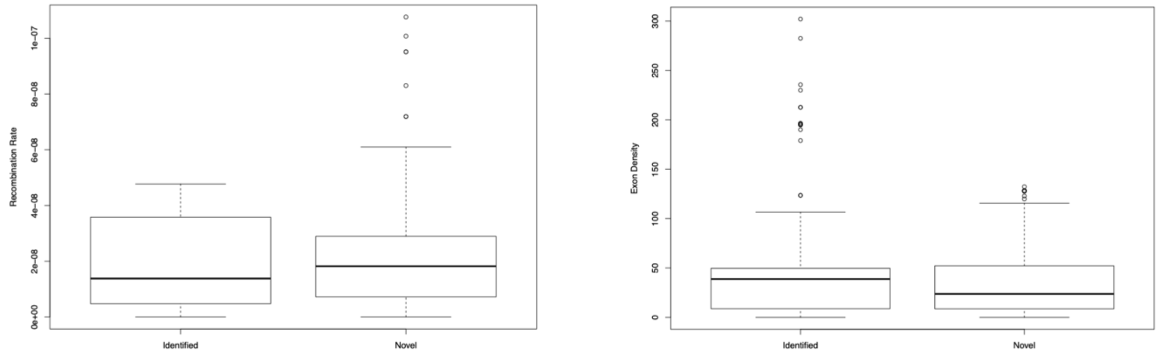

*We compared the exon density and recombination rates distribution in MaLAdapt-predicted AI candidates, and observed little differentiation in the amounts of exons and recombination between candidates that are novel discoveries and candidates that have been reported by other studies previously.*

**Supplementary Figure 10: Prediction score, exon density, and recombination rate joint distributions in *MaLAdapt* identified Neanderthal AI hits in CEU (red) and genome-wide predictions (black)**

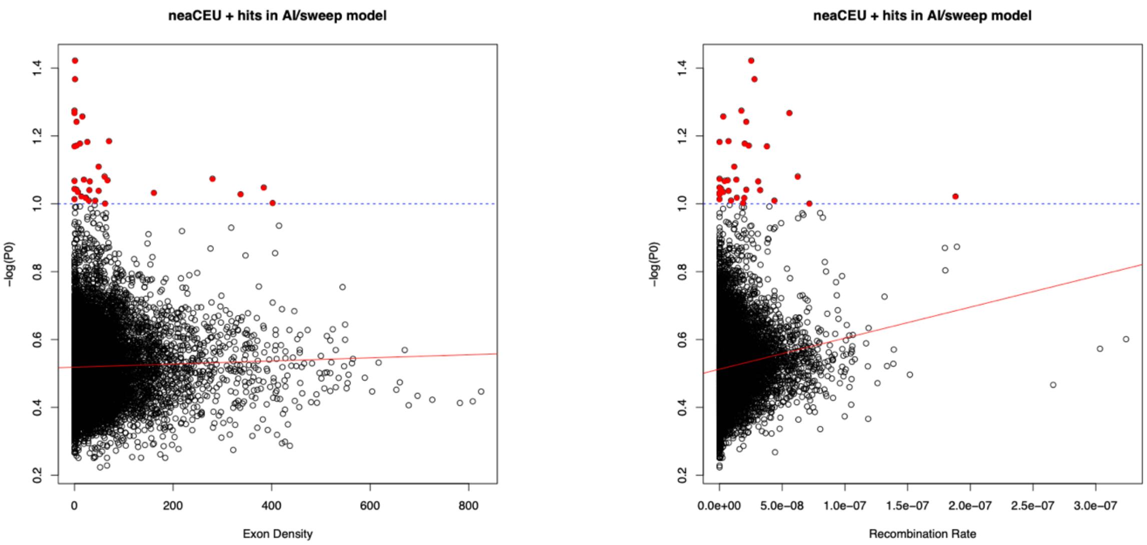

*To examine the potential of MaLAdapt predictions being confounded by heterosis or background selection, we plotted the distribution of Neanderthal AI Scores, defined as -log10(1-Pr(AI)), against the exon density and recombination rates in sliding windows across the genome of CEU. We highlight the windows where AI are predicted, and show that these windows are not outliers in terms of either exon density or recombination rates.*

**Supplementary Figure 11: Exon density and recombination rate distributions in Neanderthal adaptive introgression in CEU predicted by *MaLAdapt*, *genomatnn*, and *VolcanoFinder***

a) Exon density and recombination rate joint distribution

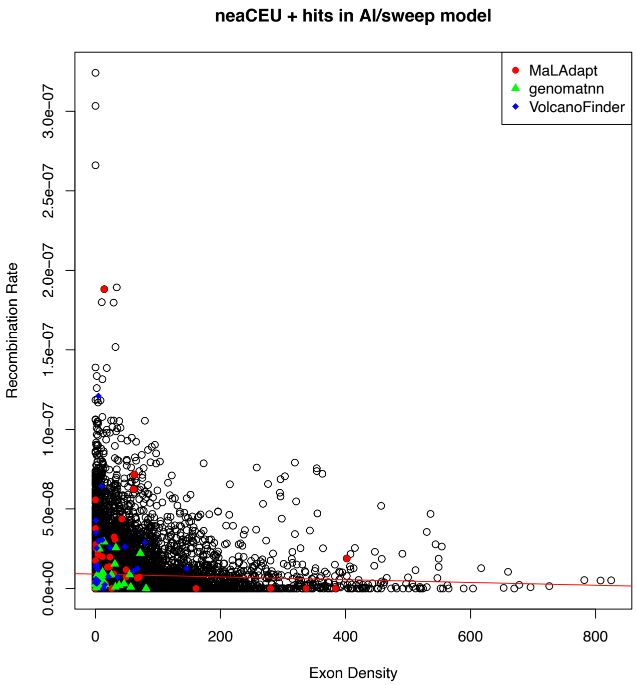

b) Exon density distribution

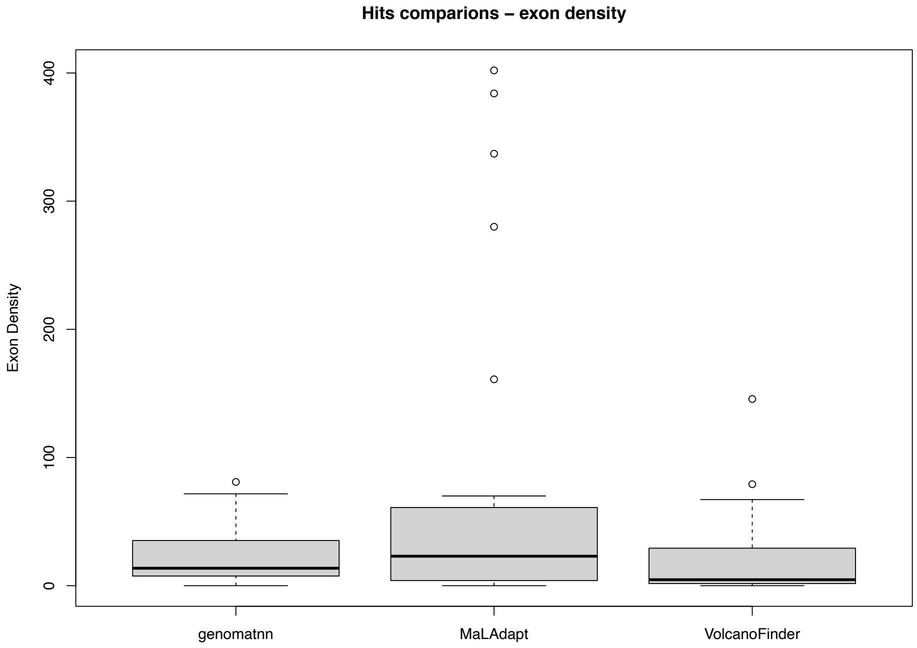

c) Recombination rate distribution

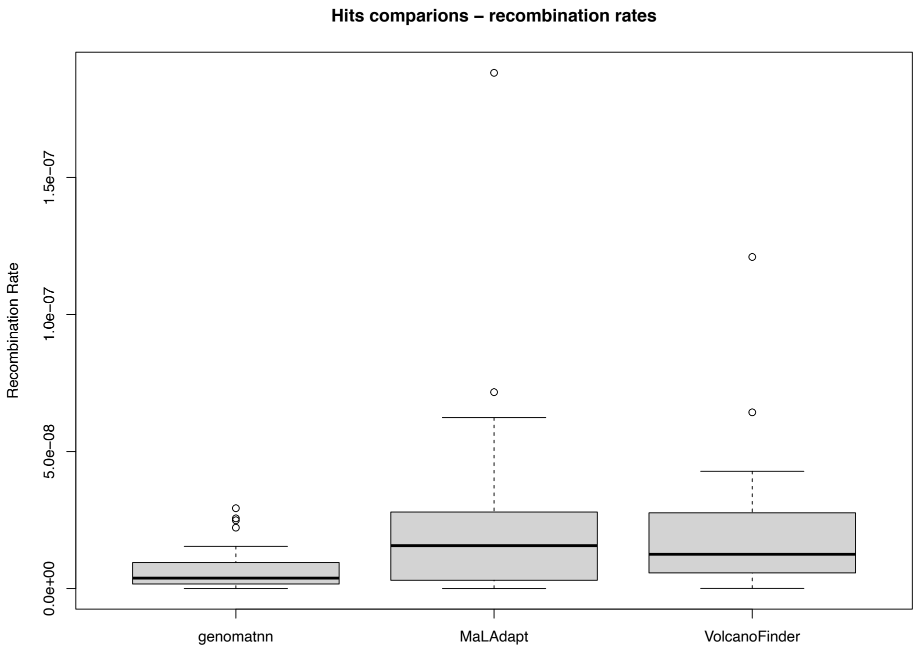

*We compared the distribution of exon density, recombination rates, and their joint distribution for Neanderthal AI regions in CEU predicted by MaLAdapt, genotmatnn and VolcanoFinder. We show that compared to the genomic background, MaLAdapt and VolcanoFinder AI regions do not have unusual exon density or recombination rates, whereas genomatnn AI regions are enriched in regions with low recombination rates, which could potentially lead to haplotype structures from non-AI processes.*

**Supplementary Figure 12: *MaLAdapt* prediction probability at AI hits reported by *genomatnn* and *VolcanoFinder* methods**

a) *genomatnn* predictions

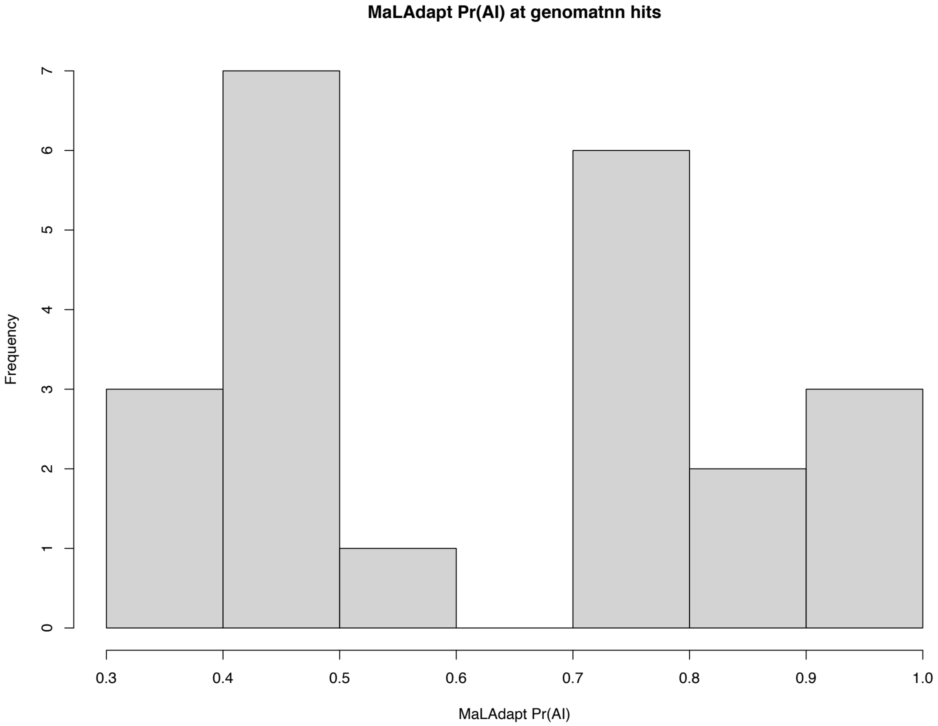

b) *VolcanoFinder* predictions

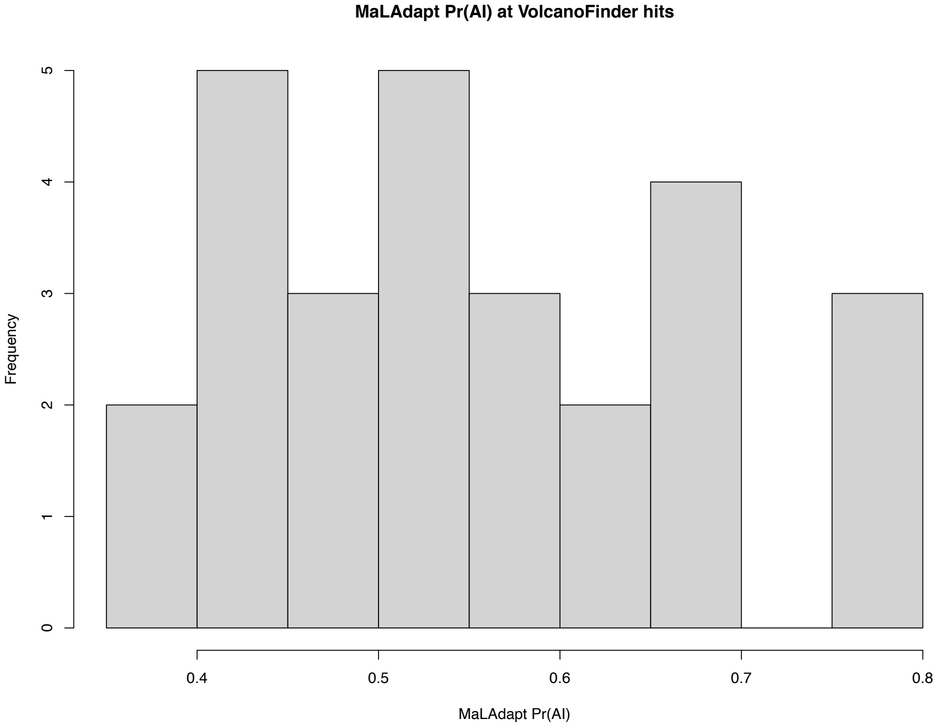

*To understand the underlying causes of why MaLAdapt missed certain AI regions predicted by genomatnn or VolcanoFinder, we extracted the Pr(AI) of MaLAdapt in all AI regions from the other two methods. We show that for genomatnn, MaLAdapt predicted a bimodal distribution in those*

*regions, in which nearly half of the genomatnn regions received low Pr(AI). And in VolcanoFinder regions, MaLAdapt's predictions are evenly distributed across different probabilities, although none passed the MaLAdapt AI threshold (0.9).*

**Supplementary Figure 13: Exon density and recombination rate in AI hits reported by *genomatnn*, annotated manually as True Positive, False Positive and uncertain given their haplotype structures (Supplementary Figure 18)**

a) Exon density

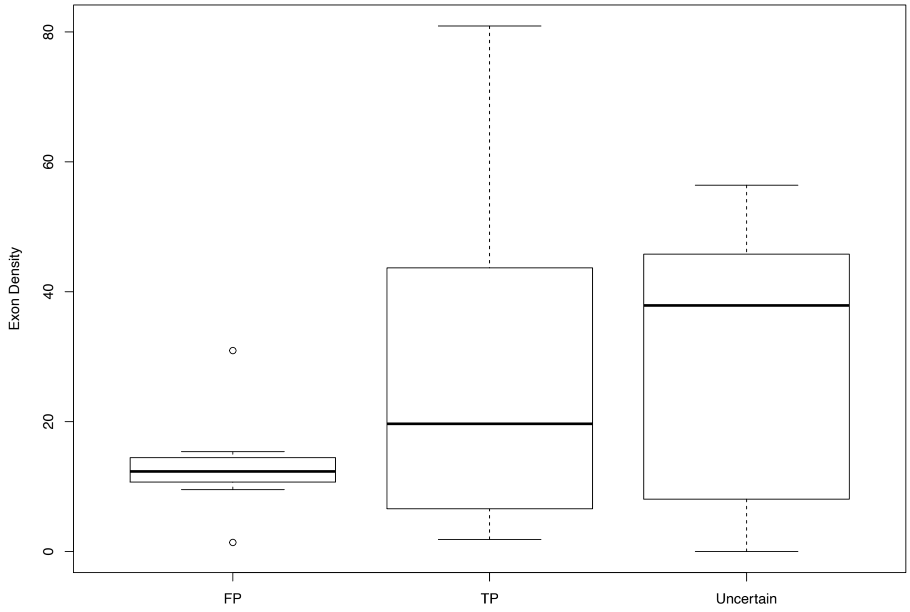

b) Recombination rate

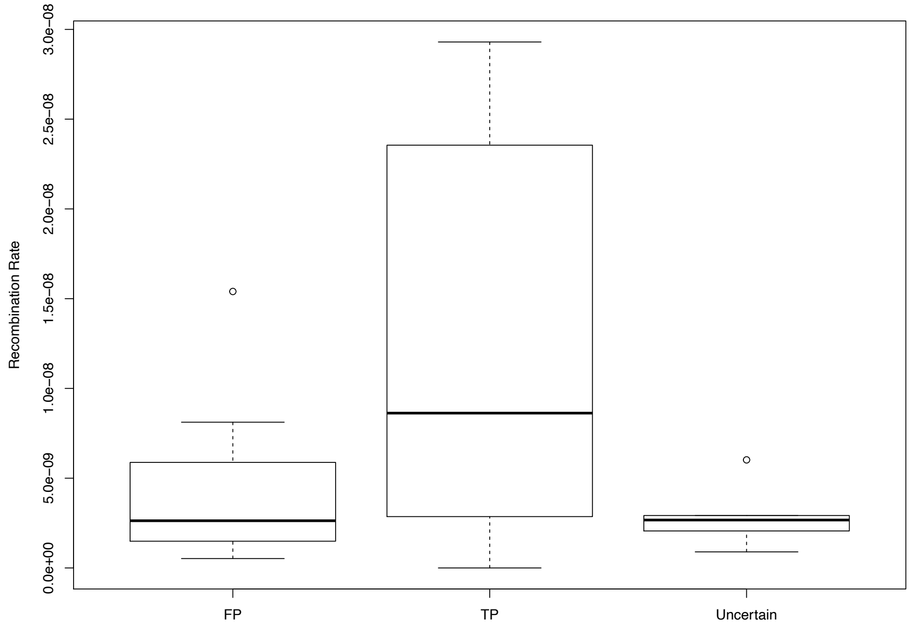

*We examined closely into the genomatnn regions to understand the bimodal Pr(AI) distribution we observed in Supplementary Figure 12. We manually annotated the AI regions as "True Positive (TP)", "False Positive (FP)", and "uncertain" given their haplotype structures (Supplementary Figure 18), and show that the likely False Positive predictions from genomatnn are regions that have especially low recombination rates and low exon density, implying that these are false positives caused by haplotype structures due to non-AI processes.*

**Supplementary Figure 14: Distribution of *MaLAdapt* AI prediction probability genomic background, human genes, VIP genes, non-VIP control genes, and VIP-AI candidates.**

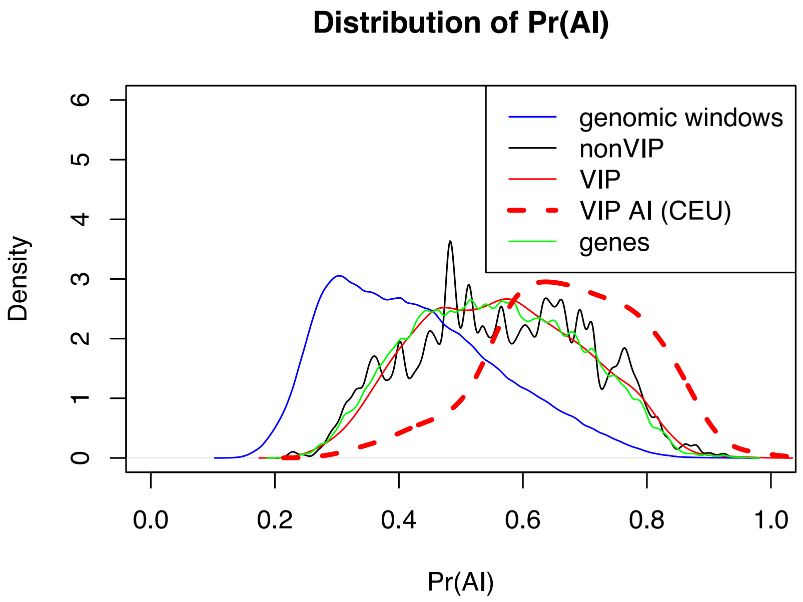

*We compare the distribution of Pr(AI) in the following 5 datasets: 1) VIP genes (Enard and Petrov 2018; 3963 genes in total); 2) 1000 sets of non-VIP genes as controls (matched size to the VIP set); 3) one set of randomly sampled human genes (regardless of VIP or non-VIP; matched size to the VIP set); 4) one set of randomly sampled genomic 50kb windows (matched size to the VIP set); 5) VIP genes that are also Neanderthal AI candidates in CEU, suggested from Enard and Petrov 2018. We show that although there is a slight increase in Pr(AI) in VIP genes compared to nonVIP genes, the difference is not significant (p-value = 0.459, two-sided Wilcoxon rank sum test). However, the VIP genes that are also AI candidates have a significant increase in Pr(AI) compared to either VIP or nonVIP sets (Wilcoxon p-value < 2.2e-16 for both comparisons).*

**Supplementary Figure 15: p-values of Wilcoxon rank sum test for the distribution between the VIP gene set and 1000 nonVIP gene sets**

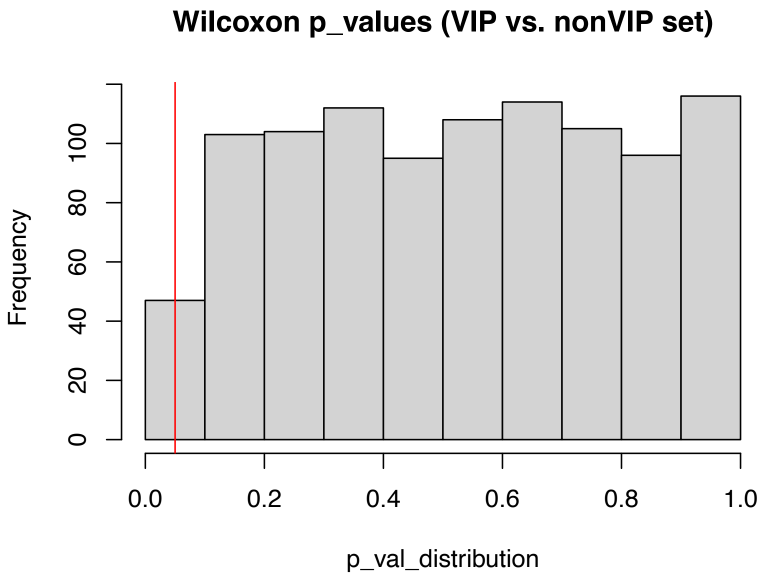

*We conducted two-sided Wilcoxon rank sum test between the Pr(AI) of VIP gene set and each of the 1000 non-VIP gene sets, and plotted the distribution of 1000 p-values using alternative hypothesis as the true location shift is not equal to 0. The red solid line indicates a reference p-value being 0.05. We show that the p-values are uniformly distributed, indicating that there is little difference in probability being AI between VIP genes and non-VIP genes.*

**Supplementary Figure 16: Neanderthal adaptive introgression in 1000 Genomes Populations**

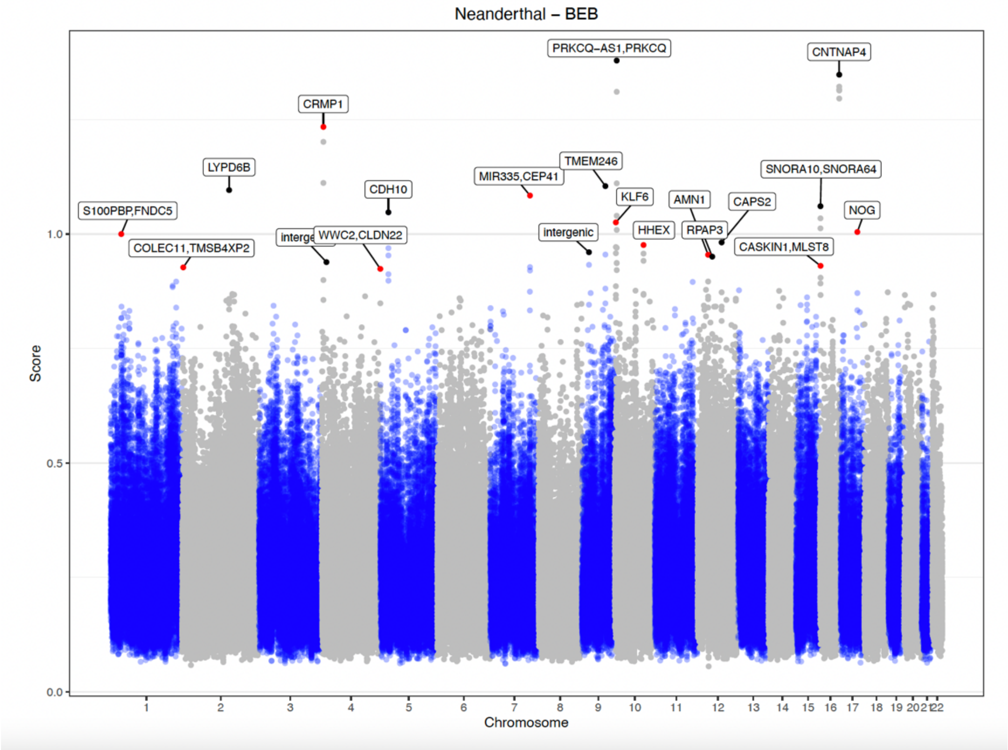

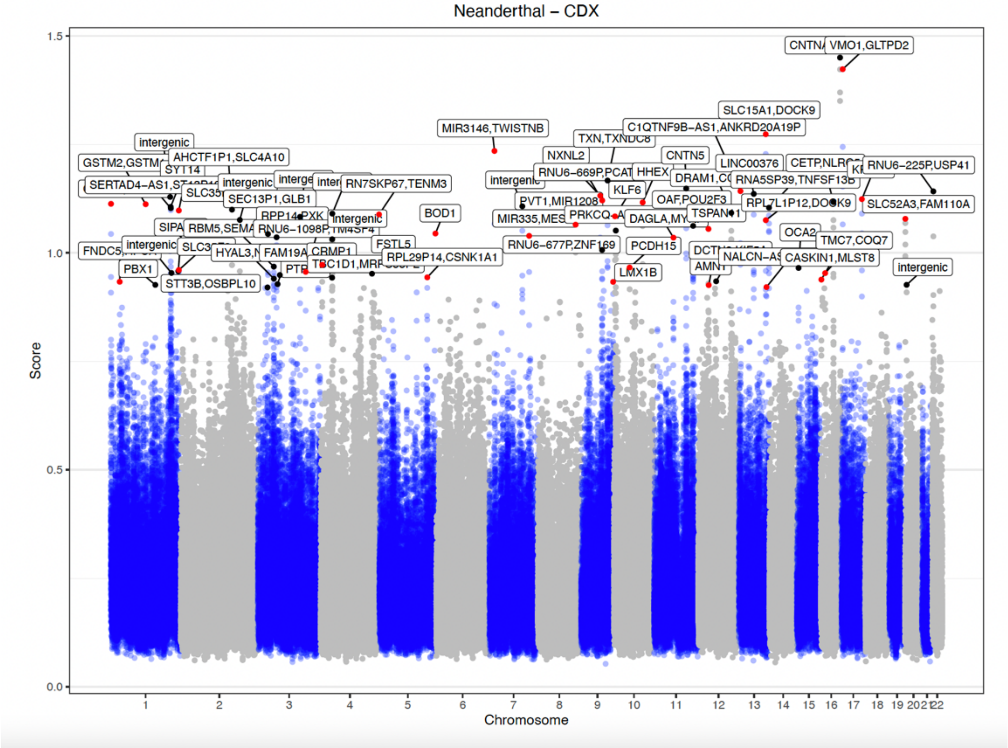

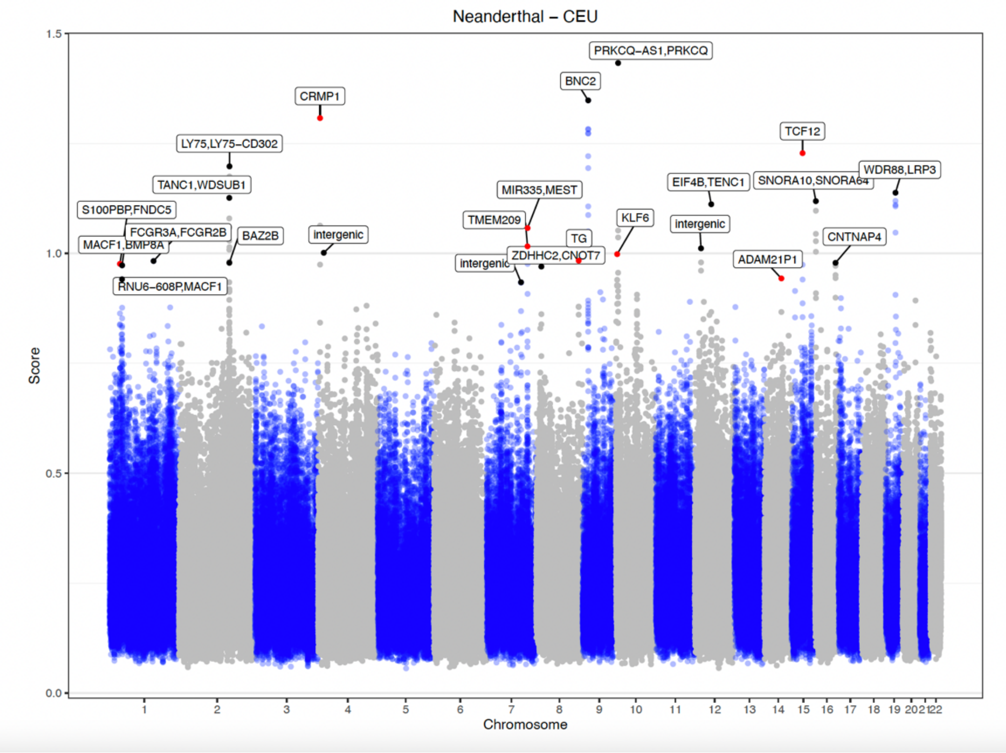

Neanderthal – CEU
Score
Chromosome
PRKCQ–AS1,PRKCQ
BNC2
CRMP1
LY75,LY75–CD302
TCF12
WDR88,LRP3
SNORA10,SNORA64
EIF4B,TENC1
TANC1,WDSUB1
MIR335,MEST
S100PBP,FNDC5
TMEM209
KLF6
intergenic
FCGR3A,FCGR2B
BAZ2B
intergenic
TG
CNTNAP4
MACF1,BMP8A
ZDHHC2,CNOT7
ADAM21P1
intergenic
RNU6–608P,MACF1

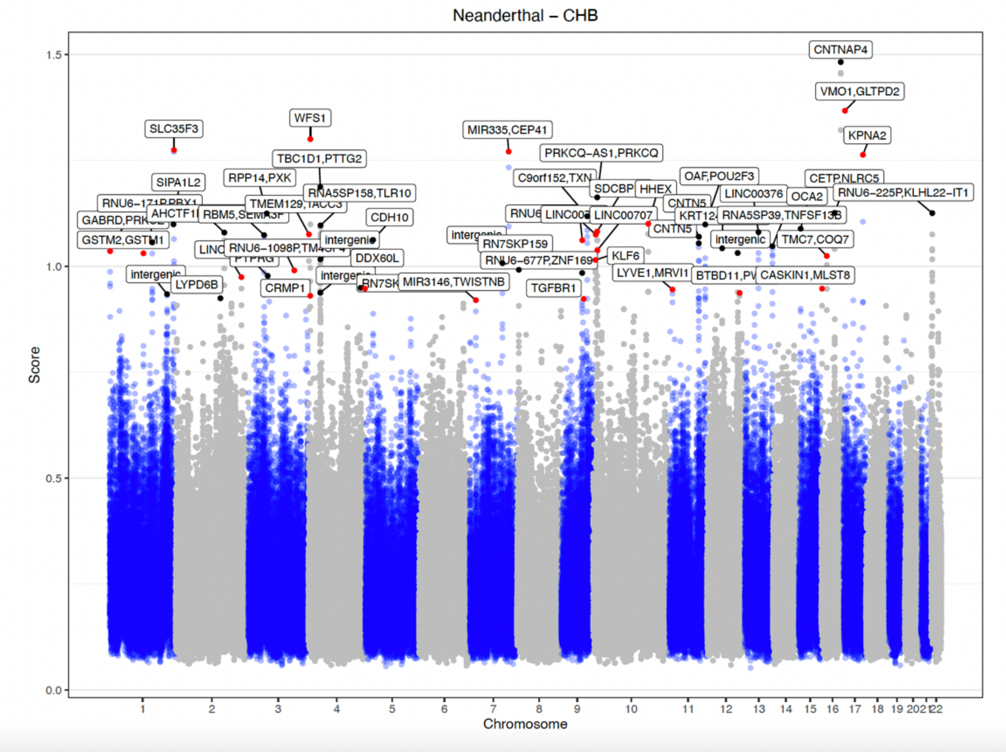

Neanderthal – CHB
Score
Chromosome
CNTNAP4
VMO1,GLTPD2
WFS1
SLC35F3
MIR335,CEP41
KPNA2
TBC1D1,PTTG2
PRKCQ–AS1,PRKCQ
SIPA1L2
RPP14,PXK
C9orf152,TXN
OAF,POU2F3
CETP,NLRC5
RNA5SP158,TLR10
SDCBP
HHEX
LINC00376
OCA2
RNU6–225P,KLHL22–IT1
CDH10
LINC00707
CNTN5
RNA5SP39,TNFSF13B
intergenic
intergenic
TMC7,COQ7
RN7SKP159
DDX60L
KLF6
intergenic
LYPD6B
CRMP1
MIR3146,TWISTNB
TGFBR1
CASKIN1,MLST8

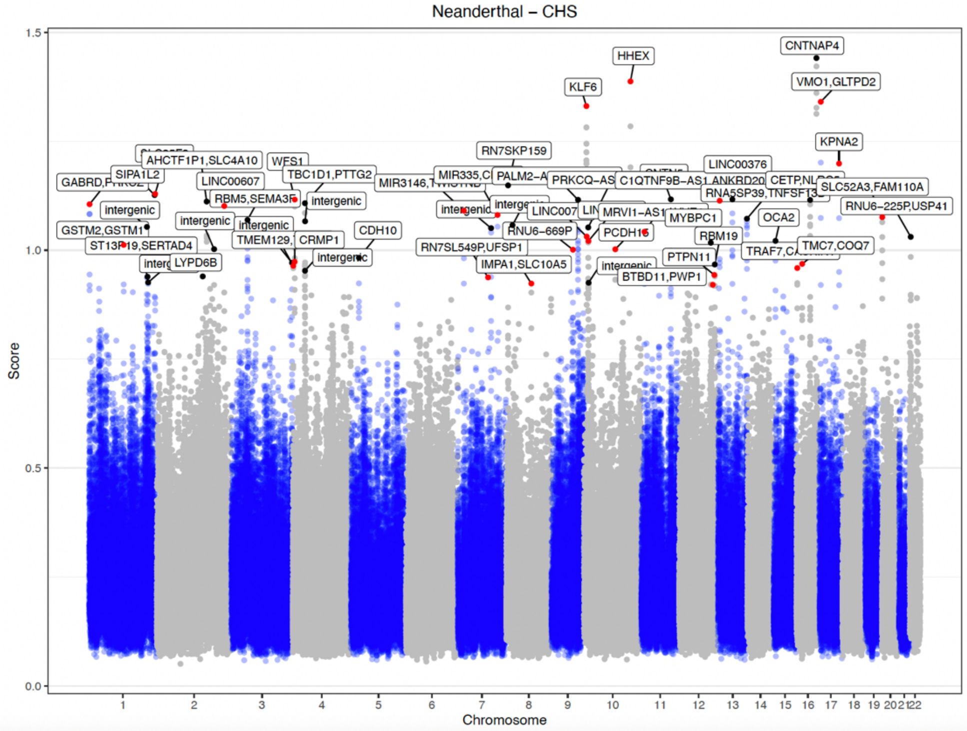
Neanderthal – CHS
Score
Chromosome
CNTNAP4
HHEX
VMO1,GLTPD2
KLF6
KPNA2
RN7SKP159
LINC00376
AHCTF1P1,SLC4A10
WFS1
SIPA1L2
GABRD
LINC00607
TBC1D1,PTTG2
MIR3146
PALM2–A
PRKCQ–AS
C1QTNF9B–AS1,ANKRD20
SLC52A3,FAM110A
RNA5SP39,TNFSF13
RNU6–225P,USP41
RBM5,SEMA3F
intergenic
GSTM2,GSTM1
ST13P19,SERTAD4
TMEM129
CRMP1
CDH10
LYPD6B
RN7SL549P,UFSP1
RNU6–669P
MRVI1–AS1
MYBPC1
OCA2
PCDH15
RBM19
TRAF7
TMC7,COQ7
PTPN11
IMPA1,SLC10A5
BTBD11,PWP1
0.0
0.5
1.0
1.5

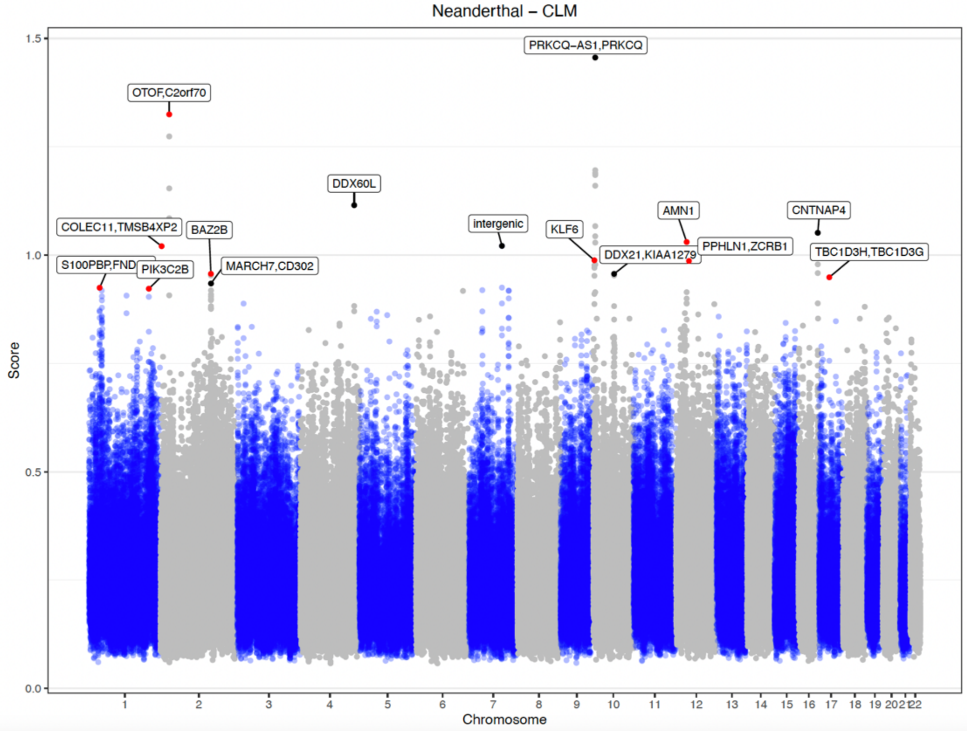
Neanderthal – CLM
Score
Chromosome
PRKCQ–AS1,PRKCQ
OTOF,C2orf70
DDX60L
COLEC11,TMSB4XP2
BAZ2B
intergenic
KLF6
AMN1
CNTNAP4
DDX21,KIAA1279
PPHLN1,ZCRB1
TBC1D3H,TBC1D3G
PIK3C2B
MARCH7,CD302
0.0
0.5
1.0
1.5

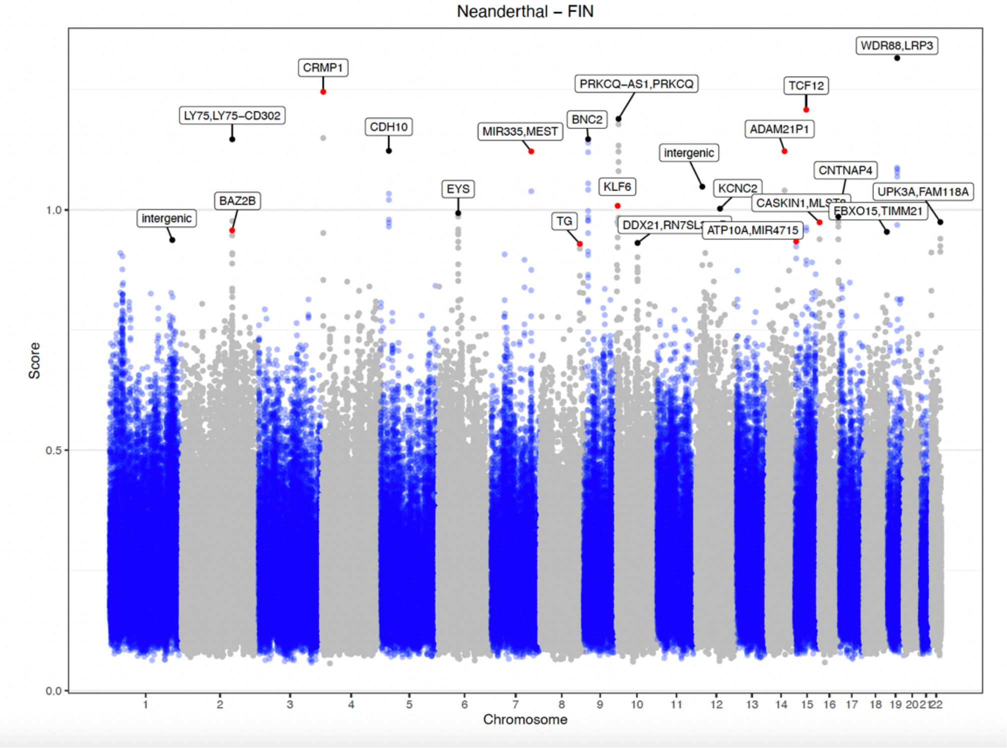
Neanderthal – FIN
Score
Chromosome
0.0
0.5
1.0
1 2 3 4 5 6 7 8 9 10 11 12 13 14 15 16 17 18 19 20 21 22
WDR88,LRP3
CRMP1
PRKCQ−AS1,PRKCQ
TCF12
LY75,LY75−CD302
CDH10
BNC2
MIR335,MEST
ADAM21P1
intergenic
CNTNAP4
EYS
KLF6
KCNC2
UPK3A,FAM118A
BAZ2B
intergenic
TG
FBXO15,TIMM21
ATP10A,MIR4715

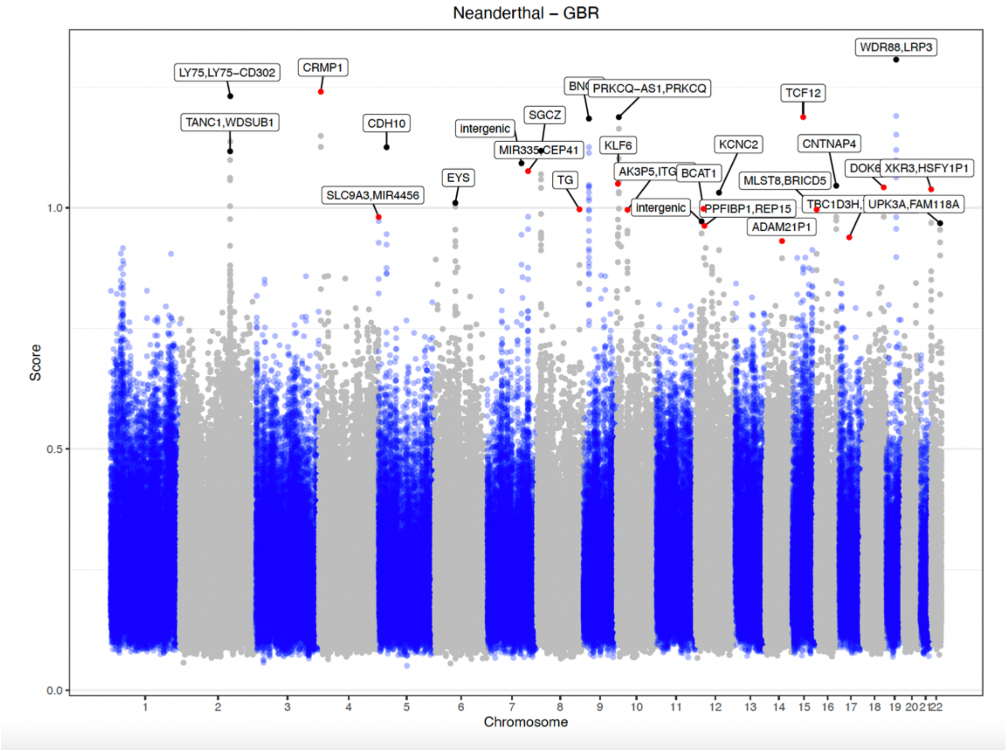
Neanderthal – GBR
Score
Chromosome
0.0
0.5
1.0
1 2 3 4 5 6 7 8 9 10 11 12 13 14 15 16 17 18 19 20 21 22
WDR88,LRP3
LY75,LY75−CD302
CRMP1
PRKCQ−AS1,PRKCQ
TCF12
TANC1,WDSUB1
CDH10
intergenic
SGCZ
KLF6
KCNC2
CNTNAP4
EYS
TG
BCAT1
XKR3,HSFY1P1
MLST8,BRICD5
SLC9A3,MIR4456
intergenic
PPFIBP1,REP15
UPK3A,FAM118A
ADAM21P1

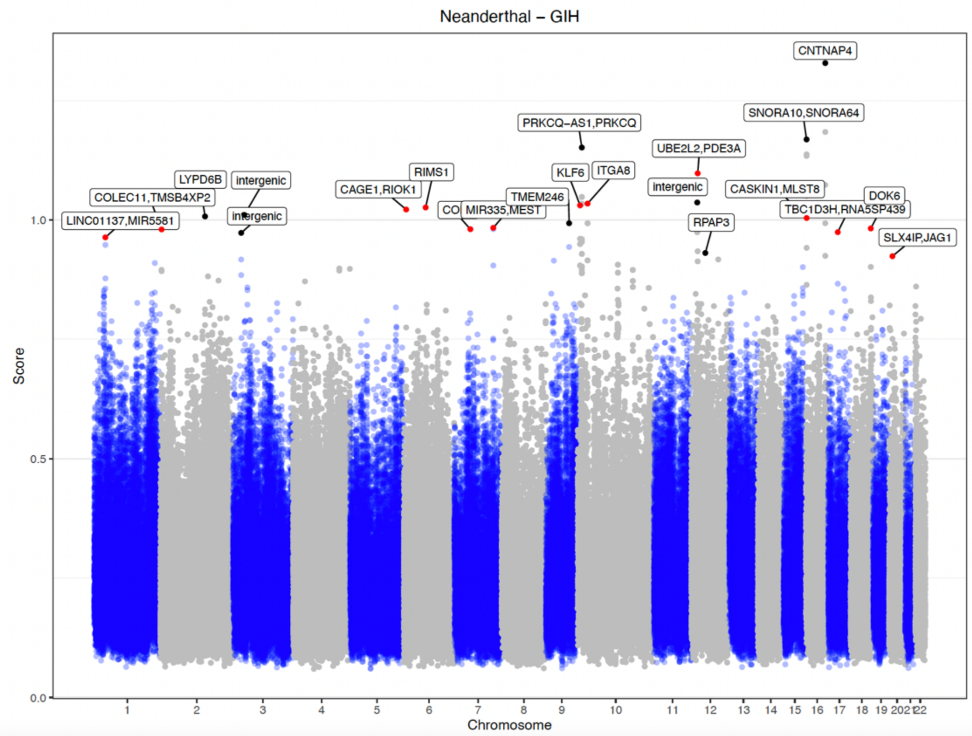
Neanderthal – GIH
CNTNAP4
SNORA10,SNORA64
PRKCQ-AS1,PRKCQ
UBE2L2,PDE3A
LYPD6B
intergenic
RIMS1
KLF6
ITGA8
COLEC11,TMSB4XP2
CAGE1,RIOK1
intergenic
CASKIN1,MLST8
DOK6
TMEM246
TBC1D3H,RNA5SP439
LINC01137,MIR5581
intergenic
MIR335,MEST
RPAP3
SLX4IP,JAG1
Score
Chromosome

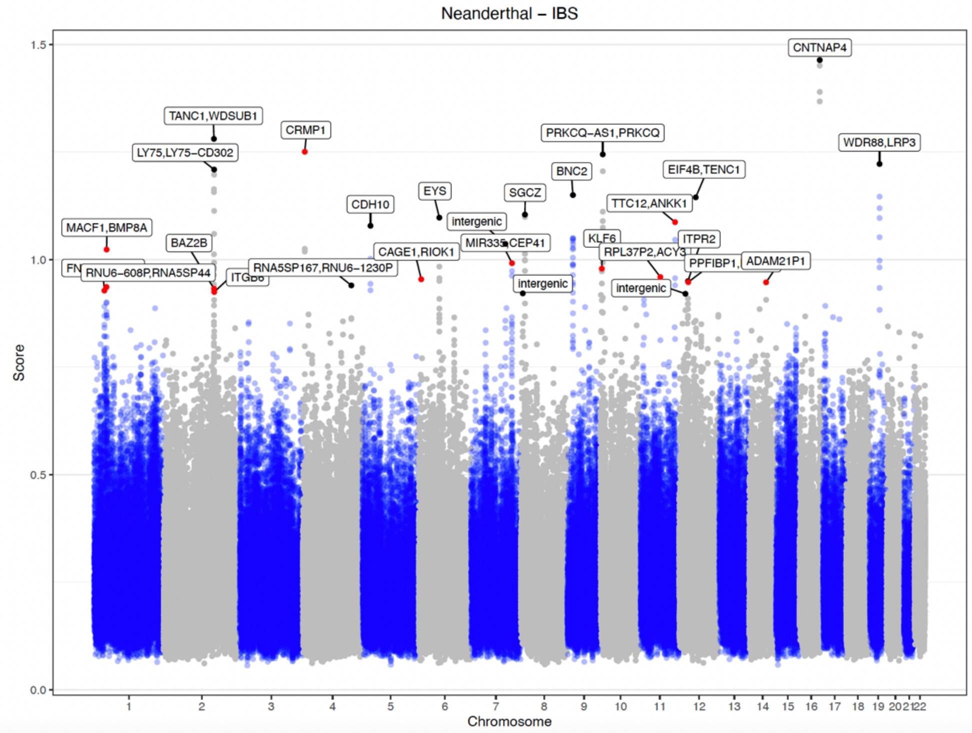
Neanderthal – IBS
CNTNAP4
TANC1,WDSUB1
CRMP1
PRKCQ-AS1,PRKCQ
LY75,LY75-CD302
WDR88,LRP3
BNC2
EIF4B,TENC1
EYS
SGCZ
CDH10
TTC12,ANKK1
MACF1,BMP8A
intergenic
BAZ2B
KLF6
ITPR2
CAGE1,RIOK1
MIR335,CEP41
RNU6-608P,RNA5SP44
RNA5SP167,RNU6-1230P
ADAM21P1
intergenic
intergenic
Score
Chromosome

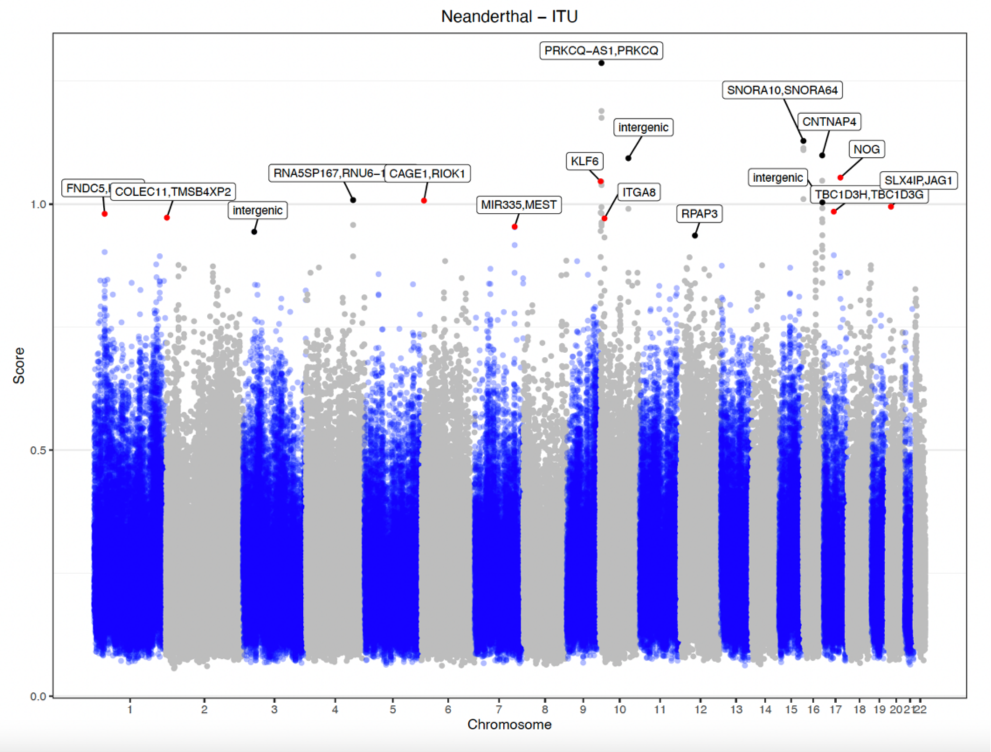
Neanderthal – ITU
PRKCQ–AS1,PRKCQ
SNORA10,SNORA64
CNTNAP4
intergenic
NOG
KLF6
RNA5SP167,RNU6–1
CAGE1,RIOK1
intergenic
SLX4IP,JAG1
COLEC11,TMSB4XP2
ITGA8
TBC1D3H,TBC1D3G
intergenic
MIR335,MEST
RPAP3
Score
Chromosome

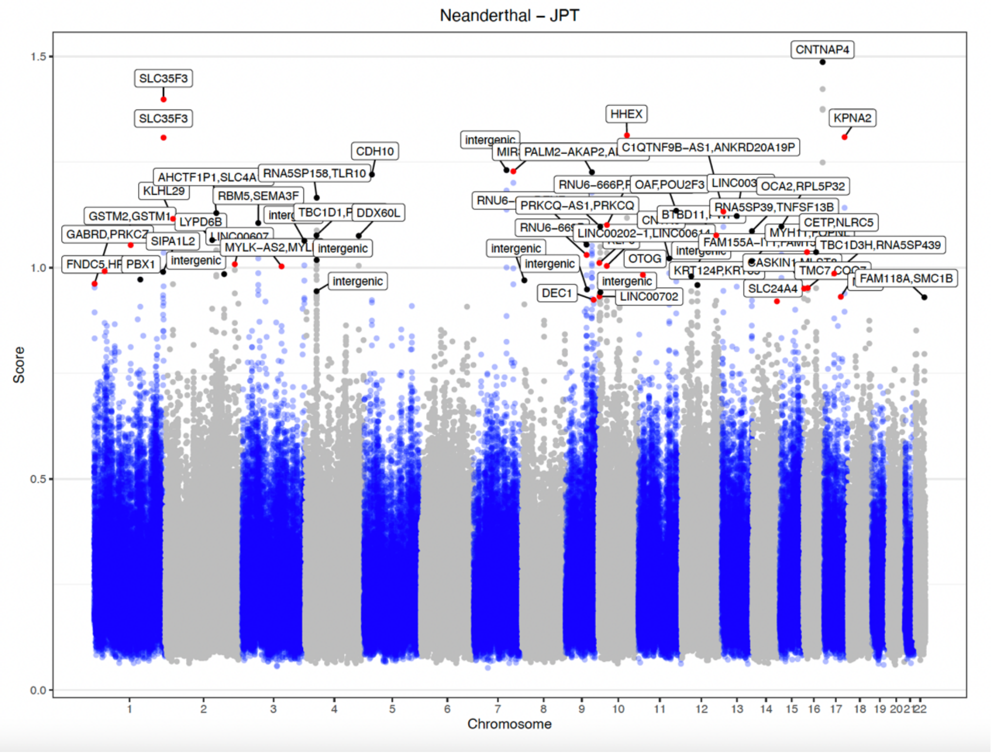
Neanderthal – JPT
CNTNAP4
SLC35F3
SLC35F3
HHEX
KPNA2
CDH10
C1QTNF9B–AS1,ANKRD20A19P
AHCTF1P1,SLC4A
RNA5SP158,TLR10
KLHL29
RBM5,SEMA3F
OAF,POU2F3
OCA2,RPL5P32
GSTM2,GSTM1
LYPD6B
DDX60L
PRKCQ–AS1,PRKCQ
RNA5SP39,TNFSF13B
CETP,NLRC5
GABRD,PRKCZ
SIPA1L2
intergenic
TBC1D3H,RNA5SP439
intergenic
intergenic
OTOG
FNDC5,HP
PBX1
intergenic
intergenic
FAM118A,SMC1B
DEC1
LINC00702
SLC24A4
Score
Chromosome

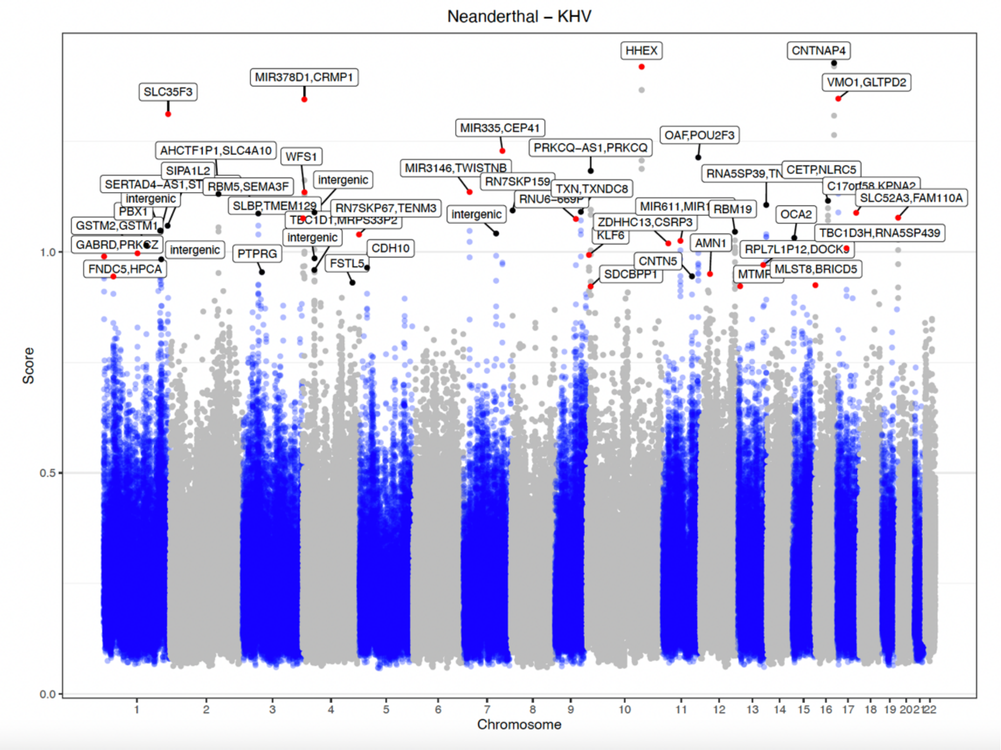
Neanderthal – KHV
Score
Chromosome
HHEX
CNTNAP4
VMO1,GLTPD2
MIR378D1,CRMP1
SLC35F3
MIR335,CEP41
OAF,POU2F3
PRKCQ–AS1,PRKCQ
AHCTF1P1,SLC4A10
WFS1
MIR3146,TWISTNB
SIPA1L2
intergenic
RN7SKP159
TXN,TXNDC8
CETP,NLRC5
SLC52A3,FAM110A
RBM5,SEMA3F
RBM19
OCA2
TBC1D3H,RNA5SP439
ZDHHC13,CSRP3
KLF6
AMN1
CNTN5
SDCBPP1
MLST8,BRICD5
PBX1
GSTM2,GSTM1
GABRD,PRKCZ
FNDC5,HPCA
PTPRG
CDH10
FSTL5
RN7SKP67,TENM3
SLBP,TMEM129

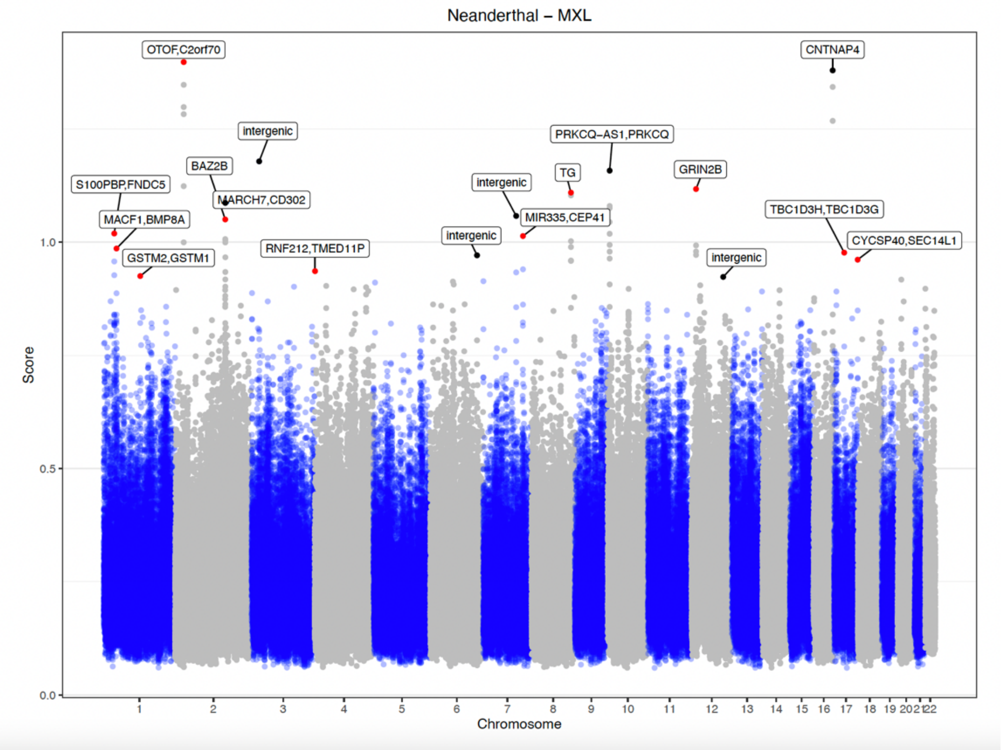
Neanderthal – MXL
Score
Chromosome
OTOF,C2orf70
CNTNAP4
intergenic
PRKCQ–AS1,PRKCQ
BAZ2B
GRIN2B
TG
S100PBP,FNDC5
MARCH7,CD302
MIR335,CEP41
TBC1D3H,TBC1D3G
MACF1,BMP8A
RNF212,TMED11P
CYCSP40,SEC14L1
GSTM2,GSTM1

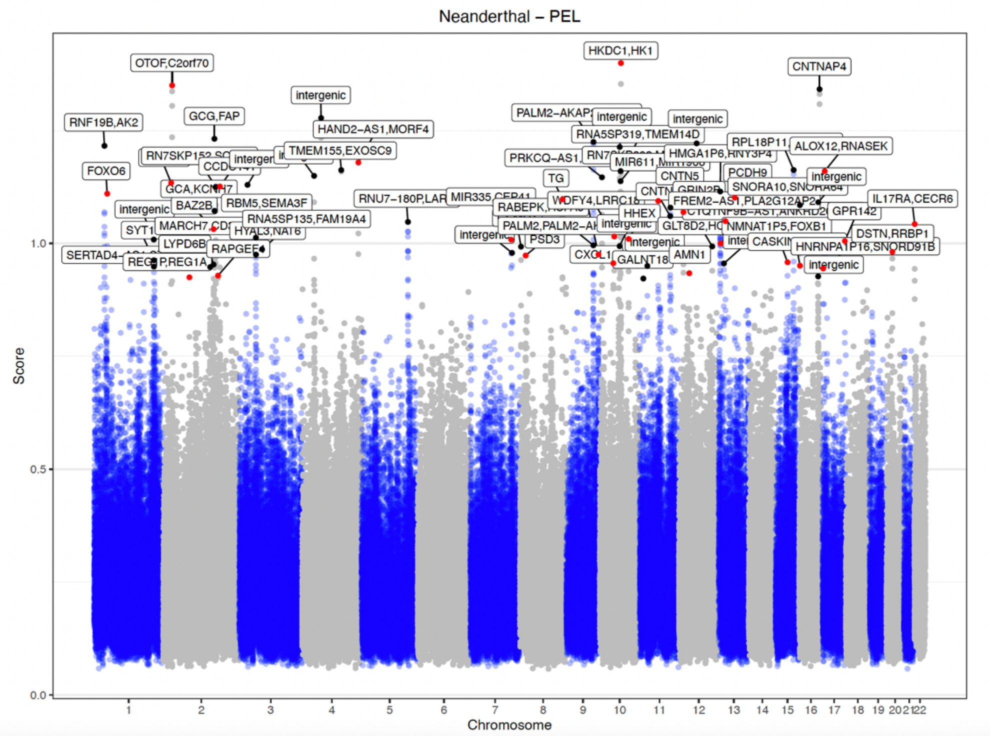
Neanderthal – PEL
Score
Chromosome
OTOF,C2orf70
HKDC1,HK1
CNTNAP4
RNF19B,AK2
GCG,FAP
intergenic
HAND2−AS1,MORF4
TMEM155,EXOSC9
FOXO6
RNU7−180P,LAR
RNA5SP135,FAM19A4
RBM5,SEMA3F
PRKCQ−AS1
RNA5SP319,TMEM14D
ALOX12,RNASEK
IL17RA,CECR6
GPR142
DSTN,RRBP1
HNRNPA1P16,SNORD91B
PSD3
AMN1
TG
CNTN5
PCDH9
HHEX
0.0
0.5
1.0

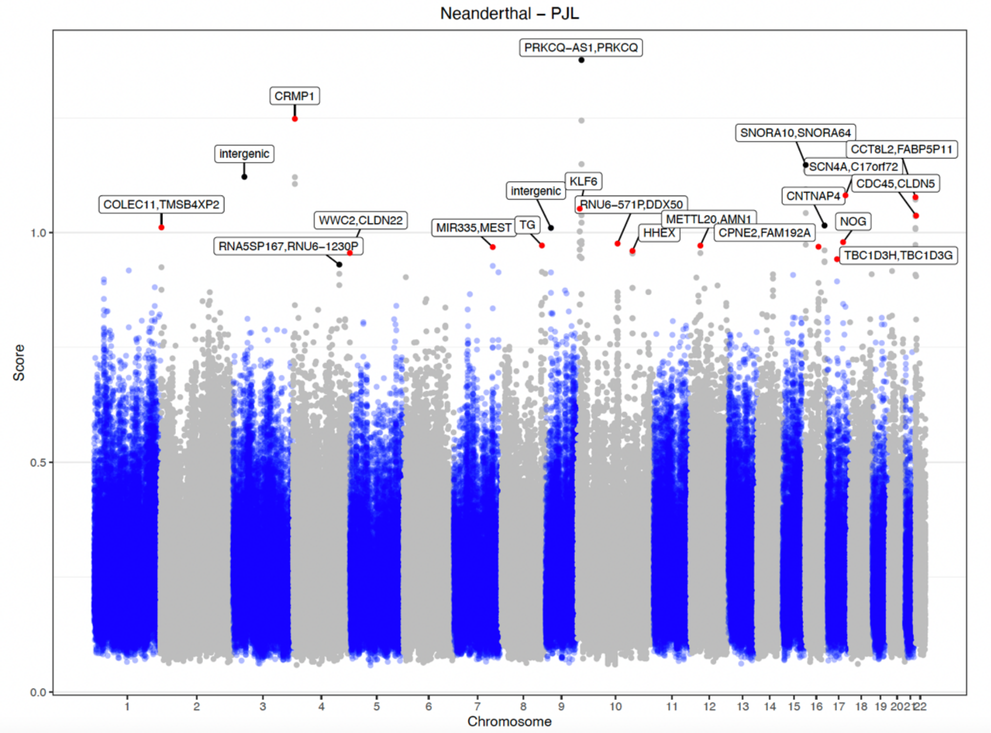
Neanderthal – PJL
Score
Chromosome
PRKCQ−AS1,PRKCQ
CRMP1
intergenic
COLEC11,TMSB4XP2
WWC2,CLDN22
RNA5SP167,RNU6−1230P
MIR335,MEST
TG
KLF6
RNU6−571P,DDX50
HHEX
METTL20,AMN1
CPNE2,FAM192A
SNORA10,SNORA64
CCT8L2,FABP5P11
SCN4A,C17orf72
CDC45,CLDN5
CNTNAP4
NOG
TBC1D3H,TBC1D3G
0.0
0.5
1.0

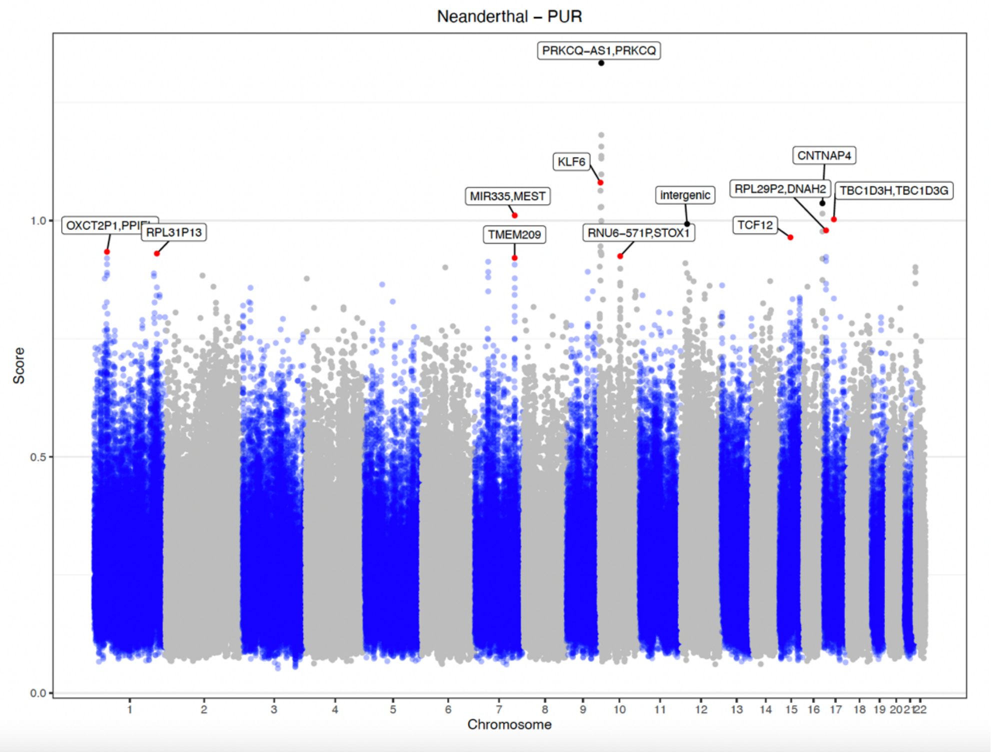

Neanderthal – PUR
PRKCQ–AS1,PRKCQ
KLF6
CNTNAP4
MIR335,MEST
intergenic
RPL29P2,DNAH2
TBC1D3H,TBC1D3G
RPL31P13
TMEM209
RNU6–571P,STOX1
TCF12
Score
Chromosome

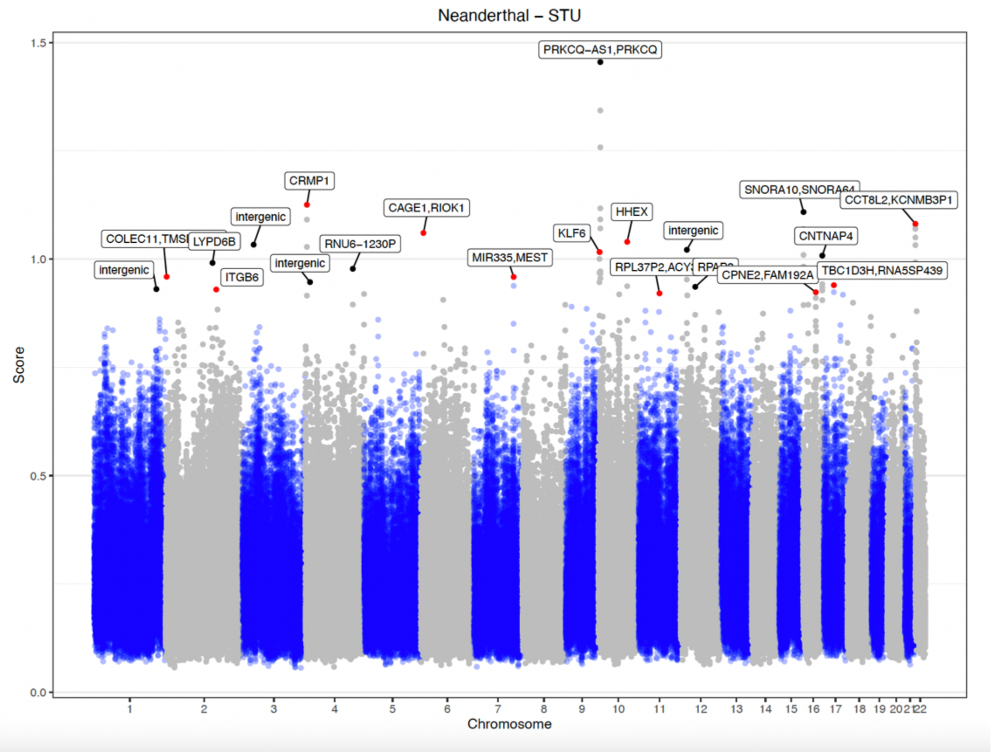

Neanderthal – STU
PRKCQ–AS1,PRKCQ
CRMP1
intergenic
CAGE1,RIOK1
CCT8L2,KCNMB3P1
LYPD6B
HHEX
KLF6
intergenic
CNTNAP4
intergenic
ITGB6
intergenic
RNU6–1230P
MIR335,MEST
CPNE2,FAM192A
TBC1D3H,RNA5SP439
Score
Chromosome

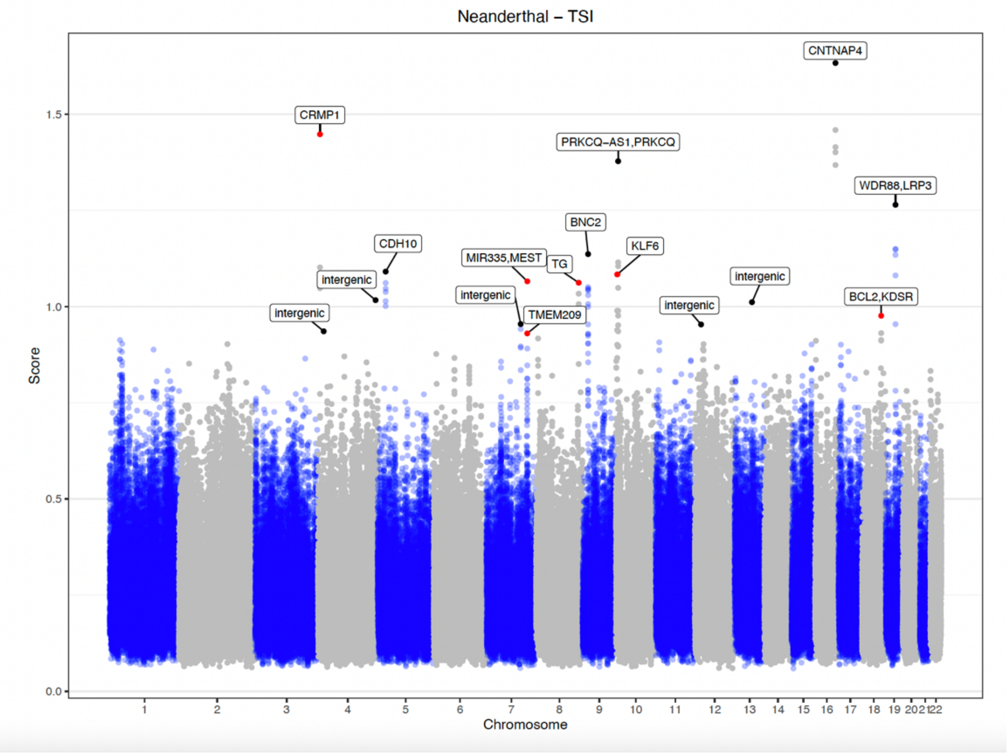

*We applied MaLAdapt to predict AI in overlapping 50kb windows (step size 10kb) along the genome of non-African populations of the 1000 Genomes data. Here we show the AI prediction results of 19 non-African populations, using African (YRI) as non-introgressed outgroup and Altai Neanderthal as the introgression donor. The Y-axis shows the AI prediction score, which equals the -Log10 transformed value of [1-Pr(AI)]. Each dot in the plot represents a 50kb window. The windows that did not reach the MaLAdapt AI threshold are colored in blue or gray depending on the chromosomes. The windows detected as AI are colored in black if they have been reported by previous studies before, or in red if they are novel findings from this study. The labels highlight the gene names that overlap with the AI windows.*

**Supplementary Figure 17: Denisovan adaptive introgression in 1000 Genomes Populations**

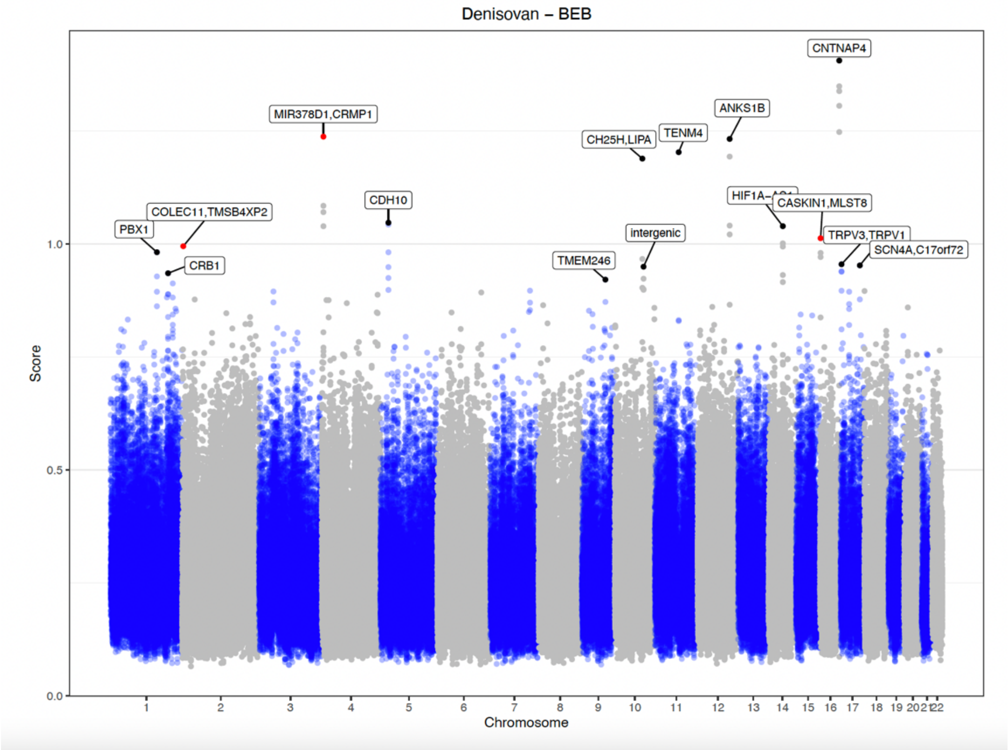

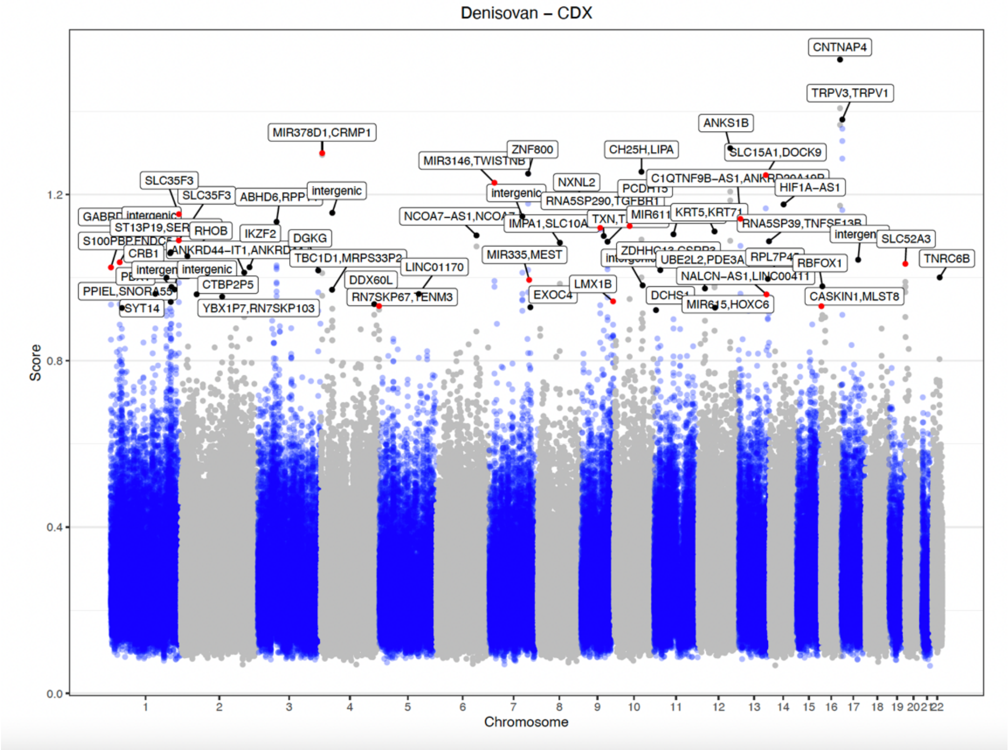

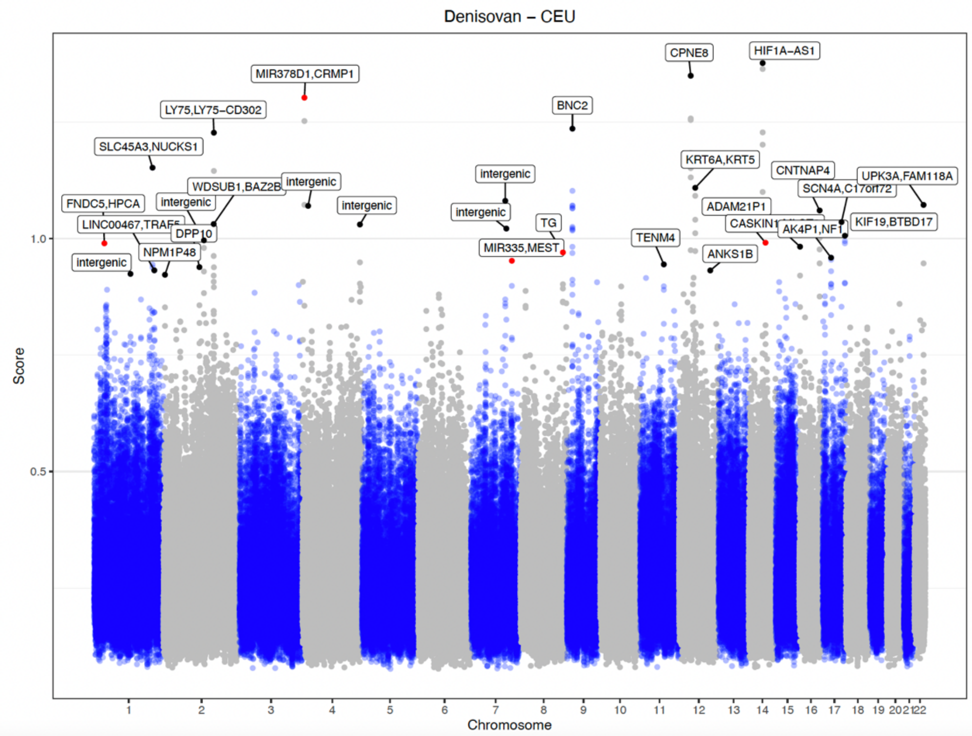
Denisovan – CEU
Score
Chromosome
1.0
0.5
CPNE8
HIF1A−AS1
MIR378D1,CRMP1
LY75,LY75−CD302
BNC2
SLC45A3,NUCKS1
KRT6A,KRT5
CNTNAP4
UPK3A,FAM118A
WDSUB1,BAZ2B
intergenic
FNDC5,HPCA
ADAM21P1
KIF19,BTBD17
TG
MIR335,MEST
TENM4
ANKS1B
NPM1P48
DPP10

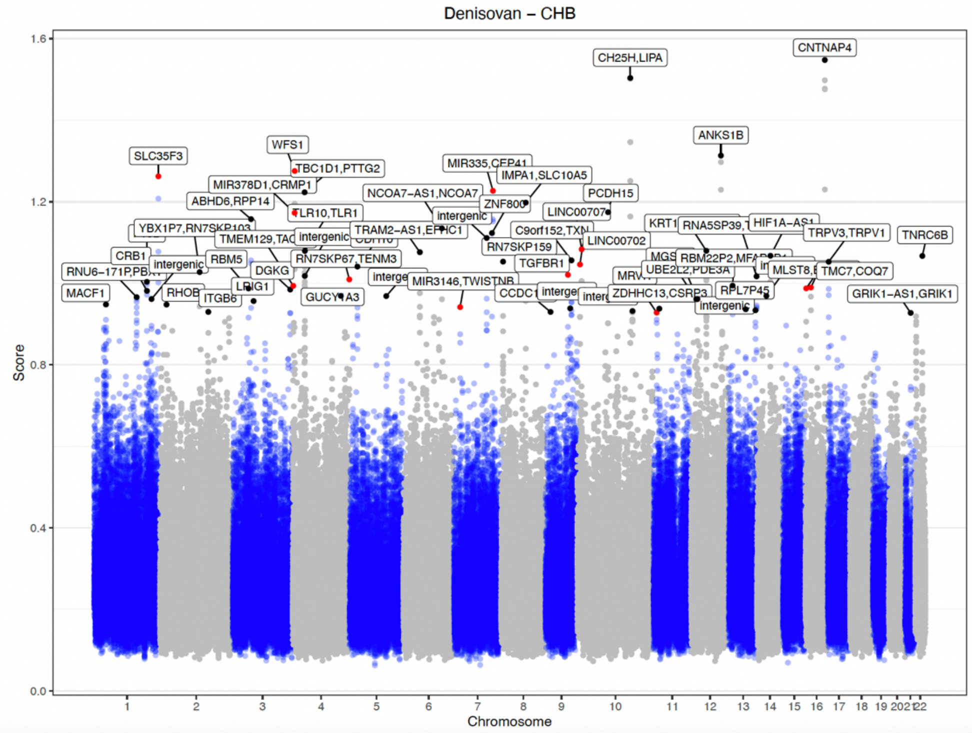
Denisovan – CHB
Score
Chromosome
1.6
1.2
0.8
0.4
0.0
CNTNAP4
CH25H,LIPA
ANKS1B
SLC35F3
WFS1
TBC1D1,PTTG2
MIR378D1,CRMP1
IMPA1,SLC10A5
NCOA7−AS1,NCOA7
ABHD6,RPP14
PCDH15
ZNF800
LINC00707
LR10,TLR1
intergenic
TRAM2−AS1,EFHC1
C9orf152,TXN
TRPV3,TRPV1
TNRC6B
RN7SKP159
LINC00702
CRB1
RBM5
RN7SKP67,TENM3
TGFBR1
TMC7,COQ7
DGKG
MIR3146,TWISTNB
UBE2L2,PDE3A
RHOB
GUCY1A3
ZDHHC13,CSRP3
RPL7P45
MACF1
ITGB6
GRIK1−AS1,GRIK1

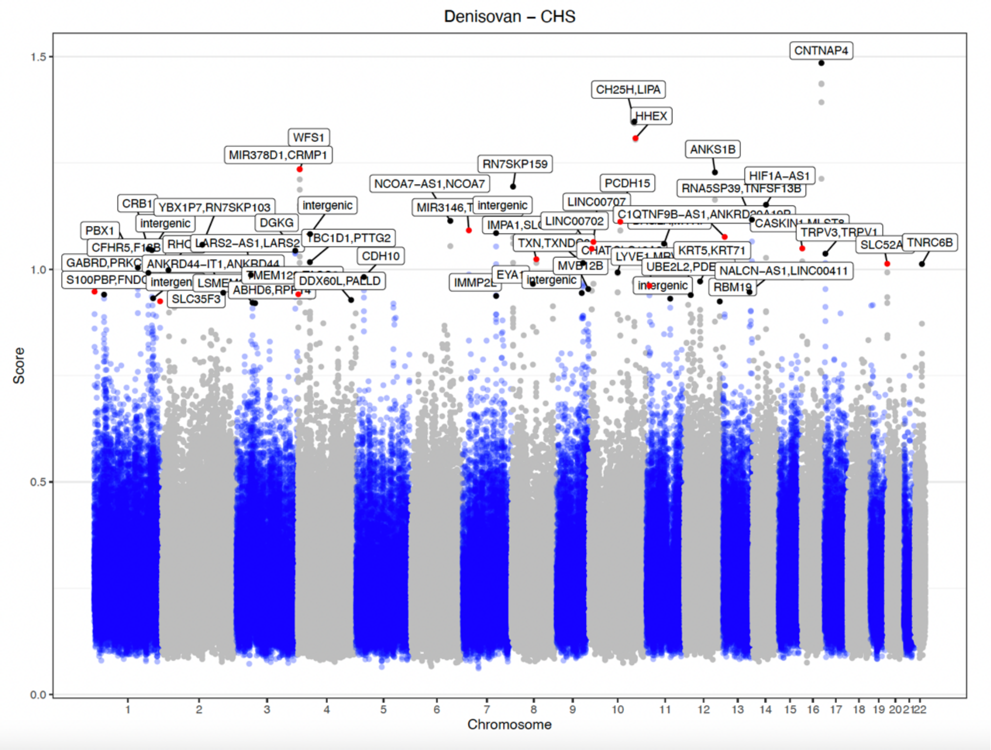
Denisovan – CHS
Score
Chromosome
CNTNAP4
CH25H,LIPA
HHEX
WFS1
MIR378D1,CRMP1
RN7SKP159
ANKS1B
HIF1A–AS1
NCOA7–AS1,NCOA7
PCDH15
RNA5SP39,TNFSF13B
LINC00707
CRB1
YBX1P7,RN7SKP103
intergenic
PBX1
DGKG
TBC1D1,PTTG2
CDH10
TRPV3,TRPV1
TNRC6B
KRT5,KRT71
NALCN–AS1,LINC00411
RBM19
DDX60L,PALLD
SLC35F3
ABHD6
IMMP2L
LINC00702
UBE2L2
MVB12B
S100PBP
GABRD,PRKC
ANKRD44–IT1,ANKRD44
LARS2–AS1,LARS2
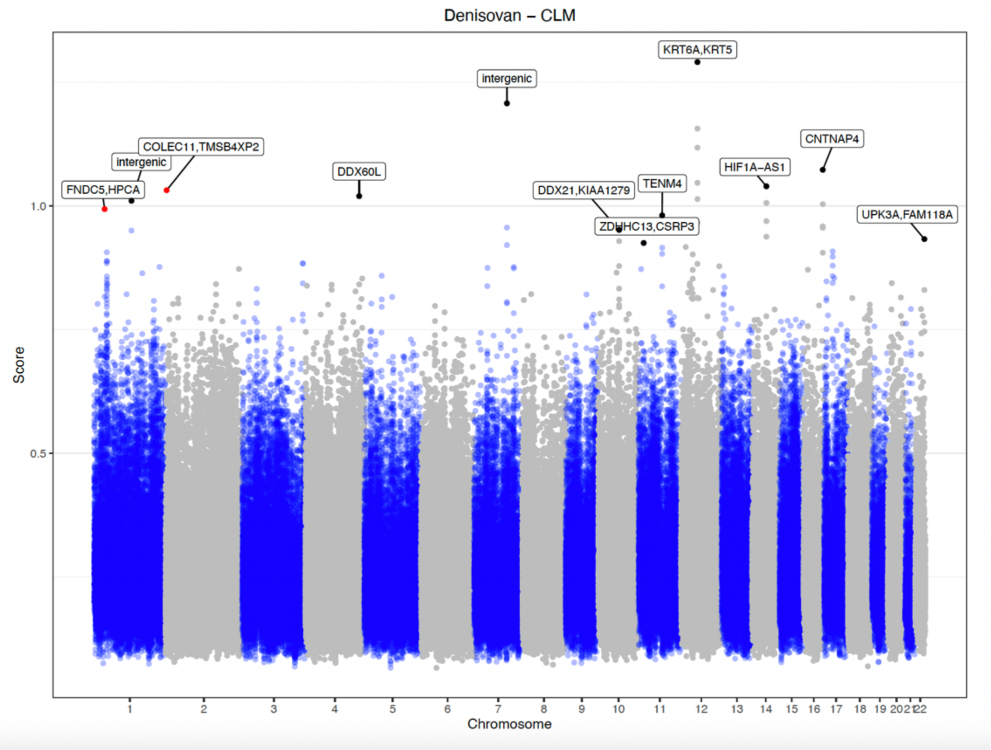
Denisovan – CLM
Score
Chromosome
KRT6A,KRT5
intergenic
COLEC11,TMSB4XP2
FNDC5,HPCA
DDX60L
CNTNAP4
HIF1A–AS1
TENM4
DDX21,KIAA1279
ZDHHC13,CSRP3
UPK3A,FAM118A

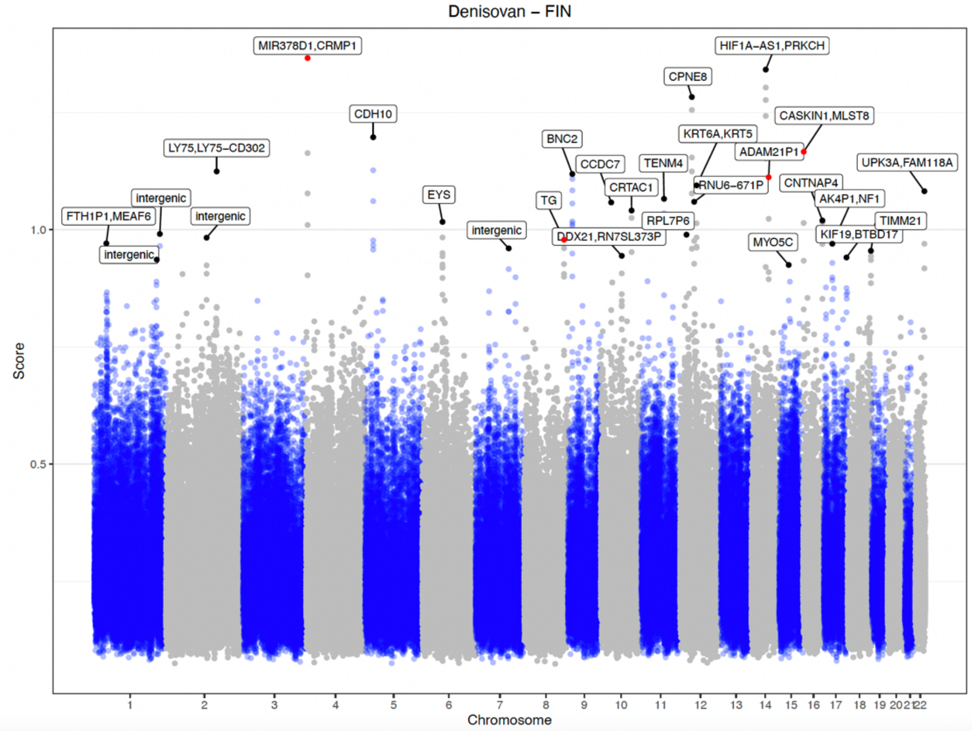
Denisovan – FIN
Score
Chromosome
MIR378D1,CRMP1
HIF1A−AS1,PRKCH
CPNE8
CDH10
CASKIN1,MLST8
KRT6A,KRT5
BNC2
LY75,LY75−CD302
ADAM21P1
CCDC7
TENM4
UPK3A,FAM118A
RNU6−671P
CRTAC1
CNTNAP4
EYS
TG
AK4P1,NF1
intergenic
FTH1P1,MEAF6
RPL7P6
TIMM21
DDX21,RN7SL373P
KIF19,BTBD17
MYO5C

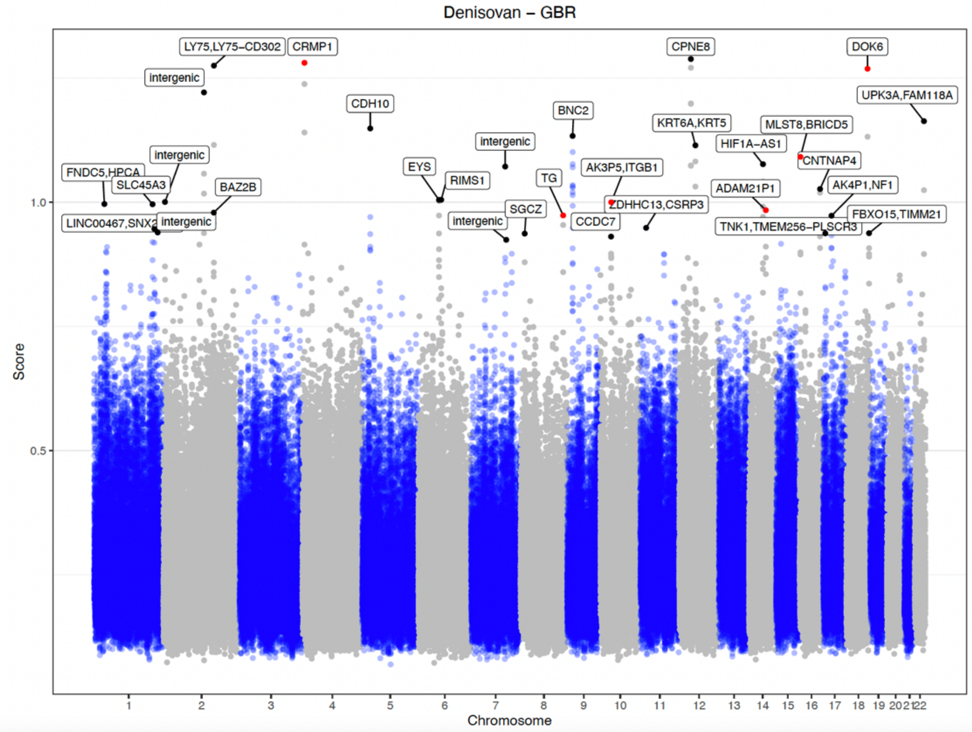
Denisovan – GBR
Score
Chromosome
LY75,LY75−CD302
CRMP1
CPNE8
DOK6
intergenic
UPK3A,FAM118A
CDH10
BNC2
KRT6A,KRT5
MLST8,BRICD5
HIF1A−AS1
FNDC5,HPCA
SLC45A3
BAZ2B
EYS
RIMS1
TG
AK3P5,ITGB1
CNTNAP4
ADAM21P1
AK4P1,NF1
SGCZ
ZDHHC13,CSRP3
LINC00467,SNX2
CCDC7
FBXO15,TIMM21
TNK1,TMEM256−PLSCR3

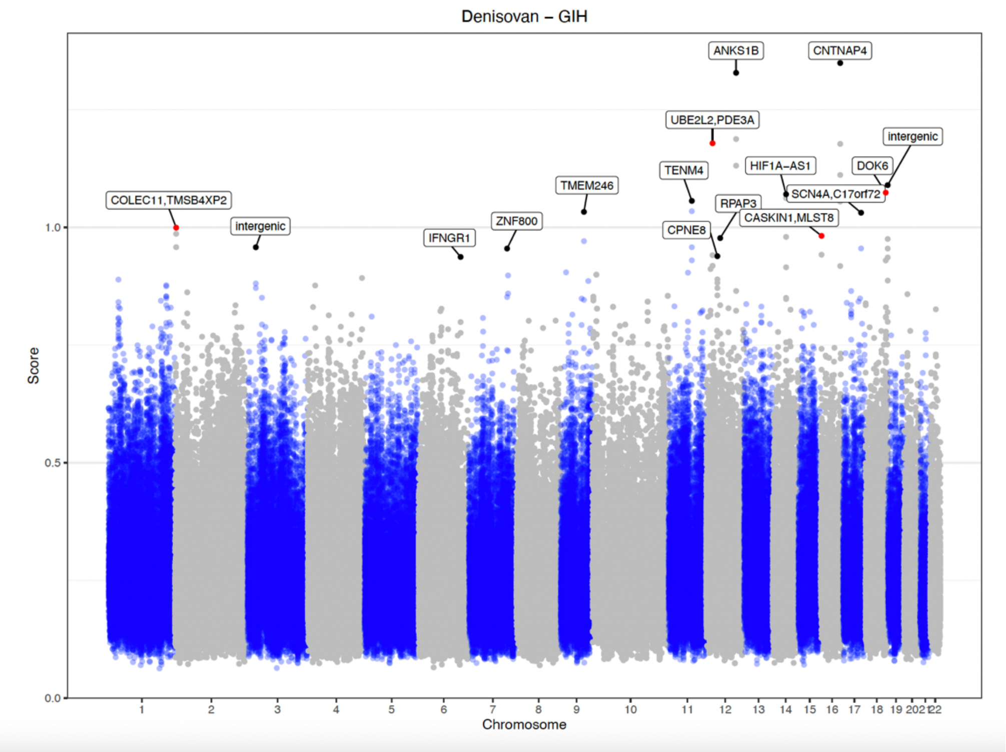
Denisovan – GIH
ANKS1B
CNTNAP4
UBE2L2,PDE3A
intergenic
TENM4
HIF1A–AS1
DOK6
TMEM246
SCN4A,C17orf72
COLEC11,TMSB4XP2
RPAP3
ZNF800
CASKIN1,MLST8
intergenic
CPNE8
IFNGR1
Score
Chromosome
1.0
0.5
0.0

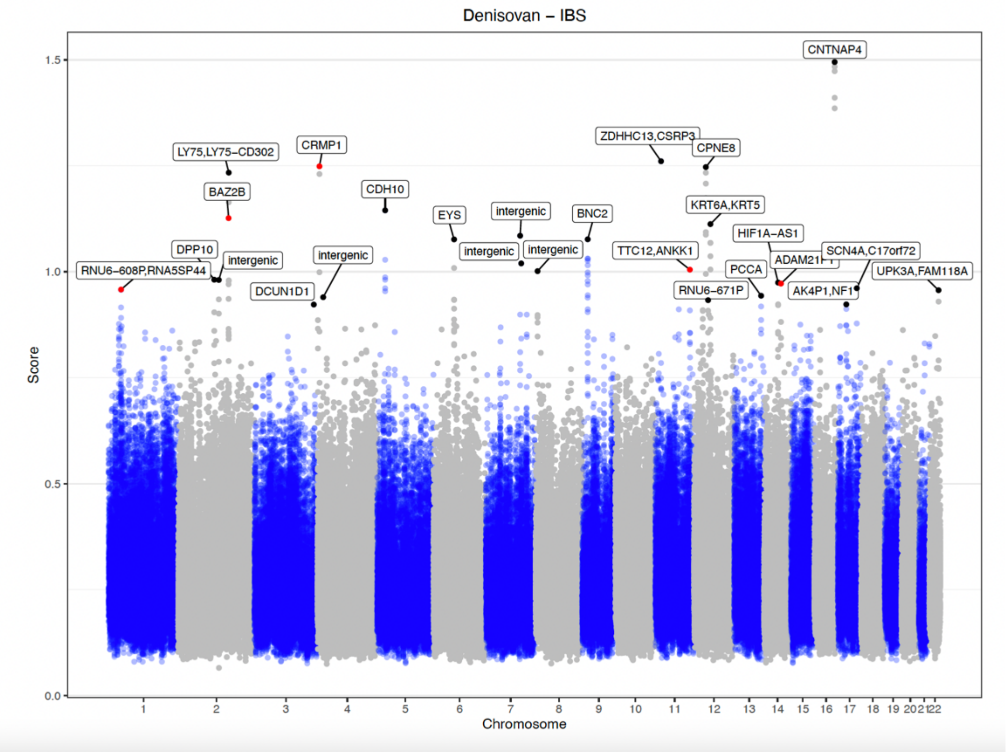
Denisovan – IBS
CNTNAP4
ZDHHC13,CSRP3
LY75,LY75–CD302
CRMP1
CPNE8
BAZ2B
CDH10
KRT6A,KRT5
EYS
intergenic
BNC2
HIF1A–AS1
DPP10
intergenic
intergenic
intergenic
TTC12,ANKK1
SCN4A,C17orf72
RNU6–608P,RNA5SP44
intergenic
PCCA
UPK3A,FAM118A
DCUN1D1
RNU6–671P
AK4P1,NF1
Score
Chromosome
1.5
1.0
0.5
0.0

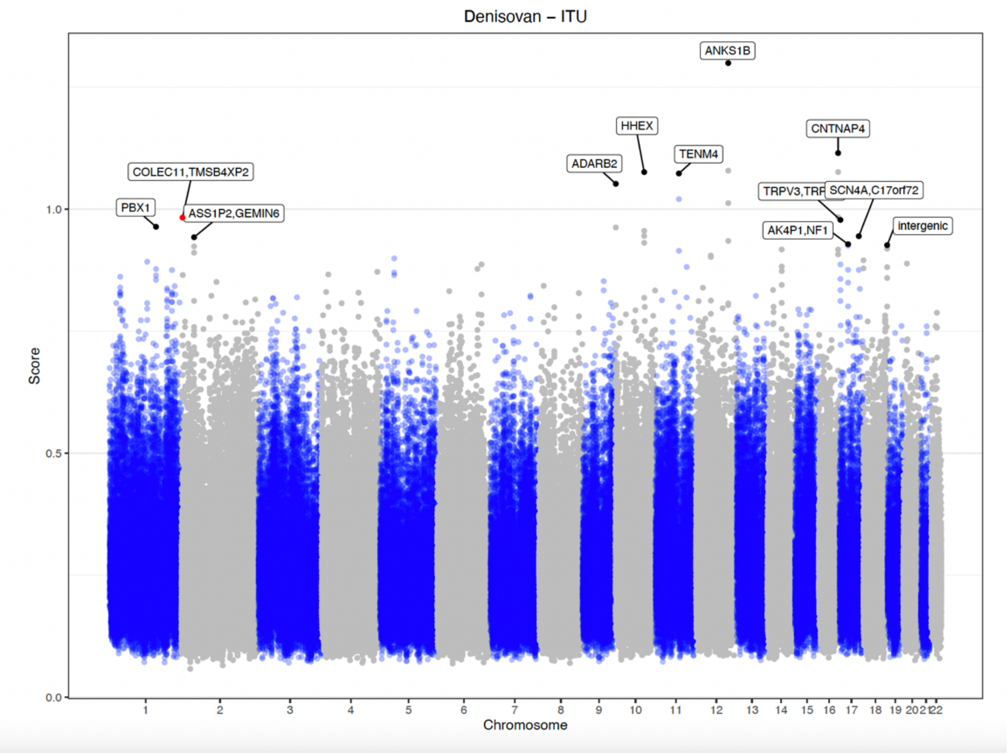
Denisovan – ITU
ANKS1B
HHEX
TENM4
CNTNAP4
ADARB2
COLEC11,TMSB4XP2
PBX1
ASS1P2,GEMIN6
TRPV3,TRP
SCN4A,C17orf72
AK4P1,NF1
intergenic
Score
1.0
0.5
0.0
Chromosome

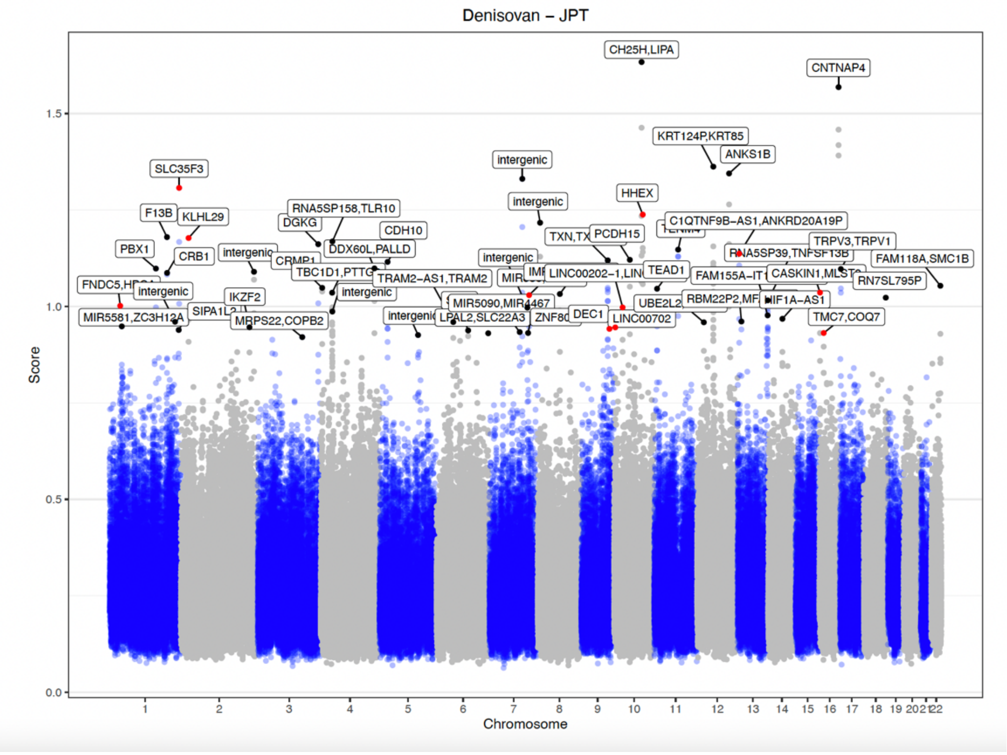
Denisovan – JPT
CH25H,LIPA
CNTNAP4
KRT124P,KRT85
ANKS1B
intergenic
SLC35F3
HHEX
F13B
KLHL29
RNA5SP158,TLR10
DGKG
CDH10
C1QTNF9B-AS1,ANKRD20A19P
TXN,TX
PCDH15
PBX1
CRB1
intergenic
DDX60L,PALLD
TRPV3,TRPV1
RNA5SP39,TNFSF13B
FAM118A,SMC1B
CRMP1
TBC1D1,PTTG
TRAM2-AS1,TRAM2
TEAD1
FAM155A-IT1
CASKIN1,MLST
RN7SL795P
FNDC5,HP
intergenic
IKZF2
LINC00202-1,LIN
MIR5090,MIR4467
UBE2L2
RBM22P2,MF
HIF1A-AS1
MIR5581,ZC3H12A
SIPA1L
MRPS22,COPB2
LPAL2,SLC22A3
ZNF8
DEC1
LINC00702
TMC7,COQ7
Score
1.5
1.0
0.5
0.0
Chromosome

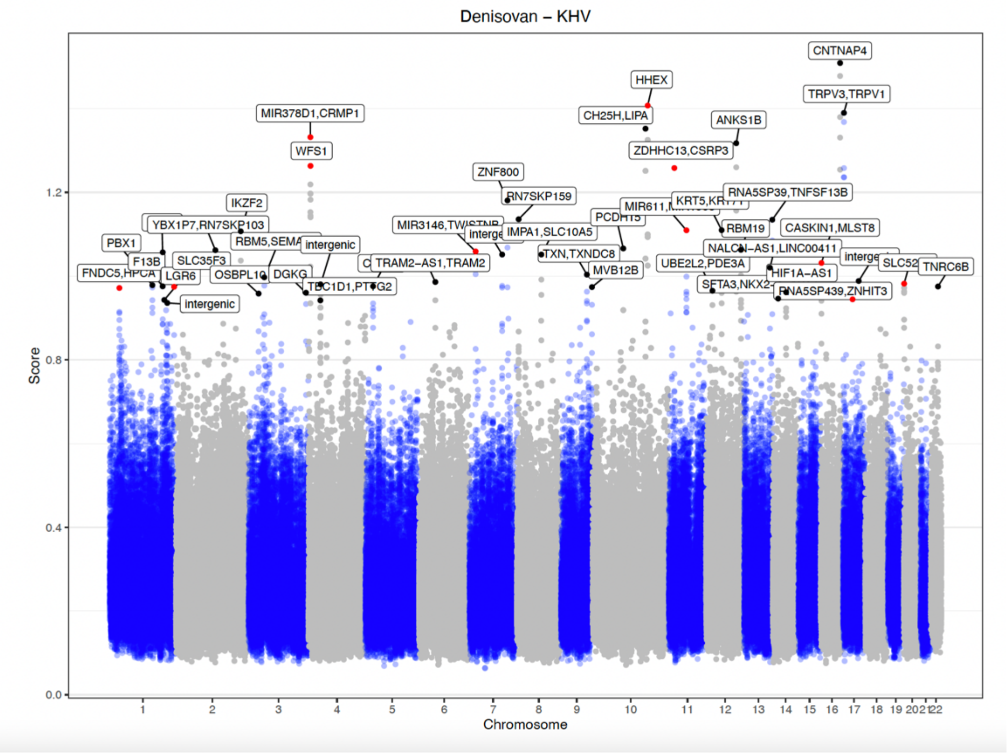
Denisovan – KHV
Score
Chromosome
CNTNAP4
HHEX
TRPV3,TRPV1
MIR378D1,CRMP1
CH25H,LIPA
ANKS1B
WFS1
ZDHHC13,CSRP3
ZNF800
RN7SKP159
IKZF2
RNA5SP39,TNFSF13B
YBX1P7,RN7SKP103
MIR3146
IMPA1,SLC10A5
RBM19
CASKIN1,MLST8
PBX1
intergenic
F13B
SLC35F3
TXN,TXNDC8
FNDC5,HPCA
LGR6
OSBPL10
DGKG
TRAM2–AS1,TRAM2
MVB12B
UBE2L2,PDE3A
HIF1A–AS1
TNRC6B
TBC1D1,PTTG2
RNA5SP439,ZNHIT3

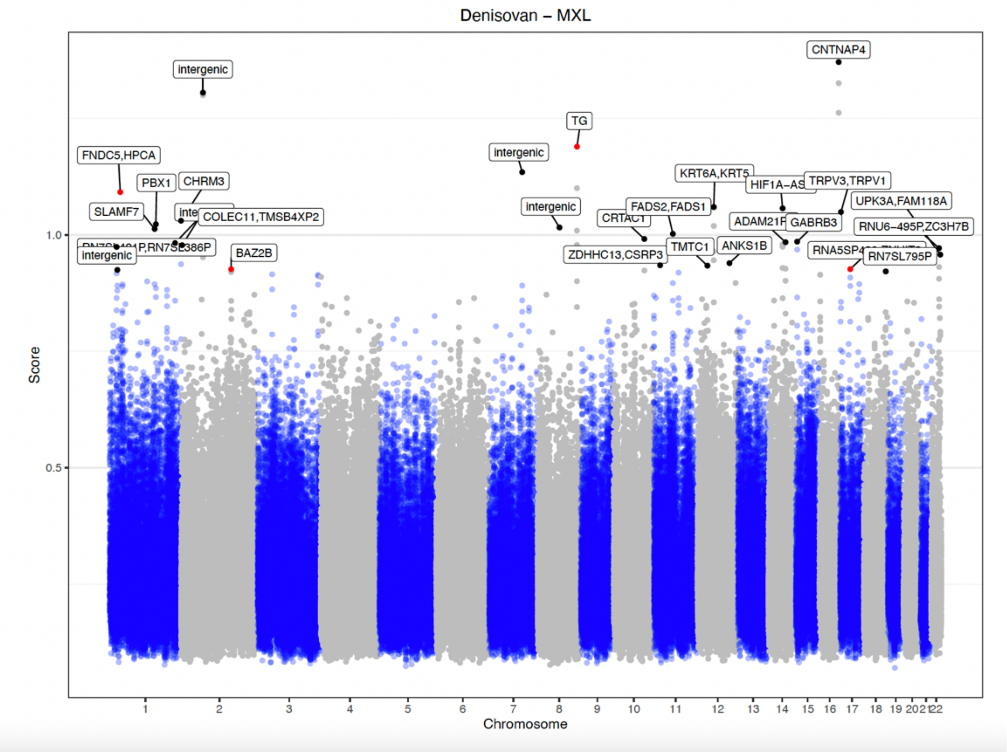
Denisovan – MXL
Score
Chromosome
CNTNAP4
intergenic
TG
FNDC5,HPCA
PBX1
CHRM3
KRT6A,KRT5
TRPV3,TRPV1
UPK3A,FAM118A
SLAMF7
COLEC11,TMSB4XP2
FADS2,FADS1
ADAM21P
GABRB3
RNU6–495P,ZC3H7B
BAZ2B
ZDHHC13,CSRP3
TMTC1
ANKS1B
RN7SL795P

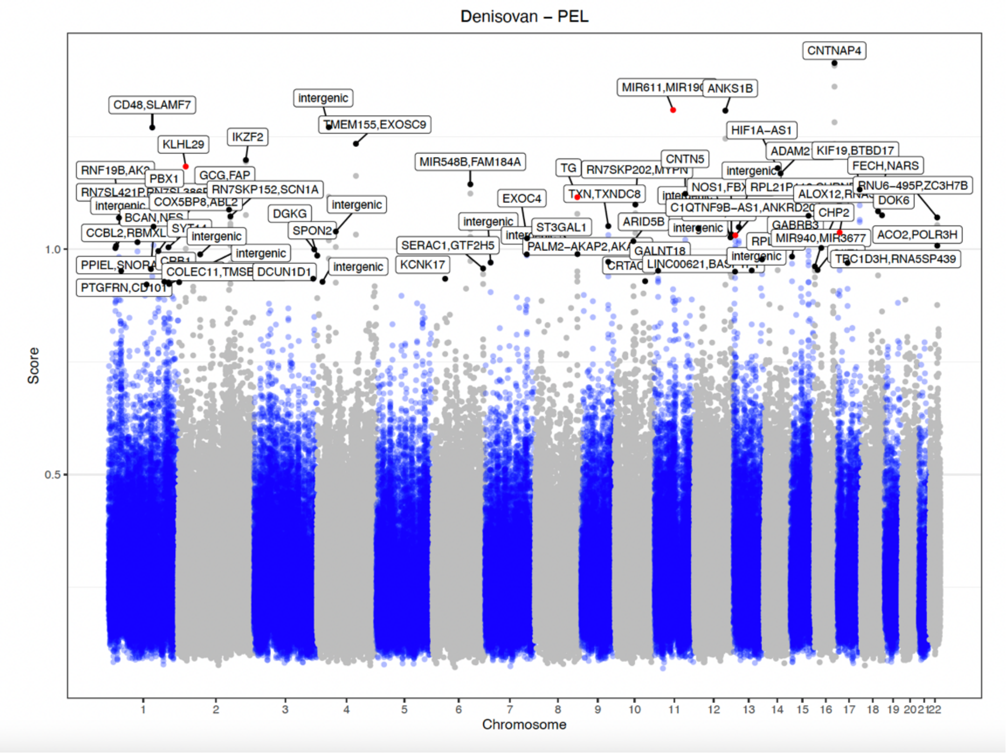
Denisovan – PEL
Score
Chromosome
0.5
1.0
CNTNAP4
ANKS1B
CD48,SLAMF7
intergenic
TMEM155,EXOSC9
KLHL29
IKZF2
HIF1A–AS1
MIR548B,FAM184A
CNTN5
KIF19,BTBD17
FECH,NARS
RNU6–495P,ZC3H7B
EXOC4
ST3GAL1
SERAC1,GTF2H5
KCNK17
DGKG
SPON2
GCG,FAP
RN7SKP152,SCN1A
PBX1
PPIEL,SNORA
PTGFRN,CD101
DCUN1D1
ARID5B
GALNT18
DOK6
CHP2
ACO2,POLR3H
TBC1D3H,RNA5SP439
MIR940,MIR3677
TXN,TXNDC8
TG
C1QTNF9B–AS1,ANKRD20

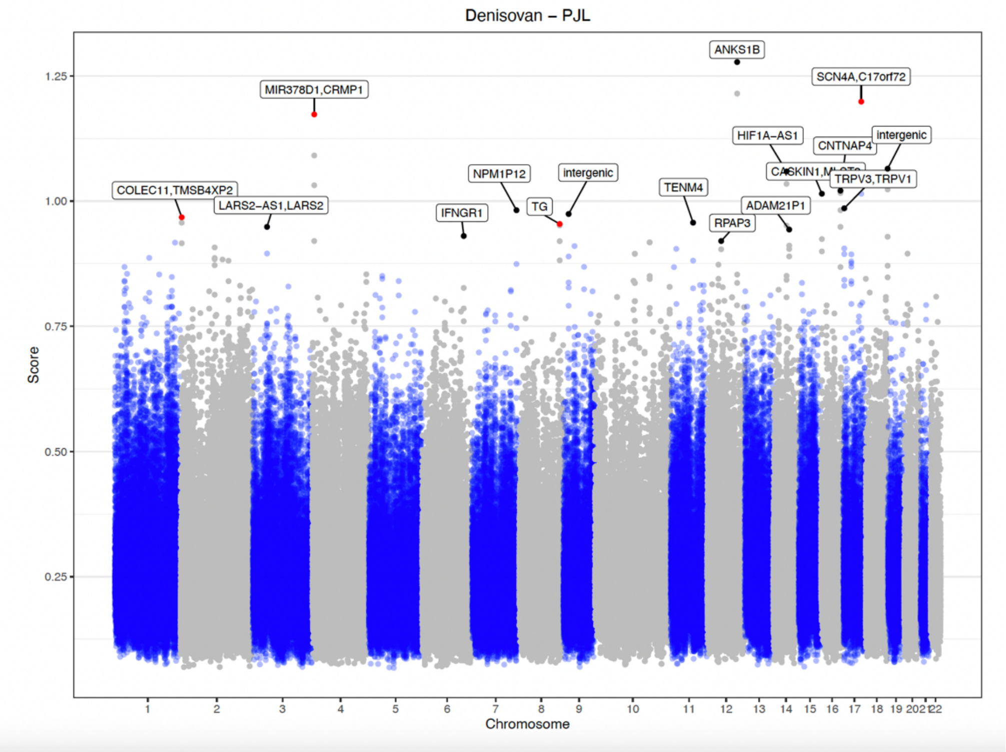
Denisovan – PJL
Score
Chromosome
0.25
0.50
0.75
1.00
1.25
ANKS1B
SCN4A,C17orf72
MIR378D1,CRMP1
HIF1A–AS1
intergenic
CNTNAP4
TRPV3,TRPV1
NPM1P12
intergenic
COLEC11,TMSB4XP2
LARS2–AS1,LARS2
IFNGR1
TG
TENM4
ADAM21P1
RPAP3

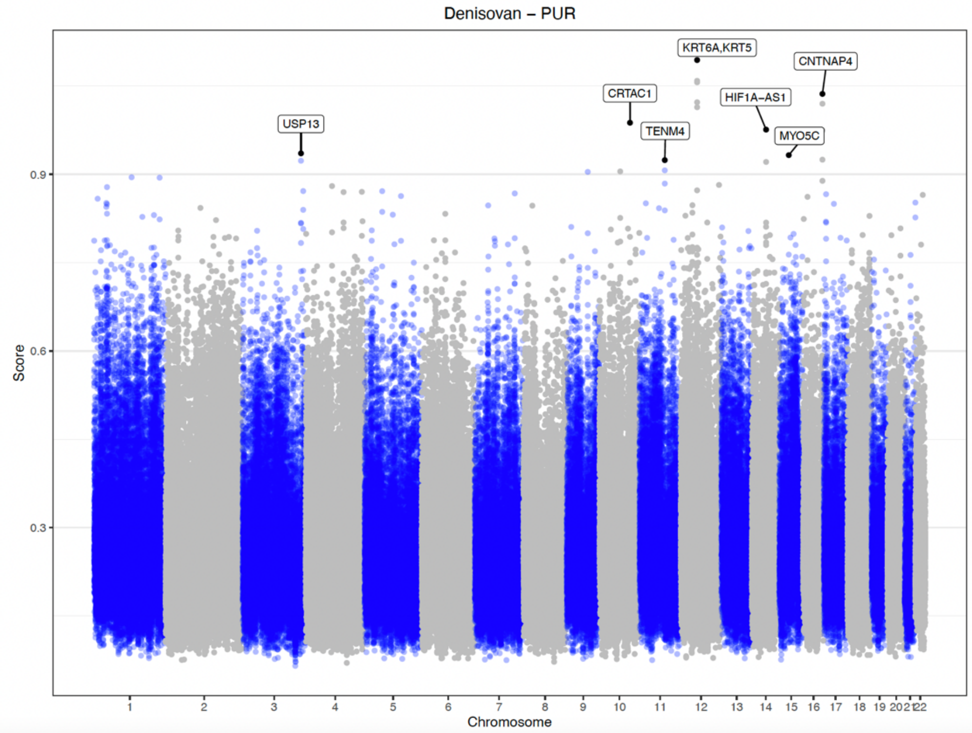
Denisovan – PUR
KRT6A,KRT5
CNTNAP4
CRTAC1
HIF1A–AS1
USP13
TENM4
MYO5C
Score
0.9
0.6
0.3
Chromosome

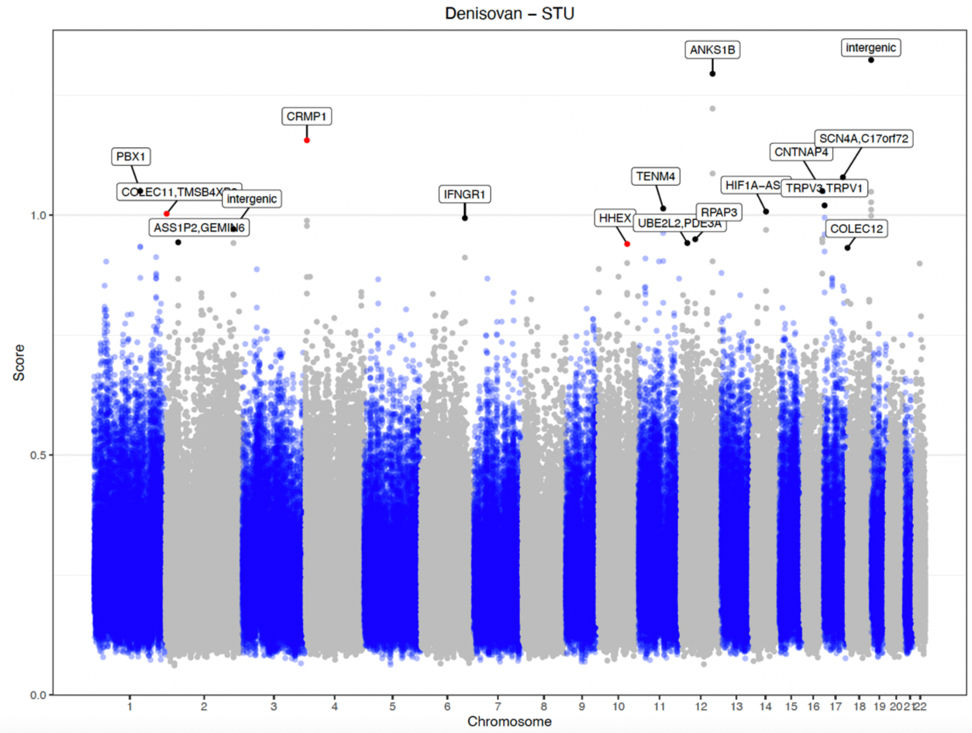
Denisovan – STU
ANKS1B
intergenic
CRMP1
SCN4A,C17orf72
CNTNAP4
PBX1
TENM4
HIF1A–AS
TRPV3,TRPV1
intergenic
IFNGR1
RPAP3
HHEX
COLEC12
Score
1.0
0.5
0.0
Chromosome

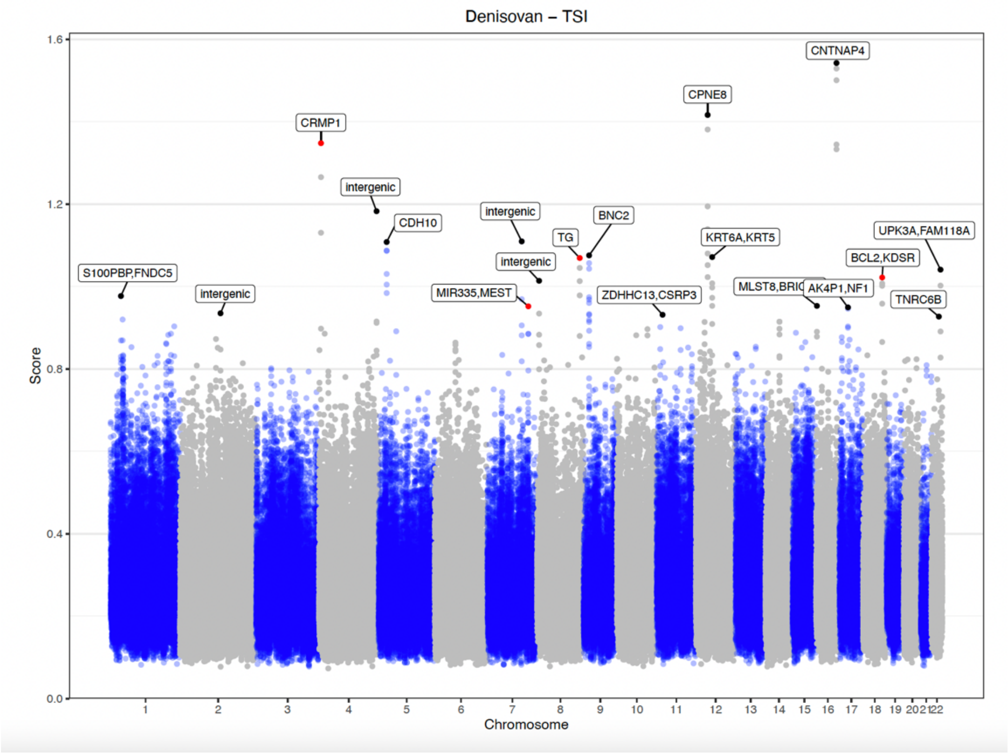

*We applied MaLAdapt to predict AI in overlapping 50kb windows (step size 10kb) along the genome of non-African populations of the 1000 Genomes data. Here we show the AI prediction results of 19 non-African populations, using African (YRI) as non-introgressed outgroup and Altai Denisovan as the introgression donor. The Y-axis shows the AI prediction score, which equals the -Log10 transformed value of [1-Pr(AI)]. Each dot in the plot represents a 50kb window. The windows that did not reach the MaLAdapt AI threshold are colored in blue or gray depending on the chromosomes. The windows detected as AI are colored in black if they have been reported by previous studies before, or in red if they are novel findings from this study. The labels highlight the gene names that overlap with the AI windows.*

**Supplementary Figure 18: Haplostrips of Neanderthal AI hits in CEU predicted by *genotmatnn* that are visually inspected as False Positives**

a) CHR1:104500001-104600000

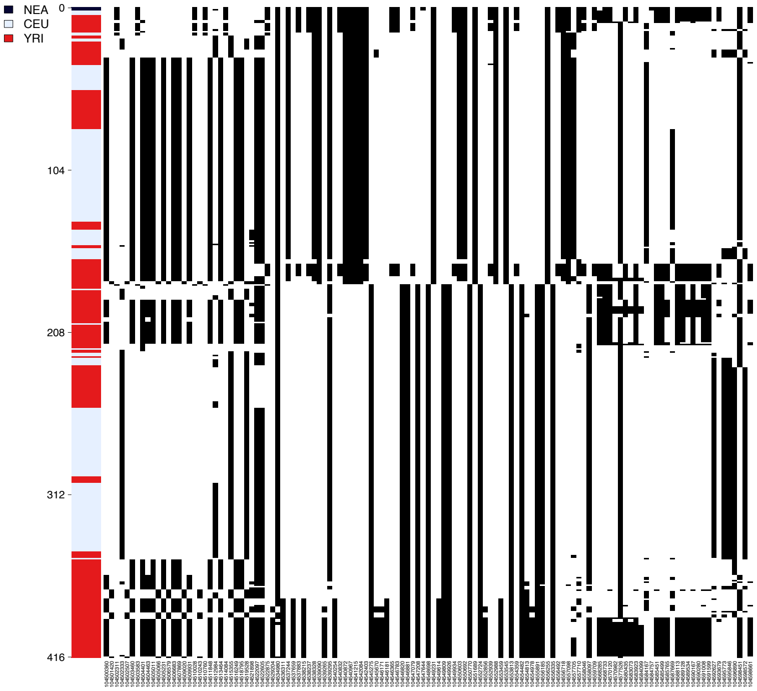

b) CHR2:227800001-227900000

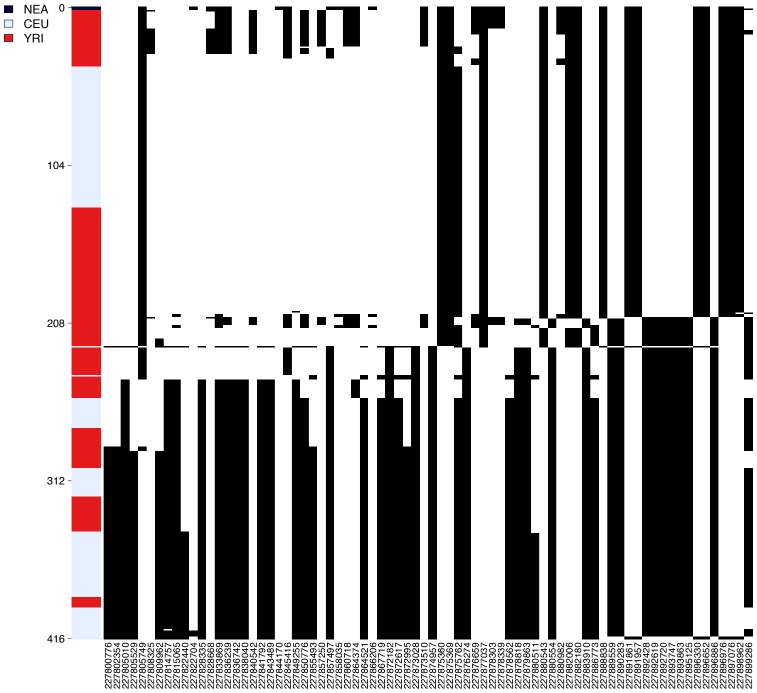

c) CHR5:39220001-39320000

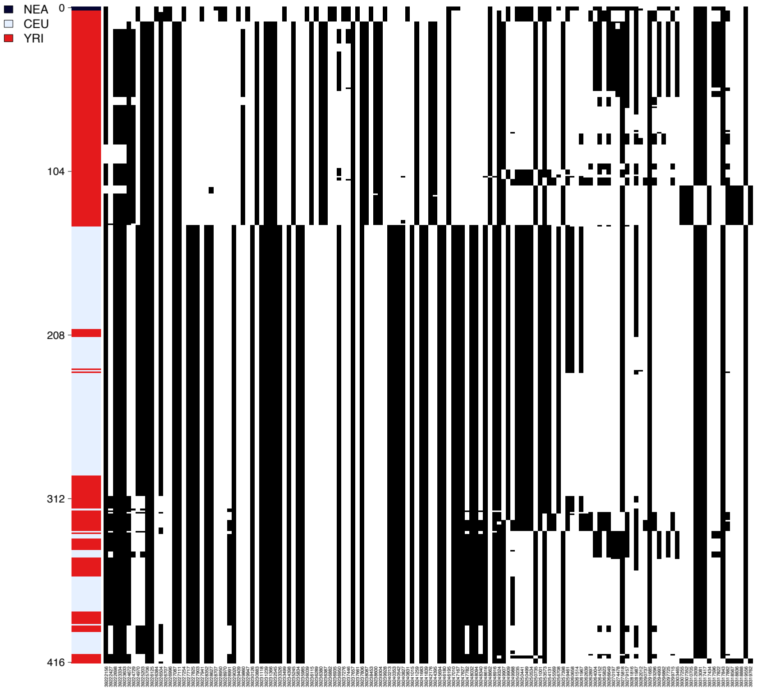

d) CHR8:91840001-91940000

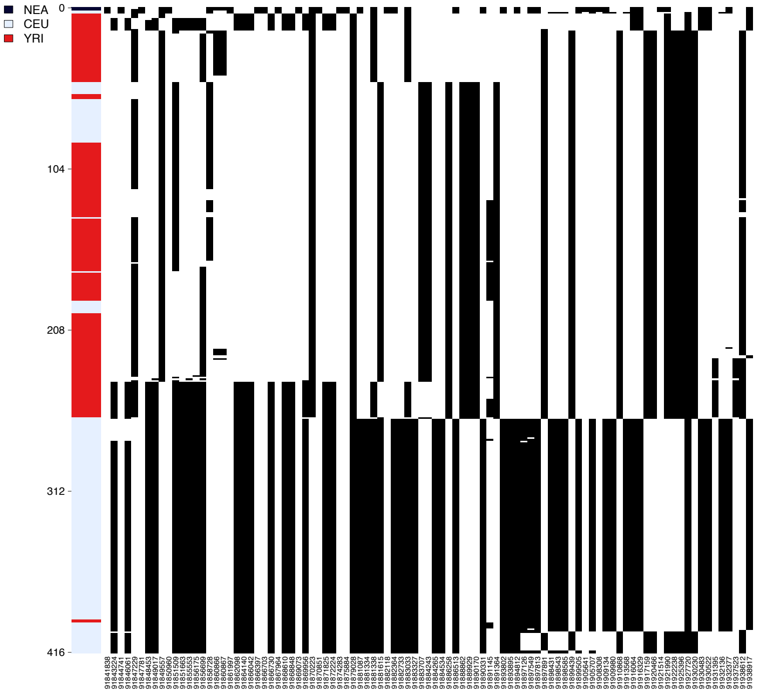

e) CHR8:143440001-143560000

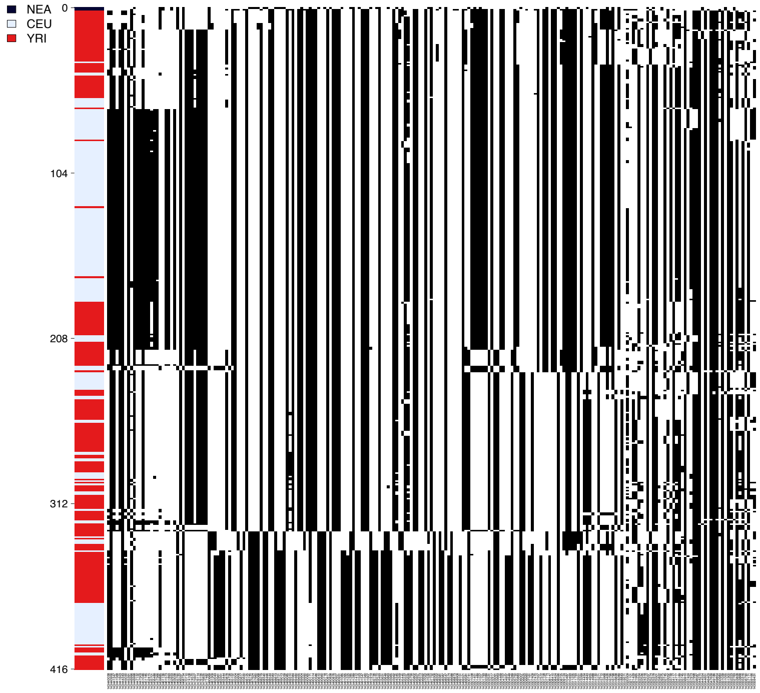

f) CHR19:20220001-20380000

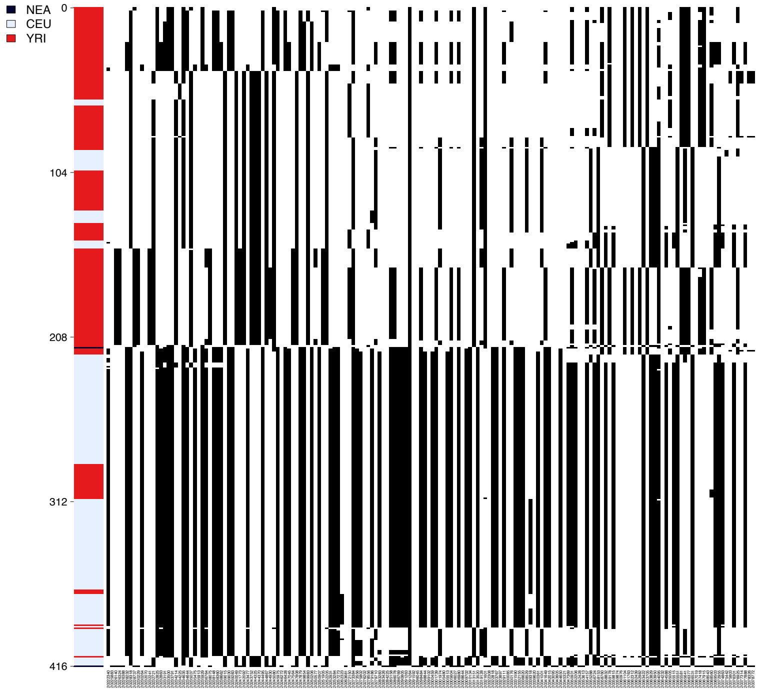

g) CHR19:20260001-20360000

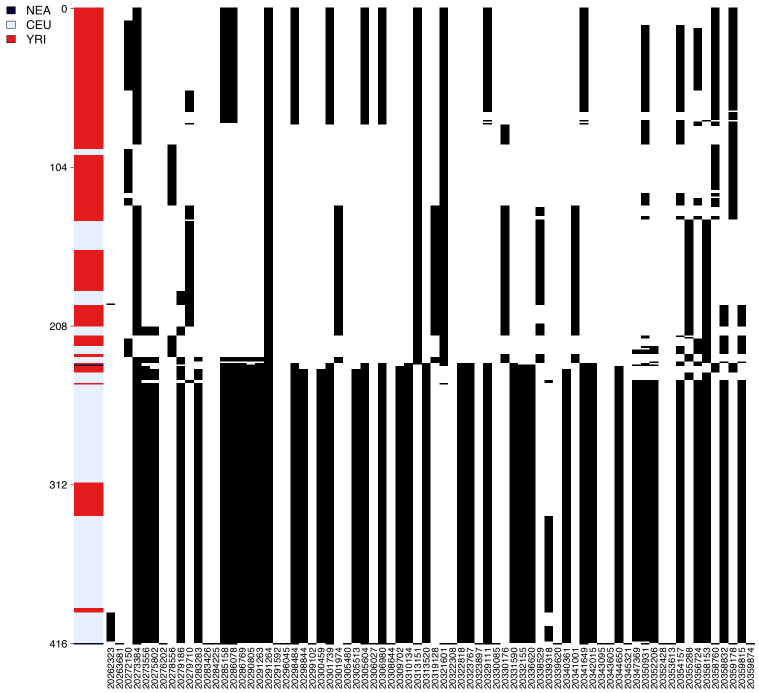

*We show the haplotype structures of 6 Neanderthal AI regions in CEU predicted by genomatnn that we believe are likely false positives. For each region, we plotted the haplotypes of Altai Neanderthal (black), CEU individuals (blue) and YRI individuals (red), and clustered and sorted the haplotypes by decreasing distance to the Neanderthal genome. Rows closer to the top of the plot represent haplotypes are more similar to that of the Neanderthal. In the haplotype structure, each row represents a haplotype, and the column denotes a SNP (black lines indicate the presence of alternative allele).*
